# SPIDER: Contextualizing Biomedical Knowledge for Evidence-Grounded Scientific Reasoning

**DOI:** 10.64898/2026.11.11.765839

**Authors:** Mingfei Liu, Weike Zhao, Jingyi Chen, Xingran Quan, Xinyi Wang, Jinghao Feng, Xiao Zhou, Jiaxuan Han, Ya Zhang, Yanfeng Wang, Xiaoman Zhang, Weidi Xie

## Abstract

Aggregated relations in biomedical knowledge graphs can obscure evidence and study conditions, complicating the distinction between scientific disagreement and contextual differences. Here we introduce **SPIDER**, a multi-agent framework grounding biomedical relations in literature evidence and extracting study conditions for interpretation. Applied to a graph integrating 48 databases and ontologies, SPIDER analyzed 1.95 million PubMed abstracts, adding 2.31 million previously unrepresented, literature-supported entity pairs to construct **SPIDER-KG**. Daily updates incorporate relations and evidence from newly indexed PubMed records. Across 89,937 evidence-supported comparisons of potentially incompatible relations, SPIDER classified 79.0% as contextual differences, and 6.1% as genuine disagreement, achieving 80.3% macro-F1 averaged across both categories against expert annotations. On SPIDER-Bench and four biomedical reasoning benchmarks, SPIDER-KG improved large language models’ interpretation of evidence-supported biomedical relationships. Retrospective analysis of newly reported gene–disease relationships further demonstrated how evidence-linked knowledge can contextualize emerging discoveries. SPIDER-KG provides a reusable, incrementally extensible evidence layer for biomedical reasoning.

## INTRODUCTION

Biomedical discovery requires determining how findings from different studies can be combined to support an explanation or a testable hypothesis. A proposed disease mechanism or therapeutic intervention may connect observations across molecular, cellular, and clinical scales^1,2^. Yet the relevance of these observations depends on the biological systems, experimental designs and treatment settings in which they were obtained^3,4^. Computational systems that assist discovery therefore need access not only to reported findings, but also to the evidence supporting them and the conditions under which they apply.

Biomedical knowledge graphs (KGs) integrate knowledge from databases, ontologies and the literature^2,5–10^, supporting target identification, drug repurposing and reasoning with large language models^11–19^. Several resources also link relations to publications, supporting passages or curated records^20,21^, or include temporal or contextual information^22,23^. However, a single publication may report several observations, and the same relation may be supported by studies involving different tissues, experimental models or interventions. Such provenance links and contextual information do not necessarily associate each individual finding with its supporting evidence and the specific study conditions.

**Figure 1.**
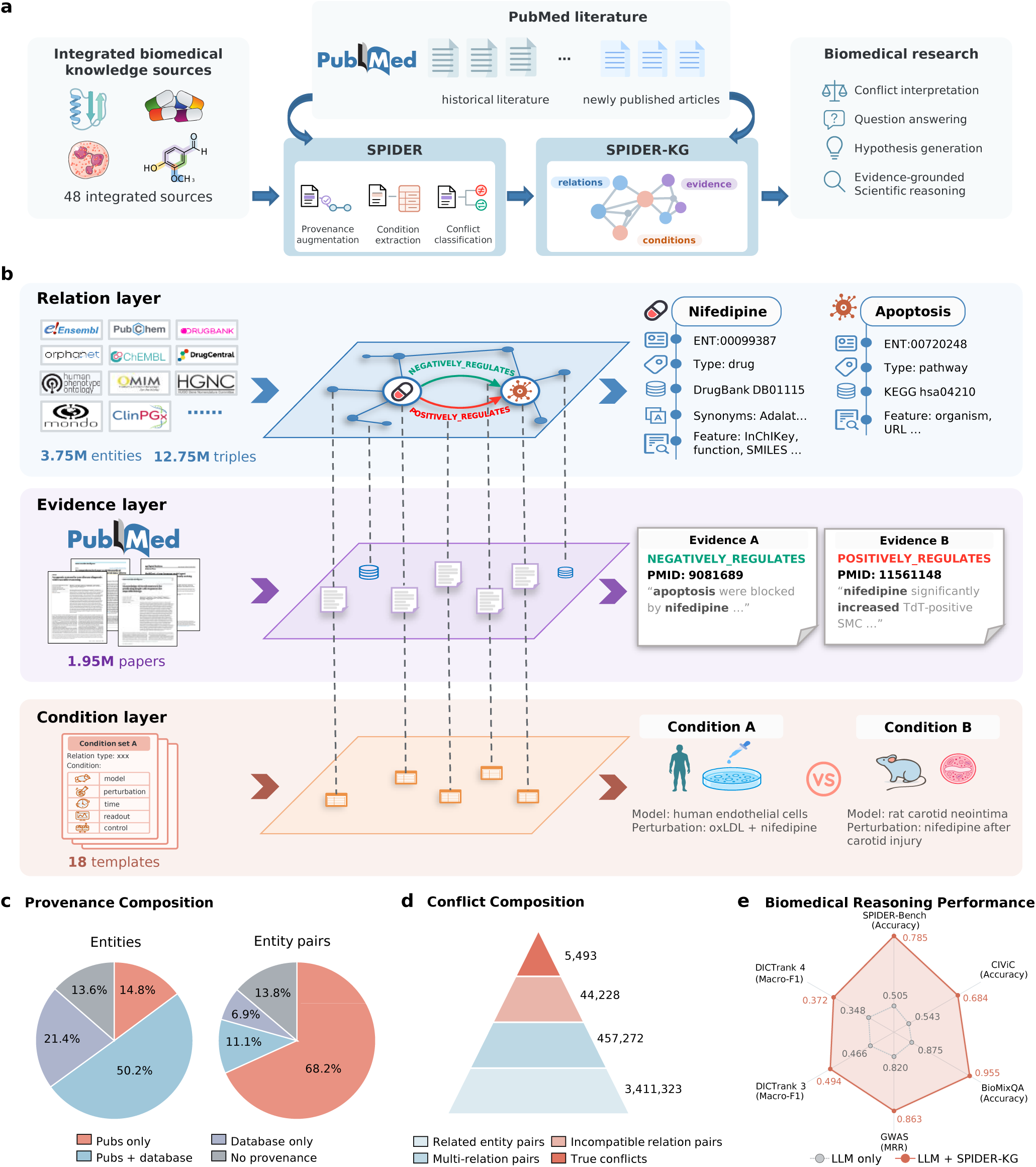
Overview and evaluation of SPIDER and SPIDER-KG. **a**, SPIDER integrates 48 biomedical databases and ontologies with PubMed literature to contextualize biomedical knowledge by linking graph relations to literature evidence and study conditions, constructing and incrementally updating SPIDER-KG. **b,** Organization of SPIDER-KG into relation, evidence and condition layers, illustrated by context-dependent findings. **c,** Provenance composition of entities and entity pairs after literature grounding. Pubs, supporting publications. **d,** Counts of multi-relation entity pairs and potentially incompatible relation comparisons, with the number of comparisons classified as true conflicts. **e,** Comparison of LLM-only and SPIDER-KG-augmented methods on SPIDER-Bench and four biomedical reasoning benchmarks.

One consequence of losing this information is the difficulty of interpreting apparently conflicting biomedical findings. A drug may inhibit a biological process in one cell type and promote it in another^3,4^. Both findings may be valid, although they appear as opposing relations between the same entities. Preserving each finding together with its evidence and reported study conditions would allow a KG to represent this variation and help researchers or language models distinguish context-dependent effects from genuine disagreement under comparable conditions.

Here, we introduce **SPIDER**, a multi-agent framework that grounds biomedical knowledge graph relations in literature evidence and captures the study conditions associated with each finding (Fig. 1). Starting from a research-oriented knowledge graph integrating 48 databases and ontologies, we analyze titles and abstracts from nearly 2 million PubMed articles to construct **SPIDER-KG**. This process adds 2.31 million previously unrepresented entity pairs together with their supporting literature evidence. SPIDER also supports incremental updates as new PubMed articles become available.

**SPIDER** achieves strong entity and relation extraction performance on the literature extraction benchmark. Across 89,937 evidence-supported comparisons of potentially incompatible relations, 79.0% are classified as context-dependent differences and 6.1% as genuine disagreement under comparable conditions; this classification achieves a macro-F1 of 80.3% averaged over the two categories against expert annotations. To assess the utility of contextualized evidence for reasoning, we introduce **SPIDER-Bench**, a conflict-aware question-answering benchmark based on findings that differ across biological and experimental contexts. SPIDER-KG notably improves performance on SPIDER-Bench and four additional biomedical reasoning benchmarks spanning biomedical knowledge judgment, candidate effector-gene ranking, oncology evidence assessment and drug safety classification. Retrospective analysis of newly reported gene–disease relationships further illustrates how SPIDER-KG connects emerging findings with relevant prior knowledge. These evaluations demonstrate the usefulness of evidence and study context for practical uses in biomedical assessment, prioritization and hypothesis development.

## RESULTS

### SPIDER-KG contextualizes biomedical relations with evidence

We constructed SPIDER-KG to preserve biomedical findings with their supporting evidence and reported study conditions. The graph comprises three layers (Fig. 1b): a relation layer, representing each biomedical finding as two entities connected by a relation type; an evidence layer, linking entities and triples to their source database records or publications; and a condition layer, recording the biological or experimental conditions reported for each finding. A triple may be supported by multiple database records or publications. The same entity pair may also be connected by multiple triples with potentially incompatible relation types, such as positive and negative regulation. For these triples, we retained findings from different supporting publications separately, together with their reported study conditions.

We integrated 48 databases and ontologies into an initial graph containing 3.75 million entities and 12.75 million triples (Fig. 2). However, as many concepts in this graph serve primarily taxonomic or hierarchical classification and are not typically reported as independent biomedical findings, we selected a research-oriented subgraph using predefined three-stage criteria (Appendix S2.5). This subgraph comprises 198,659 entities and 1,099,652 entity pairs. We grounded its relations in literature evidence and extracted structured study conditions to support comparisons of potentially incompatible relations (Methods, “Identification of potentially incompatible relations”). SPIDER-KG is updated incrementally with evidence and relations extracted from newly added PubMed records.

**Figure 2.**
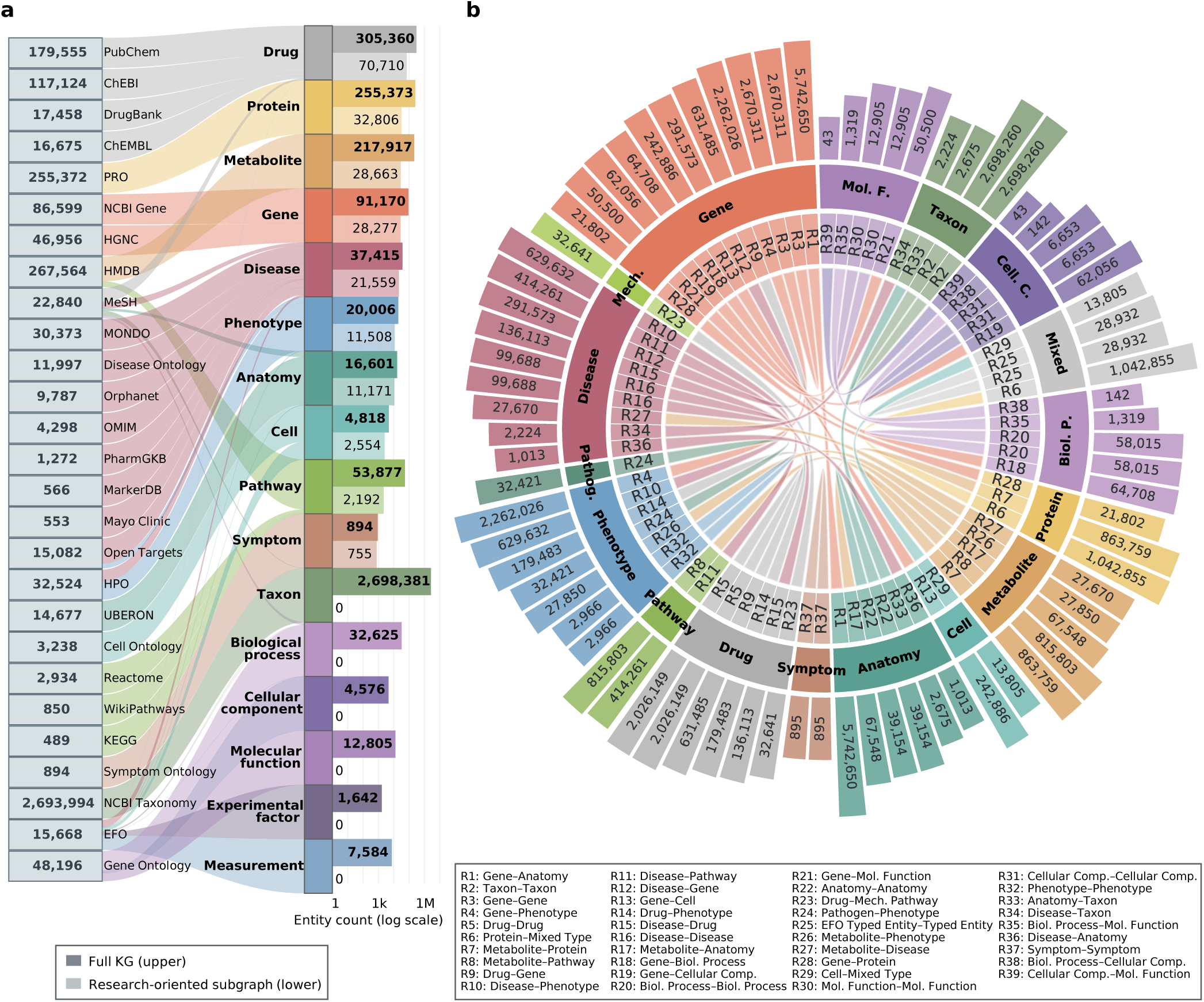
Composition of the database-derived foundation of SPIDER-KG. **a**, Source contributions and entity counts across 16 standardized entity types in the database-derived KG. Bars show entity counts in the full KG and the research-oriented subgraph used for literature analysis. **b,** Distribution of source-level relation records across 39 endpoint-type combinations. Outer bars indicate triple abundance and colors denote entity types. R1–R39 correspond to the relation groups listed below the diagram.

### SPIDER grounds biomedical relations through literature extraction

We first evaluated the proposed SPIDER framework on biomedical entity and relation extraction using a benchmark of 1,000 PubMed abstracts. We tested the framework with seven LLM backbones and compared it with three conventional methods: BERN2^24^, PubTator^25^ and GLiNER-Relex^26^. We also included a DeepSeek-V4-Flash^27^ baseline that extracted both entities and relations in a single LLM call. Compatibility-based scoring matches triples within an abstract by aligned entity pairs, allowing different relation labels subject to predefined directionality and incompatibility rules (Appendix S2.9.3).

**Figure 3.**
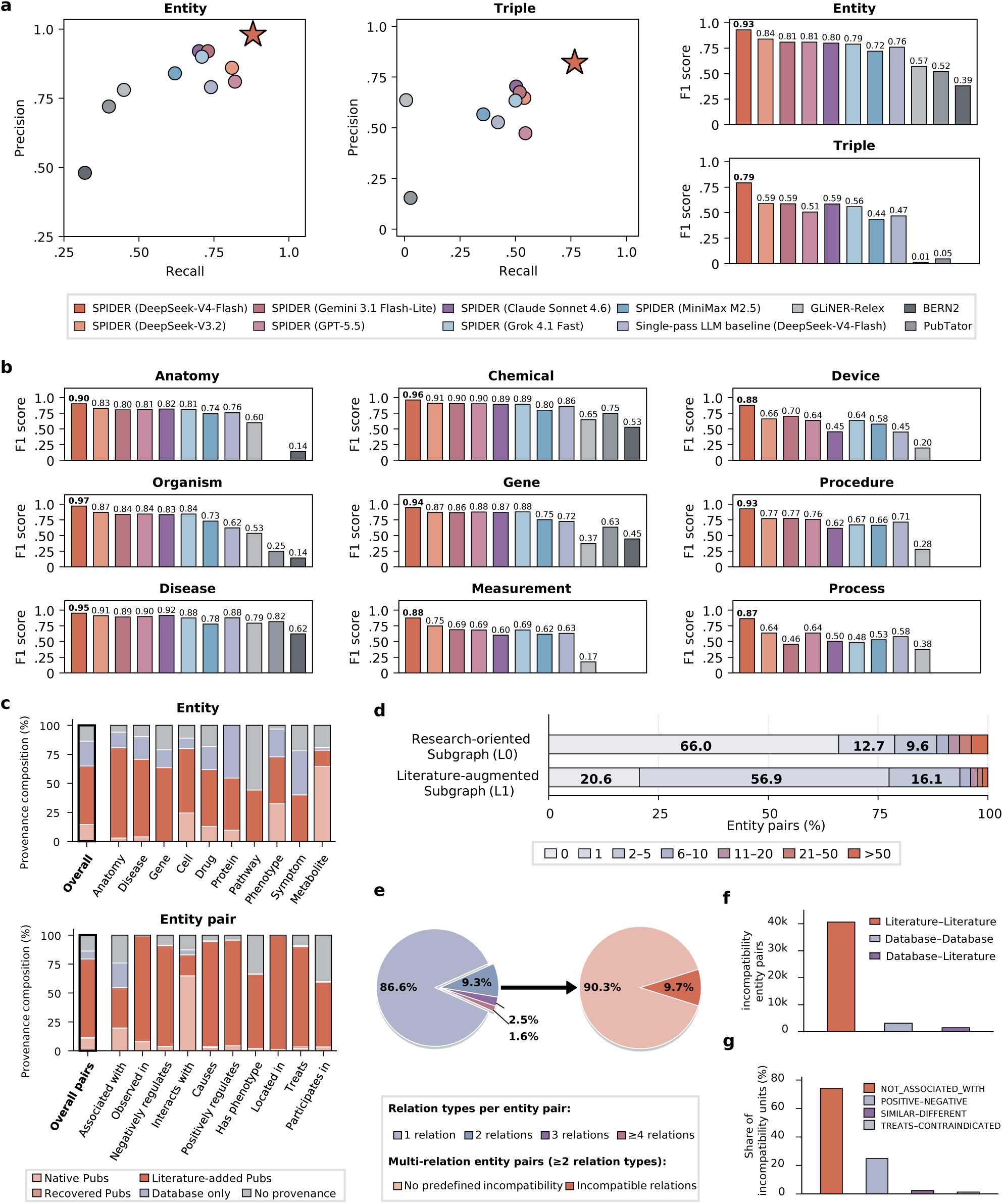
Extraction performance, publication coverage and potentially incompatible relations. **a**, Entity-extraction and compatibility-based triple-matching performance of SPIDER and baseline methods on the literature extraction benchmark. **b,** Entity-extraction F1 across 9 extraction entity categories. Missing bars indicate entity categories not supported by a method. **c,** Provenance composition of entity and entity pair after literature augmentation. Pubs, supporting publications. **d,** Distribution of linked publications per entity pair in the research-oriented subgraph before and after literature augmentation. **e,** Relation multiplicity and prevalence of predefined incompatibility patterns among *L*_1_ entity pairs. **f,** Provenance composition of potentially incompatible entity pairs. **g,** Distribution of predefined incompatibility patterns among comparison units.

SPIDER with DeepSeek-V4-Flash achieved the highest overall precision, recall and F1 for both entity extraction and compatibility-based triple matching among the evaluated methods, with F1 scores of 0.928 and 0.793, respectively (Fig. 3a). It also led on all three metrics in each of the 9 entity categories (Fig. 3b, Supplementary Table S21). With the six other LLM backbones, SPIDER achieved entity F1 scores of 0.717–0.835 and compatibility-based triple F1 scores of 0.436–0.589, all exceeding the corresponding scores of the conventional methods.

Compared with single-pass prompting using the same DeepSeek-V4-Flash backbone, the multi-agent workflow increased entity F1 from 0.761 to 0.928 and compatibility-based triple F1 from 0.468 to 0.793 (Fig. 3a). Based on these results, we used SPIDER with DeepSeek-V4-Flash for all subsequent literature-grounding analyses.

### Literature grounding expands evidence coverage of biomedical knowledge

We applied SPIDER to abstracts from 1.95 million PubMed publications to augment the research-oriented subgraph, denoting the original and augmented graphs as *L*_0_ and *L*_1_, respectively. The entity set remained unchanged, while augmentation added 2.31 million previously unrepresented entity pairs linked by literature-supported relations. The proportion of entities with at least one supporting publication increased from 14.6% in *L*_0_ to 65.0% in *L*_1_. For entity pairs, the corresponding proportion increased from 34.0% of pairs in *L*_0_ to 79.4% of pairs in *L*_1_. Literature grounding added supporting publications across all entity and relation types (Fig. 3c).

In *L*_1_, 56.9% of all entity pairs had one supporting publication, whereas approximately 22.5% had two or more (Fig. 3d). For each relation, we retained the supporting text and PMID from each publication in a separate evidence record. These records enabled comparisons of findings for the same entity pair across publications, providing the basis for the subsequent analysis of potentially incompatible relations.

### Evidence and study context help distinguish apparent conflicts from genuine disagreement

We screened *L*_1_ for entity pairs connected by potentially incompatible relations. Overall, 13.4% of entity pairs were connected by two or more relation types; among these, 9.7% (44,228 pairs) matched predefined incompatibility patterns (Fig. 3e; Methods, “Identification of potentially incompatible relations”). Most candidate pairs involved relations extracted from the literature, with fewer involving relations from database records or both sources (Fig. 3f). The most common pattern paired an explicitly reported absence of a relationship (`NOT_ASSOCIATED_WITH`) with another relation type for the same entity pair. Other patterns included positive versus negative regulation, similarity versus difference, and treatment versus contraindication (Fig. 3g).

To distinguish context-dependent differences from genuine disagreement under comparable study conditions, we defined a comparison unit as a distinct pair of potentially incompatible relations for the same entity pair. Each unit included the literature evidence records associated with both relations, and we retained only units with literature support for each. Because an entity pair could yield multiple relation combinations, applying these criteria to the 44,228 candidate pairs produced 89,937 comparison units (Methods, “Identification of potentially incompatible relations”). Using DeepSeek-V4-Flash, SPIDER classified each unit based on its supporting text and extracted study conditions. The model classified 79.0% of units as context-dependent differences, 6.1% as genuine disagreement and 2.9% as relation-mapping errors; 12.1% remained unresolved (Fig. 4b). Taken together, these classifications suggest that most apparent incompatibilities in the analyzed comparisons reflected differences in study conditions rather than genuine scientific disagreement. These results also showed that potentially incompatible relation types alone do not establish genuine disagreement; their interpretation also requires examination of the supporting evidence and study conditions.

**Figure 4.**
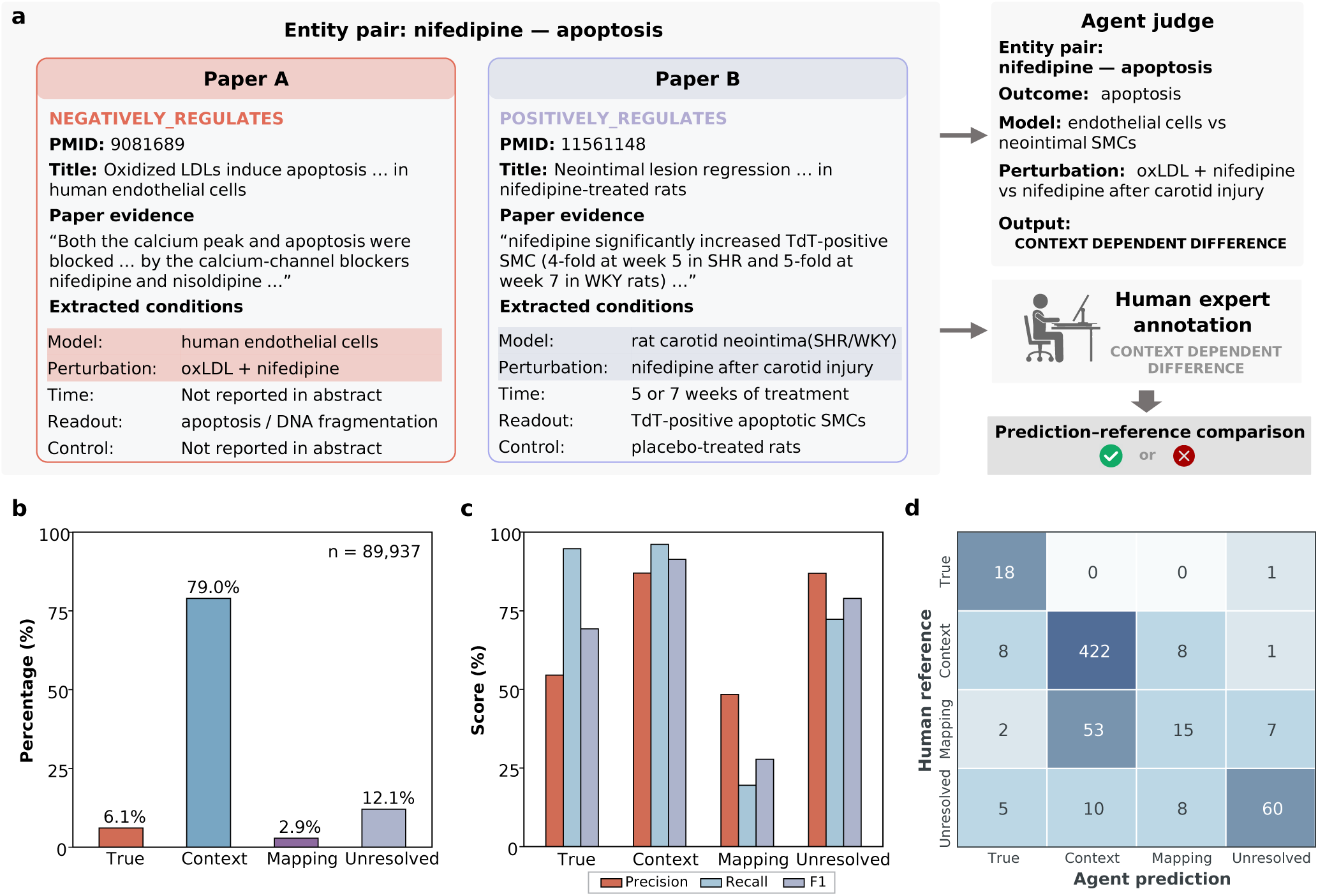
Condition-aware interpretation of potentially conflicting biomedical relations. **a**, Nifedipine–apoptosis example showing how supporting evidence and study conditions distinguish context-dependent findings. **b,** Classification of 89,937 evidence-supported comparison units into context-dependent differences, genuine disagreement, relation-mapping errors, and unresolved cases. **c,** Class-specific precision, recall and F1 scores against expert annotations of 618 sampled comparisons. **d,** Confusion matrix comparing SPIDER classifications with expert-reference labels.

The nifedipine–apoptosis comparison illustrates how study conditions informed classification (Fig. 4a). One study reported that nifedipine inhibited apoptosis in human endothelial cells treated with oxidized LDL (PMID 9081689^28^). Another reported an increase in apoptotic smooth-muscle cells in the rat carotid neointima after 5–7 weeks (PMID 11561148^29^). Given the differences in biological models and treatment conditions, SPIDER classified this comparison as a context-dependent difference. Additional examples are provided in Appendix S1.

To evaluate classification performance, eight medical professionals provided reference annotations for 618 randomly sampled comparison units using the same four categories. Against these annotations, SPIDER achieved an accuracy of 83.3% across all four categories and a macro-F1 of 80.29% averaged over the two categories of context-dependent differences and genuine disagreement (Fig. 4c,d). The model achieved high recall for both of these categories but tended to misclassify relation-mapping errors as context-dependent differences. Together, these results suggest that using SPIDER to compare supporting evidence and reported study conditions helps interpret potentially incompatible relations.

### Structured study conditions improve question answering about conflicting findings

We used SPIDER-Bench to evaluate whether structured study conditions extracted by SPIDER improve question answering about conflicting findings. The benchmark contains 200 multiple-choice questions constructed from 118 comparison units, each based on potentially incompatible findings involving the same entity pair. We used the same answering model in both comparisons. We first compared an LLM-only baseline with four input configurations from SPIDER-KG: triples alone, triples with structured study conditions, triples with literature evidence, and triples with both. Compared with the baseline, providing triples with structured study conditions significantly increased accuracy from 50.5% to 77.0% and macro-F1 from 50.3% to 77.0%. Both scores approached those obtained with triples and literature evidence or with all three input types. Triples alone did not significantly improve either metric (Fig. 5a). These results suggest that structured study conditions capture much of the information useful for answering questions about conflicting findings in SPIDER-Bench.

We next compared question answering supported by SPIDER-KG with that supported by five other biomedical KGs, using the same retrieval workflow throughout. SPIDER-KG supplied triples, structured study conditions and literature evidence. CROssBARv2-KG^30^ and MedKGent-KG^31^ supplied relations linked to literature evidence, whereas RTX-KG2^9^, BIOS v3^32^ and PrimeKG^2^ supplied graph relations alone. The model achieved the highest accuracy and macro-F1 with SPIDER-KG, with accuracy reaching 78.5% compared with 42.5%–50.0% for the other KG-supported settings (Fig. 5b).

### SPIDER-KG improves performance across biomedical reasoning tasks

We evaluated SPIDER-KG on four biomedical reasoning benchmarks, comparing an LLM answering directly with the same model supported by SPIDER-KG or other biomedical KGs. Within each task, all systems used the same answering model, and KG-supported systems used a common retrieval workflow (Fig. 5e).

On BioMixQA, which tests discrimination between true and false biomedical statements^33^, SPIDER-KG increased accuracy from 87.46% with the LLM alone to 95.50%. CIViC assesses whether a publication supports a clinical interpretation in a specified molecular, disease and treatment context^35^. For this task, KG-supported systems retrieved relevant text from the specified publication when available in their respective KGs’ fixed literature collections. SPIDER-KG increased accuracy from 54.27% to 68.44% (Fig. 5c). Improvements were also observed on the subsets of questions for which SPIDER-KG supplied evidence to the answering model (Fig. 5d).

We assessed gene prioritization using the Open Targets GWAS benchmark, in which each system ranked the same predefined set of candidate effector genes for each locus–trait pair^34^. SPIDER-KG increased mean reciprocal rank (MRR) from 0.8195 to 0.8630, close to the highest score of 0.8674 achieved by MedKGent-KG (Fig. 5c).

DICTrank evaluates the classification of drugs into cardiotoxicity-concern categories derived from drug labeling^36^. SPIDER-KG modestly increased macro-F1 from 0.4664 to 0.4942 in the three-class setting and from 0.3476 to 0.3718 in the four-class setting (Fig. 5c). Across all four benchmarks, SPIDER-KG improved on the LLM-only baseline and achieved the highest accuracy among the evaluated systems on BioMixQA and CIViC.

### SPIDER-KG helps interpret newly reported gene–disease associations

We retrospectively examined whether SPIDER-KG could connect newly reported gene–disease associations to relevant prior findings in 22 cases with genetic or functional evidence reported in 2026 (Table S3). The analysis included cases without a direct gene–disease association in the graph.

**Figure 5.**
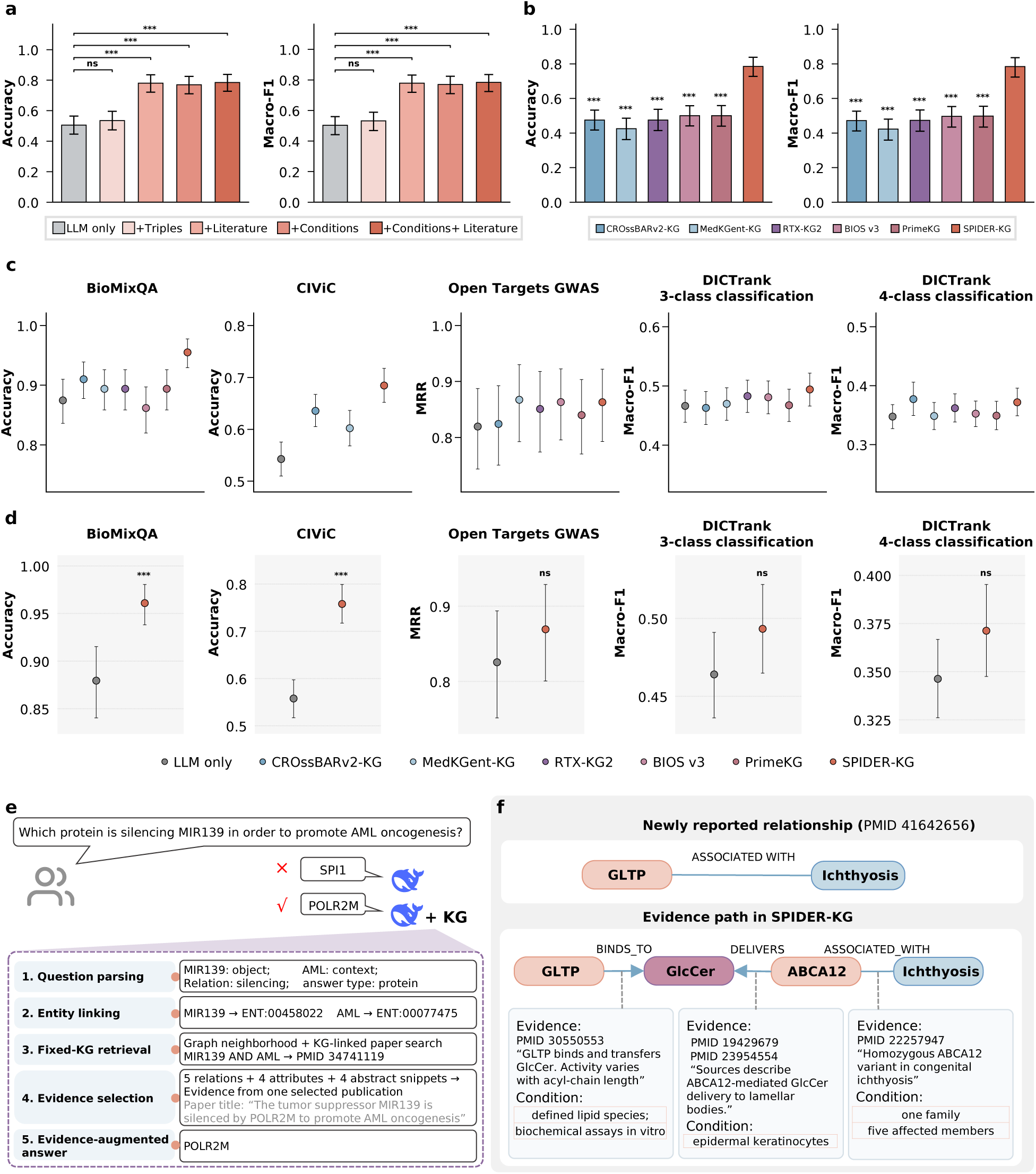
Evaluation of SPIDER-KG on SPIDER-Bench and four downstream biomedical tasks. **a,b**, SPIDER-Bench performance across evidence-input settings (**a**) and KGs (**b**). **c,** LLM-only and KG-augmented performance on BioMixQA^33^, Open Targets GWAS gold standards^34^, CIViC^35^ and DICTrank^36^ (three- and four-class settings). **d,** LLM-only versus SPIDER-KG-augmented performance on evidence-available subsets. **e,** KG-augmented answering workflow. **f,** GLTP–ichthyosis evidence path through GlcCer and ABCA12. Error bars in **a–d** show pointwise 95% cluster-bootstrap confidence intervals (10,000 resamples). Significance in **a,b,d** used two-sided paired cluster permutation tests (10,000 permutations), with separate Holm correction across eight tests in **a**, ten in **b** (both metrics jointly), and five primary-metric tests in **d**. References were LLM-only in **a,d** and SPIDER-KG in **b**. ns, *P*_adj_ *≥* 0.05; *, *P*_adj_ *<* 0.05; **, *P*_adj_ *<* 0.01; ***, *P*_adj_ *<* 0.001.

For example, a study published in February 2026 identified biallelic GLTP variants in five unrelated families with an epidermal differentiation disorder presenting with ichthyosis, with supporting evidence from cellular and mouse models^37^. Although SPIDER-KG lacked a direct GLTP–ichthyosis association, it connected the gene and disease through glucosylceramide (GlcCer) and ABCA12 (Fig. 5f).

The supporting publications all predated the 2026 GLTP study. Biochemical experiments showed that GLTP binds and transfers GlcCer^38^. Another publication described ABCA12-mediated delivery of GlcCer to epidermal lamellar bodies as established background^39^, while a review discussed ABCA12’s role in epidermal lipid transport^40^. A family study linked a homozygous ABCA12 variant to non-bullous congenital ichthyosiform erythroderma^41^.

These findings connect GLTP’s lipid-transfer activity, a shared lipid substrate and an epidermal transporter implicated in ichthyosis, providing biological context for the newly reported association. They do not establish functional coupling between GLTP and ABCA12 or explain how GLTP variants cause the disease. This case illustrates how SPIDER-KG can assemble prior evidence relevant to a candidate explanation while identifying mechanistic questions that remain to be tested.

## DISCUSSION

### SPIDER-KG links biomedical relations to the evidence and study conditions needed to interpret them

Its three linked layers preserve graph relations, supporting literature evidence and reported study conditions, allowing findings concerning the same entities to be examined individually. This organization supports tracing assertions to their sources, comparing findings across studies and retrieving evidence relevant to a specific question. The incremental update pipeline allows newly extracted relations and evidence to be incorporated as additional PubMed records become available.

### Study context is central to interpreting apparent conflicts

Most screened, evidence-supported comparisons were classified as context-dependent differences. This finding suggests that potentially incompatible relation types provide a starting point for further assessment, while establishing genuine disagreement requires determining whether the underlying findings concern comparable biological and experimental settings. Preserving negative and context-specific observations makes these distinctions available for scrutiny. More broadly, this perspective empha-sizes the applicability of individual findings as an essential component of biomedical knowledge interpretation.

### Structured study conditions provide useful information for biomedical reasonin

On SPIDER-Bench, triples paired with structured study conditions improved performance over the LLM-only baseline and approached the scores obtained with triples and literature evidence, whereas triples alone did not yield statistically significant improvements. These results suggest that extracted conditions capture information useful for interpreting conflicting findings. Improvements across four additional benchmarks support the broader utility of SPIDER-KG for biomedical knowledge assessment, evidence interpretation, gene prioritization and drug safety classification, although gains varied across tasks. The accompanying interface enables researchers to inspect the relations, evidence sources and conditions underlying retrieved information.

### Evidence connections can inform mechanistic exploration and hypothesis development

The retrospective gene–disease cases illustrate how earlier studies can provide biological context for newly reported associations, including when the graph lacks a direct relation. Connecting these findings makes the evidence relevant to candidate explanations accessible and highlights missing mechanistic links. Such connections may help researchers identify where independent findings converge and design targeted experiments to test possible mechanisms. Extending this approach to relationships among compounds, proteins, biological processes and phenotypes could broaden its use in hypothesis development.

## FUTURE WORK

### Broader literature coverage and more flexible representations could extend the scope of SPIDER-KG

The current framework primarily uses PubMed titles and abstracts and operates within a predefined entity set, incompatibility patterns and condition templates. Relevant details available only in full text or supplementary materials may therefore be missed, and incomplete study conditions can leave the comparability of findings uncertain. Incorporating these additional sources and expanding the representations of study context could support a wider range of findings and scientific questions.

### Refining extraction and classification could further strengthen evidence interpretation

Errors in entity alignment, relation and condition extraction, and conflict classification can propagate into subsequent analyses. Expert evaluation identified a tendency to classify relation-mapping errors as context-dependent differences, indicating a specific opportunity for refinement. The reported classification proportions should therefore be interpreted alongside the expert-validation results and within the screened, evidence-supported comparison set. Additional expert annotations and targeted improvements to relation mapping could help distinguish these categories more reliably.

### Further evaluation could extend the demonstrated utility to additional research settings

The current benchmarks show improvements across several biomedical tasks. Evaluation on independently constructed benchmarks with broader question types and biological contexts could further characterize generalization. Because the KGs supplied different information, comparisons that more closely match evidence coverage could help clarify the contributions of literature evidence and structured study conditions. The retrospective gene–disease cases also motivate prospective studies to assess hypothesis prioritization and experimental follow-up to test the mechanisms suggested by connected evidence.

## METHODS

We constructed SPIDER-KG by integrating heterogeneous biomedical databases and ontologies, harmonizing entities and relations while retaining available source-database names, original record identifiers and literature references. We then applied SPIDER to augment the graph with literature evidence, recover study-specific conditions, and enable evidence-grounded interpretation of biomedical relationships. Here, we describe the graph representation, construction, and literature augmentation, followed by condition extraction and downstream interpretation analyses.

### Core representation of SPIDER-KG

SPIDER-KG contains entities, standardized triples and supporting evidence records:

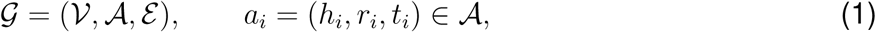

where *V* and *A* are the sets of entities and standardized triples, respectively, and ε associates each entity or triple with its source-specific evidence records. Each triple connects a head entity *h_i_* to a tail entity *t_i_* through a relation type *r_i_*.

For an entity or triple *x*, ε (*x*) denotes its supporting evidence records. These include database records, ε ^db^(*x*), which retain source-database names and original record identifiers where available, and literature records, ε ^lit^(*x*), which retain the PMID and supporting text.

Each standardized triple is stored once but may have multiple supporting records. We represent a triple paired with its *j*-th evidence record as an evidence-linked triple:

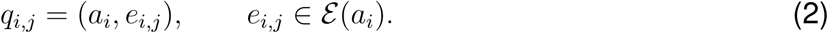

This allows findings supporting the same relation to be examined separately.

For literature evidence records selected for condition extraction, Λ(*e_i,j_*) denotes the structured study conditions recovered from the supporting text. These conditions remain linked to the individual evidence record. If no conditions are recovered, Λ(*e_i,j_*) = Ø, and the evidence-linked triple remains available.

### Construction of the integrated biomedical knowledge graph

#### Data sources

We constructed the initial graph by integrating 48 biomedical databases and ontologies^42–89^. Resources were selected according to four criteria: (i) accessibility through bulk download, FTP, version-controlled repositories or programmatic interfaces, including registered or licensed access, or, where no bulk distribution existed, automated retrieval of public web pages; (ii) documented provenance, whether from expert curation, experimental measurement or computational pipelines, with stable identifiers; (iii) coverage of entity or relation types within the scope of the graph, including resources no longer actively updated; and (iv) traceability through source identifiers or stable URLs, together with the release version or retrieval date. Integrative resources were retained where they contributed relations or cross-references unavailable from the original sources. The complete resource list is provided in Supplementary Table S16.

#### Schema harmonization

##### Entity harmonization

We assigned source entities to 16 predefined types, including genes, proteins, drugs, metabolites, cells, diseases, and phenotypes, primarily based on each resource’s classifications (Supplementary Table S17). Shared identifiers and normalized names or synonyms were used to identify candidate matches. Records were merged only when identifier-based or type-specific evidence supported a shared identity and the records passed the consistency checks described in Appendix S2.2. Each consolidated entity received a sequential identifier unique within the build (for example, ENT:00000001) and retained its source identifiers, available descriptive metadata and one or more compatible entity types.

##### Relation harmonization

We mapped source relation descriptions to a set (*R*) of 58 standardized relation types, covering associations, molecular interactions, regulatory effects and therapeutic relationships (Supplementary Table S19). Source relation descriptions were normalized and mapped using predefined rules. For descriptions not covered by these rules, DeepSeek-V4-Flash^27^ was used to assign a standardized relation type. Mapping rules and prompts are detailed in Appendix S2.1. Original descriptions were retained alongside their standardized types.

#### Graph construction

We linked relation endpoints to consolidated entities using source identifiers, with names and synonyms used when identifier-based matching was unavailable (Appendix S2.3). Records with both endpoints resolved were converted into triples using the standardized relation types. Records with unresolved endpoints were excluded from the graph but retained separately for inspection. Triples were deduplicated by head entity, relation type, and tail entity; different relation types connecting the same entity pair remained separate triples.

Available database evidence records were linked to the corresponding entities or triples, retaining source-database annotations, source-local identifiers, literature references and original relation descriptions where available.

### Multi-agent system for literature-based graph augmentation

#### System architecture

SPIDER takes an existing biomedical KG and a collection of PubMed records as inputs and augments the graph with literature-supported relations and evidence. Entity names and synonyms guide literature retrieval and alignment to existing nodes. The same workflow processes the initial literature collection and subsequent update batches.

Each retrieved abstract is processed by three specialized agents:

1. **Entity Identification and Consolidation Agent.** Identifies entities, assigns extraction categories, and consolidates names and aliases referring to the same entity.
2. **Relation Discovery Agent.** Uses the entity inventory and abstract text to propose candidate relations and identify supporting evidence, including evidence spanning multiple sentences.
3. **Relation Classification Agent.** Assigns standardized relation types by checking candidates against the supporting text, entity categories, and relation definitions, including direction and polarity where applicable.

Structured outputs preserve entity references and supporting text between different agents. Candidate relations are excluded when the source text does not provide supporting evidence or when they violate the schema. Agents that return no valid output are retried up to a predefined limit; processing stops if valid output remains unavailable. Agent prompts, model settings, validation checks and retry procedures are detailed in Appendix S2.6.

We use 9 entity categories and 58 standardized relation types in *R* for literature extraction, with relation-specific constraints on head–tail entity-category combinations. The extraction categories were defined separately from the 16 KG entity types, which primarily reflect source-database classifications, as some of these finer-grained distinctions may not be explicit in scientific text. For example, we grouped drugs, metabolites, and other chemical substances under `CHEMICAL`, while including `DEVICE` and `PROCEDURE` to capture additional concepts in the literature. Extracted entities were aligned to existing KG nodes, and only relations with both entities successfully aligned were integrated into the graph. No new entities were introduced during this process.

#### Literature-based graph augmentation

For literature grounding and subsequent conflict analysis, we constructed a research-oriented subgraph through entity-type filtering, exclusion of structural relation types, and PubMed-based screening of entities. The resulting subgraph comprised 198,659 entities and 1,099,652 unique undirected entity pairs (Appendix S2.5). We used this subgraph as the basis for subsequent literature retrieval and provenance analysis. Provenance coverage was assessed by whether an entity or entity pair had a recorded source-database name, a PMID link, or both. “No provenance” denotes entities or entity pairs with neither a recorded source-database name nor a PMID link in the analyzed graph.

To identify relevant publications for evidence extraction, we searched a PubMed snapshot dated 30 December 2025 using Elasticsearch, with queries based on individual KG entities and entity pairs. Entity-based searches retrieved publications by matching KG entity names and synonyms against titles, abstracts, and MeSH major-topic terms, providing broad coverage of literature relevant to individual entities. Entity-pair-based searches required both entities to appear in the title or abstract, focusing on publications that might describe relationships between them. Results from both strategies were ranked by textual relevance and publication recency, with additional criteria including review status and title or MeSH matches for entity-based searches, and entity co-occurrence and relation-keyword matches for entity-pair-based searches. We retained up to 15 publications per query and pooled the retrieved PMIDs separately for each strategy. To prioritize publications identified by both retrieval strategies, we retained the intersection of the two PMID sets, yielding 1,953,760 unique PubMed records for evidence extraction.

The multi-agent system processed the selected PubMed records to extract entities, candidate triples and supporting evidence. We first aligned extracted entities to nodes in the fixed research-oriented subgraph through exact matching against normalized KG names and aliases. For entities without an exact match, we used BioLORD-2023^90^ embeddings to retrieve the node with the highest cosine similarity. We accepted the candidate when its similarity reached 0.85 and left the entity unmapped otherwise. We selected the similarity threshold using 100 manually annotated entity–candidate pairs as a development set. Thresholds from 0.50 to 1.00 were evaluated in increments of 0.05, and 0.85 achieved the highest development-set balanced accuracy of 92.9%. The threshold was fixed before evaluation on an independent set of 300 entity–candidate pairs, yielding a precision of 95.43% (95% Wilson confidence interval, 91.24–97.67%) and a balanced accuracy of 93.06% (Appendix S2.13.2).

Extracted relations were integrated into SPIDER-KG by adding new standardized triples or attaching literature evidence to existing triples. Each standardized triple was stored once, while evidence records from different publications were retained separately to preserve their provenance. Literature evidence for aligned entity mentions was also linked to the corresponding KG entities. For downstream evidence prioritization, extraction confidence was combined with publication metadata using the ranking function described in Appendix S2.4.

For incremental updates, SPIDER runs daily and retrieves PubMed records newly indexed since the preceding successful update. It deduplicates these records by PMID and applies the same extraction, alignment, and integration workflow. These updates add evidence and relations among the retained entities without rebuilding the graph constructed from databases. All analyses reported in this study used a fixed graph snapshot and were independent of subsequent updates.

### Condition-aware analysis of potentially incompatible relations

#### Identification of potentially incompatible relations

We screened standardized triples for four predefined patterns of potential incompatibility: (i) `NOT_ASSOCIATED_WITH` paired with any other relation type; (ii) `POSITIVELY_REGULATES` paired with `NEGATIVELY_REGULATES`; (iii) `SIMILAR_TO` paired with `DIFFERENT_FROM`; and (iv) `TREATS` paired with `CONTRAINDICATED_FOR`. Here, `NOT_ASSOCIATED_WITH` denotes an explicitly reported negative find-ing. These patterns identified candidates for evidence-based comparison but did not themselves establish genuine scientific disagreement.

Each comparison unit was defined by an entity pair, a distinct combination of potentially incompatible relation types and, where applicable, direction. For patterns (ii) and (iv), matching was direction-sensitive: both triples had to have identical head and tail entities, and matches in the reverse direction were counted separately. For patterns (i) and (iii), matching was direction-invariant: triples were matched by their unordered entity pair, irrespective of stored endpoint order.

The 44,228 candidate entity pairs could each contribute multiple comparison units, each comparing a specific pair of potentially incompatible relations. For each relation in a comparison unit, we collected all supporting literature evidence records, retaining multiple supporting publications within the same unit. We retained only units with usable literature evidence for both relations being compared, yielding 89,937 comparison units for subsequent analysis of evidence and study conditions.

#### Condition schema and extraction

We extracted study conditions from the literature evidence records associated with the comparison units retained above. This step captured the biological, experimental and observational variables needed to assess whether potentially incompatible findings arose under comparable conditions. We defined 18 condition templates based on the incompatibility patterns and endpoint extraction categories. For comparisons involving `NOT_ASSOCIATED_WITH`, template selection depended on the semantic family of the paired relation. For the other three patterns, selection used the incompatibility pattern and the unordered pair of endpoint extraction categories. Depending on the template, fields captured the study population or experimental model, entity or disease state, intervention or exposure, measured attribute or outcome, timing, analytical method or decision criterion, and control or reference condition.

For each applicable evidence-record–template pair, an LLM-based extraction module received the standardized triple, relation definition, selected template and the corresponding publication title and abstract. The module extracted and normalized condition values and retained the source excerpts supporting each value.

For a literature evidence record *e*, the extracted condition instances were represented as

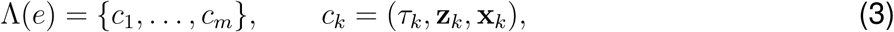

where *m* is the number of condition instances retained for evidence record *e*, *τ_k_* identifies the template, **z***_k_* contains the structured condition values, and **x***_k_* contains the source excerpts supporting those values. Multiple instances were retained when the evidence described the triple under distinct study conditions. If no conditions were recovered, *m* = 0 and Λ(*e*) = ∅Ø, and the evidence record remained available for classification.

The conditions remained linked to their evidence record. For an evidence-linked triple *q_i,j_*= (*a_i_, e_i,j_*), the annotated representation was

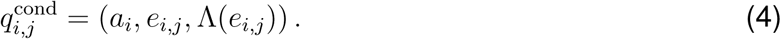

Template definitions and selection rules are provided in the GitHub code repository. Extraction prompts, structured output schemas and model settings are detailed in Appendix S2.7.

#### Condition-aware classification

An LLM-based module assigned each eligible comparison unit to one of four categories using its supporting evidence and extracted study conditions:

- `TRUE_CONFLICT`. Source evidence supported both relations, but their conclusions about the same attribute or outcome were incompatible under scientifically comparable conditions.
- `CONTEXT_DEPENDENT_DIFFERENCE`. Source evidence supported both relations, and differences in relevant study conditions could account for their divergent conclusions.
- `RELATION_MAPPING_ERROR`. For at least one relation, every available evidence record indicated a mismatch between the source evidence and the mapped entities, relation type or direction.
- `UNRESOLVED`. The available evidence was insufficient to assign one of the other three categories.

The module received the entity pair, the labels and definitions of the relations being compared, the required condition fields, and the supporting evidence records and extracted conditions for each relation. It first checked whether the evidence supported the mapped entities, relation types and directions. For supported relations, it assessed whether the findings expressed incompatible conclusions about the same attribute or outcome. It then examined whether differences in study conditions could explain the incompatibility or whether the findings conflicted under scientifically comparable conditions. Equivalent descriptions and convertible measurement units were considered, while missing condition values were treated as uncertainty.

The module considered all evidence records supplied for the comparison and returned one label with supporting source excerpts. Raw publication titles and abstracts were not provided at this stage. Complete prompts, output schemas and model settings are provided in Appendix S2.8.

### Evaluation design

We evaluated SPIDER and SPIDER-KG through complementary assessments of literature extraction, condition-aware classification, conflict-aware question answering and downstream biomedical tasks.

#### Literature extraction benchmark

##### Benchmark construction

We randomly sampled 1,000 PubMed abstracts available as of 16 March 2026. Reference annotations were constructed using a model-assisted annotation procedure. Initial entity and text-supported triple annotations were generated by DeepSeek-V4-Flash in a single pass, following the direct-baseline approach. Four biomedical annotators revised these drafts by correcting errors, adding missing annotations, and consolidating names and aliases referring to the same entity within each abstract. A separate panel of four biomedical experts cross-reviewed the revised annotations. Disagreements were resolved before the reference dataset was finalized. The benchmark contained 15,468 reference entities across 9 extraction categories and 14,165 text-supported triples spanning 58 relation types.

##### Experimental setup and evaluation

We evaluated the extraction workflow with seven LLM backbones: DeepSeek-V4-Flash^27^, DeepSeek-V3.2^91^, Gemini 3.1 Flash-Lite^92^, Claude Sonnet 4.6^93^, GPT-5.5^94^, Grok 4.1 Fast^95^ and MiniMax M2.5^96^. Task instructions, the agent work-flow, extraction schema and post-processing procedures were held constant across backbones. DeepSeek-V4-Flash served as the primary backbone and was also evaluated with single-pass prompting to assess the contribution of the staged workflow.

We additionally compared SPIDER with three specialized extraction systems: BERN2^24^, PubTator^25^ and GLiNER-Relex^26^. BERN2 was evaluated only for entity extraction because it does not predict relations. System-specific entity categories were mapped to the 9 benchmark categories before scoring (Appendix S2.9.1).

We matched predicted and reference entities within the same abstract and harmonized entity category using BioLORD-2023 embeddings^90^. A cosine similarity of at least 0.70 defined an eligible match. We selected this threshold by maximizing entity-pair matching F1 on 50 development abstracts outside the extraction benchmark (Appendix S2.13.1). On 50 separate validation abstracts, it achieved a matching precision of 93.64%, recall of 91.76%, and F1 of 92.69%. We applied the same threshold to all evaluated systems.

After entity alignment, relation labels were normalized to the benchmark vocabulary (Appendix S2.9.2). Triple evaluation used compatibility-based matching: predicted and reference triples within the same abstract had to connect the same aligned entity pair and satisfy label-dependent directionality and predefined incompatibility constraints, without requiring identical relation labels (Appendix S2.9.3). These metrics assess compatibility with the reference annotations rather than exact relation-label accuracy.

For entity extraction, precision was the proportion of predictions with at least one eligible reference match, and recall was the proportion of reference entities matched by at least one prediction. Compatibility-based triple precision and recall were calculated analogously using compatible matches. Overall precision and recall were calculated by pooling their respective numerator and denominator counts across all 1,000 abstracts, and *F*_1_ was their harmonic mean. Entity scores were also reported separately for each of the 9 extraction categories.

##### Expert validation of condition-aware classification

We randomly sampled 618 evidence-eligible comparison units from the 89,937 units included in the condition-aware analysis. Eight medical professionals independently assigned one of the four predefined labels while blinded to the classification module’s predictions. They assessed the relations being compared, their supporting literature evidence and structured study conditions. Disagreements were resolved by consensus to establish the reference labels. The primary metric was macro-*F*_1_ averaged over `TRUE_CONFLICT` and `CONTEXT_DEPENDENT_DIFFERENCE`, with all 618 cases retained in the calculation. We additionally reported class-specific *F*_1_, overall four-class accuracy and the confusion matrix.

#### SPIDER-Bench: conflict-aware question-answering benchmark

##### Benchmark construction

We constructed SPIDER-Bench to assess relation inference under specified study conditions and recognition of genuine scientific disagreement. Questions were derived from context-dependent differences and true conflicts retained after provenance-based quality review. Each conflict group corresponded to a comparison unit comprising two potentially incompatible relations for the same entity pair and their supporting evidence.

For context-dependent differences, questions specified the conditions needed to identify the applicable relation. Questions concerning the same entity pair could therefore have different correct answers under different study conditions. For true conflicts, questions specified shared or scientifically comparable conditions, with the reference answer indicating that the evidence did not support a unique relation.

All questions used a four-option format. Options A and B represented the two competing relation outcomes, C indicated no significant effect or partial or conditional applicability, and D indicated an unresolved conflict between the two supported outcomes under the stated conditions. The stated conditions and reference answers were required to be traceable to source publications. At least one biomedical expert verified each question for evidential support, answer consistency and clarity. The final benchmark contained 200 questions from 118 conflict groups: 191 condition-resolution questions from 109 context-dependent-difference groups and 9 true-conflict-recognition questions from 9 true-conflict groups. Question-construction and exclusion criteria are detailed in Appendix S2.10.

##### Experimental setup

Retrieval used only the question text and answer options, without access to reference answers, conflict labels or evidence annotations used during question construction.

The pipeline identified entities and study conditions in each question, linked the entities to KG nodes and retrieved direct relations between them. Triples and source publications were selected according to the question semantics and condition cues. Structured conditions were obtained from extraction results associated with the selected publications, whereas literature evidence comprised publication titles and abstracts.

Using a frozen SPIDER-KG snapshot, we compared five information configurations: LLM only, triples alone, triples with literature evidence, triples with structured study conditions, and triples with both conditions and literature evidence. The retrieved triples and publications were held constant across the four KG-supported configurations, with only the information supplied to the answering model varied. All SPIDER-Bench configurations used DeepSeek-V4-Flash with a temperature of 0 and a maximum output length of 384 tokens. They were evaluated on identical questions with the same answer-option ordering. Questions without retrieved evidence remained in the evaluation.

For the cross-KG comparison, we used a common question-driven retrieval and answering framework with SPIDER-KG and five external resources: CROssBARv2-KG, MedKGent-KG, RTX-KG2, BIOS v3 and PrimeKG^2,9,30–32^. Entity linking was adapted to each resource. SPIDER-KG used exact-name matching followed by BioLORD-based matching, whereas the external KGs used normalized exact matching followed by FTS5 lexical matching. SPIDER-KG supplied triples, structured study conditions and literature evidence. CROssBARv2-KG and MedKGent-KG supplied triples and literature evidence; RTX-KG2, BIOS v3 and PrimeKG supplied triples alone. Each resource contributed the fullest information available through its adapter, without a matched evidence budget.

##### Evaluation metrics and statistical analysis

For SPIDER-Bench, we reported accuracy and macro-*F*_1_, with macro-*F*_1_ calculated across the four displayed answer labels (A–D). Detailed metric definitions and statistical procedures are provided in Appendix S2.11.

We estimated 95% percentile confidence intervals using 10,000 bootstrap resamples of conflict groups. Questions belonging to the same group were kept together, and identical resamples were used across configurations within each analysis. Two-sided paired permutation tests used 10,000 permutations, with predictions swapped between configurations jointly for all questions within each conflict group. Holm correction was applied jointly across accuracy and macro-*F*_1_ within each of two separate comparison families, rather than separately for each metric. The information-ablation family comprised eight tests: each of the four configurations (triples alone, triples with literature evidence, triples with structured study conditions, and triples with both conditions and literature evidence) was compared with the LLM-only reference on both metrics. The cross-KG family comprised ten tests: each of the five baselines (CROssBARv2-KG, MedKGent-KG, RTX-KG2, BIOS v3, and PrimeKG) was compared with SPIDER-KG on both metrics. Statistical significance was assessed at a Holm-adjusted *P <* 0.05.

##### Interactive interface

We implemented a web interface for free-text biomedical questions (https://spiderkg.internal.onexai.com.cn). The interface uses the question-driven retrieval workflow to generate answers supported by KG relations, literature evidence and available study conditions. It displays the supporting subgraph and source links for inspection.

#### Downstream biomedical task evaluation

We evaluated four additional benchmarks covering biomedical knowledge judgment, oncology evidence assessment, effector-gene prioritization and drug safety classification. Within each task, we compared direct-answer DeepSeek-V4-Flash (deepseek-v4-flash; temperature = 0) with the same model supported by SPIDER-KG or the external KGs described above. Public task inputs, answering-model settings, output requirements and scoring were held constant across configurations. KG-supported configurations used a common evidence-selection procedure with resource-specific entity linking and knowledge access. Retrieval was restricted to fixed KG snapshots and their linked literature evidence.

##### BioMixQA

We used the complete 311-question true/false subset of BioMixQA^33^, retaining the original questions and reference labels. KG-supported retrieval selected relevant relations, entity attributes, source definitions and literature evidence according to each statement and its contextual constraints, including negation and disease context. Performance was measured by accuracy.

##### Open Targets GWAS gold standards

We constructed 138 locus–trait ranking tasks from high-confidence, functionally supported records in the Open Targets GWAS gold standards^34^. For each task, all configurations ranked the same complete candidate set: protein-coding genes with a GENCODE v49 transcription start site within 500 kb of the sentinel variant. KG-supported retrieval collected trait-relevant relations, gene attributes and literature evidence for each candidate. Performance was measured by mean reciprocal rank (MRR), using the rank of the highest-ranked reference gene for each task.

##### CIViC

We constructed 1,616 questions from the 1 February 2026 CIViC release^35^. Each question asked whether a specified publication supported a stated clinical interpretation, preserving the molecular-profile, variant, disease, variant-origin and treatment conditions. The LLM-only baseline received the question, including these conditions, the stated interpretation, the source type and publication identifier (e.g., PMID), together with the answer instructions; no publication title, abstract or full text was provided. KG-supported configurations received the same question and answer instructions, supplemented with relevant text retrieved from the specified publication when available in the corresponding KG’s fixed literature collection. Performance was measured by accuracy.

##### DICTrank

DICTrank is an FDA reference dataset that assigns 1,318 human drugs to cardiotoxicity-concern categories based on drug labeling^36^. We evaluated a three-class task comprising 1,211 drugs labeled NO, LESS or MOST concern, and a four-class task that additionally included 107 AMBIGUOUS records. All configurations received the same public category definitions. KG-supported retrieval selected relevant drug attributes, relations and literature evidence while preserving drug identity and exposure context. Performance was measured by macro-*F*_1_ across the categories in each task.

##### Scoring and statistical analysis

All instances in each fixed cohort were retained in scoring, including those with retrieval failures, abstentions or invalid responses. For CIViC and BioMixQA accuracy, abstentions and invalid responses counted as incorrect. For DICTrank macro-F1, they counted as false negatives for the reference class, without introducing an additional class. For Open Targets GWAS MRR, abstentions and invalid rankings received zero. Rankings had to include every candidate exactly once in a strict order; incomplete rankings or tied ranks received zero, with no completion or tie averaging. We compared the LLM-only baseline with SPIDER-KG-supported and applicable external-KG configurations on the full cohorts. We additionally compared the LLM-only and SPIDER-KG-supported configurations on matched subsets of instances for which SPIDER-KG supplied evidence.

For both full cohorts and evidence-available subsets, pointwise 95% percentile confidence intervals for primary-metric estimates were calculated using 10,000 cluster-bootstrap resamples.

Subset comparisons against the LLM-only baseline used two-sided paired cluster-permutation tests with 10,000 resamples. The resulting *P* values were jointly adjusted using the Holm procedure across five primary-metric comparisons: CIViC accuracy, BioMixQA accuracy, GWAS MRR, and DICTrank three-and four-class macro-F1. Subset definitions and task-specific clustering procedures are provided in Appendix S2.12.

## DATA AVAILABILITY

Data of SPIDER-KG are publicly available at https://zenodo.org/records/23009923. The primary biomedical databases and ontologies integrated into SPIDER-KG, including source versions, access information, and retrieval methods, are summarized in Supplementary Table S16.

## CODE AVAILABILITY

The source code for this work is available at https://github.com/MAGIC-AI4Med/SPIDER. The SPIDER-KG web platform is hosted at https://spiderkg.internal.onexai.com.cn/.

## ACKNOWLEDGMENTS

This work is supported by the Innovative Drug Research and Development National Science and Technology Major Project (No. 2025ZD1803101), the Scientific Research Innovation Capability Support Project for Young Faculty (ZYGXQNJSKYCXNLZCXM-I22), and the Science and Technology Innovation Action Plan of Shanghai Municipality (No. 24QA2703800).

## AUTHOR CONTRIBUTIONS

Conceptualization, Xiaoman Zhang and Weidi Xie; methodology, Mingfei Liu, Weike Zhao, Xi-aoman Zhang, and Weidi Xie; software, Mingfei Liu and Weike Zhao; investigation, Mingfei Liu, Weike Zhao, Jingyi Chen, Xingran Quan, Xinyi Wang, Jinghao Feng, and Jiaxuan Han; data curation, Jingyi Chen, Xingran Quan, Xinyi Wang, and Jinghao Feng; formal analysis, Mingfei Liu and Weike Zhao; writing–original draft, Mingfei Liu; writing–review & editing, Mingfei Liu, Weike Zhao, Xiao Zhou, Yanfeng Wang, Xiaoman Zhang, and Weidi Xie; visualization, Mingfei Liu; supervision, Ya Zhang, Yanfeng Wang, Xiaoman Zhang, and Weidi Xie; project administration, Ya Zhang, Yanfeng Wang, Xiaoman Zhang, and Weidi Xie.

## REFERENCES

1. Karczewski, K.J., and Snyder, M.P. (2018). Integrative omics for health and disease. Nature Reviews Genetics 19, 299–310. doi: 10.1038/nrg.2018.4.

2. Chandak, P., Huang, K., and Zitnik, M. (2023). Building a knowledge graph to enable precision medicine. Scientific Data 10, 67. URL: https://doi.org/10.1038/s41597-023-01960-3. doi: 10.1038/s41597-023-01960-3.

3. Rosemblat, G., Fiszman, M., Shin, D., and Kilicoglu, H. (2019). Towards a characterization of apparent contradictions in the biomedical literature using context analysis. Journal of Biomedical Informatics 98, 103275. URL: https://doi.org/10.1016/j.jbi.2019.103275. doi: 10.1016/j.jbi.2019.103275.

4. Sosa, D.N., and Altman, R.B. (2022). Contexts and contradictions: a roadmap for computational drug repurposing with knowledge inference. Briefings in Bioinformatics 23, bbac268. URL: https://doi.org/10.1093/bib/bbac268. doi: 10.1093/bib/bbac268.

5. Nicholson, D.N., and Greene, C.S. (2020). Constructing knowledge graphs and their biomedical applications. Computational and Structural Biotechnology Journal 18, 1414–1428. URL: https://doi.org/10.1016/j.csbj.2020.05.017. doi: 10.1016/j.csbj.2020.05.017.

6. Zhang, Y., Sui, X., Pan, F., Yu, K., Li, K., Tian, S., Erdengasileng, A., Han, Q., Wang, W., Wang, J., Wang, J., Sun, D., Chung, H., Zhou, J., Zhou, E., Lee, B., Zhang, P., Qiu, X., Zhao, T., and Zhang, J. (2025). A comprehensive large-scale biomedical knowledge graph for AI-powered data-driven biomedical research. Nature Machine Intelligence 7, 602–614. URL: https://doi.org/10.1038/s42256-025-01014-w . doi:10.1038/s42256-025-01014-w.

7. Morris, J.H., Soman, K., Akbas, R.E., Zhou, X., Smith, B., Meng, E.C., Huang, C.C., Cerono, G., Schenk, G., Rizk-Jackson, A., Harroud, A., Sanders, L., Costes, S.V., Bharat, K., Chakraborty, A., Pico, A.R., Mardirossian, T., Keiser, M., Tang, A., Hardi, J., Shi, Y., Musen, M., Israni, S., Huang, S., Rose, P.W., Nelson, C.A., and Baranzini, S.E. (2023). The scalable precision medicine open knowledge engine (SPOKE): a massive knowledge graph of biomedical information. Bioinformatics 39, btad080. URL: https://doi.org/10.1093/bioinformatics/btad080. doi: 10.1093/bioinformatics/btad080.

8. Unni, D.R., Moxon, S.A.T., Bada, M., Brush, M., Bruskiewich, R., Caufield, J.H., Clemons, P.A., Dancik, V., Dumontier, M., Fecho, K., Glusman, G., Hadlock, J.J., Harris, N.L., Joshi, A., Putman, T., Qin, G., Ramsey, S.A., Shefchek, K.A., Solbrig, H., Soman, K., Thessen, A.E., Haendel, M.A., Bizon, C., and Mungall, C.J. (2022). Biolink Model: A universal schema for knowledge graphs in clinical, biomedical, and translational science. Clinical and Translational Science 15, 1848–1855. URL: https://doi.org/10.1111/cts.13302. doi: 10.1111/cts.13302.

9. Wood, E.C., Glen, A.K., Kvarfordt, L.G., Womack, F., Acevedo, L., Yoon, T.S., Ma, C., Flores, V., Sinha, M., Chodpathumwan, Y., Termehchy, A., Roach, J.C., Mendoza, L., Hoffman, A.S., Deutsch, E.W., Koslicki, D., and Ramsey, S.A. (2022). RTX-KG2: a system for building a semantically standardized knowledge graph for translational biomedicine. BMC Bioinformatics 23, 400. URL: https://doi.org/10.1186/s12859-022-04932-3. doi: 10.1186/s12859-022-04932-3.

10. Callahan, T.J., Tripodi, I.J., Stefanski, A.L., Cappelletti, L., Taneja, S.B., Wyrwa, J.M., Casiraghi, E., Matentzoglu, N.A., Reese, J., Silverstein, J.C., Hoyt, C.T., Boyce, R.D., Malec, S.A., Unni, D.R., Joachimiak, M.P., Robinson, P.N., Mungall, C.J., Cavalleri, E., Fontana, T., Valentini, G., Mesiti, M., Gillenwater, L.A., Santangelo, B., Vasilevsky, N.A., Hoehndorf, R., Bennett, T.D., Ryan, P.B., Hripcsak, G., Kahn, M.G., Bada, M., Baumgartner, W.A., and Hunter, L.E. (2024). An open source knowledge graph ecosystem for the life sciences. Scientific Data 11, 363. URL: https://doi.org/10.1038/s41597-024-03171-w. doi: 10.1038/s41597-024-03171-w.

11. Bonner, S., Barrett, I.P., Ye, C., Swiers, R., Engkvist, O., Bender, A., Hoyt, C.T., and Hamilton, W.L. (2022). A review of biomedical datasets relating to drug discovery: a knowledge graph perspective. Briefings in Bioinformatics 23, bbac404. URL: https://doi.org/10.1093/bib/bbac404. doi: 10.1093/bib/bbac404.

12. Alsentzer, E., Li, M.M., Kobren, S.N., Noori, A., Kohane, I.S., and Zitnik, M. (2025). Few shot learning for phenotype-driven diagnosis of patients with rare genetic diseases. npj Digital Medicine 8, 380. URL: https://doi.org/10.1038/s41746-025-01749-1. doi: 10.1038/s41746-025-01749-1.

13. Sun, J., Xu, C., Tang, L., Wang, S., Lin, C., Gong, Y., Ni, L., Shum, H., and Guo, J. (2024). Think-on-Graph: Deep and Responsible Reasoning of Large Language Model on Knowledge Graph. In The Twelfth International Conference on Learning Representations. URL: https://openreview.net/forum?id=nnVO1PvbTv.

14. Huang, K., Zhang, S., Wang, H., Qu, Y., Lu, Y., Roohani, Y., Li, R., Qiu, L., Li, G., Zhang, J., et al. (2025). Biomni: A general-purpose biomedical ai agent. biorxiv.

15. OpenAI (2026). Introducing GPT-Rosalind for life sciences research. OpenAI. URL: https://openai.com/index/introducing-gpt-rosalind/.

16. Anthropic (2026). Claude Science, an AI workbench for scientists, is now available. Anthropic. URL: https://www.anthropic.com/news/claude-science-ai-workbench.

17. Gottweis, J., Weng, W.H., Daryin, A. et al. (2026). Accelerating scientific discovery with Co-Scientist. Nature 655, 487–496. URL: https://doi.org/10.1038/s41586-026-10644-y. doi: 10.1038/s41586-026-10644-y.

18. Ghareeb, A.E., Chang, B., Mitchener, L., Yiu, A., Szostkiewicz, C.J., Shved, D., Gyimesi, G.J., Laurent, J.M., Wright, S.M., Razzak, M.T., White, A.D., Finnemann, S.C., Hinks, M.M., and Rodriques, S.G. (2026). A multi-agent system for automating scientific discovery. Nature 655, 497–505. URL: https://doi.org/10.1038/s41586-026-10652-y. doi: 10.1038/s41586-026-10652-y.

19. White, A.D., Braza, J.D., Pieler, M., Skarlinski, M., and Narayanan, S. (2026). Introducing PaperQA3: A Frontier Multimodal Deep Research Agent for Science. Edison Scientific. . URL: https://edisonscientific.com/articles/edison-literature-agent.

20. Ernst, P., Siu, A., and Weikum, G. (2015). Knowlife: a versatile approach for constructing a large knowledge graph for biomedical sciences. BMC bioinformatics 16, 157.

21. Piñero, J., Ramírez-Anguita, J.M., Saüch-Pitarch, J., Ronzano, F., Centeno, E., Sanz, F., and Furlong, L.I. (2020). The disgenet knowledge platform for disease genomics: 2019 update. Nucleic acids research 48, D845–D855.

22. Siu, P.C., Cabrera, C., Mudaliar, M., and Zubiaga, A. (2026). THBKG: A Temporal Biomedical Knowledge Graph for Decision-Aligned Clinical Advancement Prediction. . URL: https://arxiv.org/abs/2608.05982. doi: 10.48550/arXiv.2608.05982. arXiv:2608.05982.

23. Zheng, Y., Liu, W., Zeng, B., Feng, Y., Du, X., Zhou, L., and Li, Y. (2026). Automating biomedical knowledge graph construction for context-aware scientific inference. bioRxiv pp. 2026–01.

24. Sung, M., Jeong, M., Choi, Y., Kim, D., Lee, J., and Kang, J. (2022). BERN2: an advanced neural biomedical named entity recognition and normalization tool. Bioinformatics 38, 4837– 4839. URL: https://doi.org/10.1093/bioinformatics/btac598. doi: 10.1093/bioinformatics/b tac598.

25. Wei, C.H., Allot, A., Leaman, R., and Lu, Z. (2019). PubTator central: automated concept annotation for biomedical full text articles. Nucleic Acids Research 47, W587–W593. URL: https://doi.org/10.1093/nar/gkz389. doi: 10.1093/nar/gkz389.

26. Stepanov, I., Lukashov, O., Shtopko, M., and Kalyanarangan, V. (2026). Gliner-relex: A unified framework for joint named entity recognition and relation extraction. arXiv preprint arXiv:2605.10108.

27. Xu, A., Lin, B., Xue, B., Wang, B., Xu, B., Wu, B., Zhang, B., Lin, C., Dong, C., Ling, C., et al. (2026). Deepseek-v4: Towards highly efficient million-token context intelligence. arXiv preprint arXiv:2606.19348.

28. Escargueil-Blanc, I., Meilhac, O., Pieraggi, M.T., Arnal, J.F., Salvayre, R., and Nègre-Salvayre, A. (1997). Oxidized ldls induce massive apoptosis of cultured human endothelial cells through a calcium-dependent pathway: prevention by aurintricarboxylic acid. Arteriosclerosis, Thrombosis, and Vascular Biology 17, 331–339.

29. Lemay, J., Tea, B.S., Hamet, P., and deBlois, D. (2001). Regression of neointimal lesions in the carotid artery of nifedipine-treated shr and wky rats: possible role of apoptosis. Journal of vascular research 38, 462–470.

30. Șen, B., Ulusoy, E., Darcan, M., Ergün, M., Lobentanzer, S., Rifaioglu, A.S., Türei, D., Saez-Rodriguez, J., and Doğan, T. (2026). CROssBARv2: A unified computational framework for heterogeneous biomedical data representation and LLM-driven exploration. bioRxiv preprint. . URL: https://doi.org/10.64898/2026.04.12.718028. doi: 10.64898/2026.04.12.718028.

31. Zhang, D., Wang, Z., Li, Z.Z., Yu, Y., Jia, S., Dong, J., Xu, H., Wu, X., Zhang, Y., Zhang, T., Yang, J., Chen, X., and Song, L. (2026). Medkgent: a large language model agent framework for constructing temporally evolving medical knowledge graph. npj Digital Medicine. URL: https://doi.org/10.1038/s41746-026-03058-7 .doi: 10.1038/s41746-026-03058-7.

32. Yu, S., Yuan, Z., Xia, J., Luo, S., Ying, H., Zeng, S., Ren, J., Yuan, H., Zhao, Z., Lin, Y., Lu, K., Wang, J., Xie, Y., and Shum, H.Y. (2022). BIOS: An algorithmically generated biomedical knowledge graph. . URL: https://arxiv.org/abs/2203.09975. arXiv:2203.09975.

33. Soman, K., Rose, P.W., Morris, J.H., Akbas, R.E., Smith, B., Peetoom, B., Villouta-Reyes, C., Cerono, G., Shi, Y., Rizk-Jackson, A., Israni, S., Nelson, C.A., Huang, S., and Baranzini, S.E. (2024). Biomedical knowledge graph-optimized prompt generation for large language models. Bioinformatics 40, btae560. URL: https://doi.org/10.1093/bioinformatics/btae560. doi: 10.1093/bioinformatics/btae560.

34. Mountjoy, E., Schmidt, E.M., Carmona, M., Schwartzentruber, J., Peat, G., Miranda, A., Fumis, L., Hayhurst, J., Buniello, A., Karim, M.A., Wright, D., Hercules, A., Papa, E., Fauman, E.B., Barrett, J.C., Todd, J.A., Ochoa, D., Dunham, I., and Ghoussaini, M. (2021). An open approach to systematically prioritize causal variants and genes at all published human GWAS trait-associated loci. Nature Genetics 53, 1527–1533. URL: https://doi.org/10.1038/s41588-021-00945-5. doi: 10.1038/s41588-021-00945-5.

35. Griffith, M., Spies, N.C., Krysiak, K. et al. (2017). CIViC is a community knowledgebase for expert crowdsourcing the clinical interpretation of variants in cancer. Nature Genetics 49, 170–174. URL: https://doi.org/10.1038/ng.3774. doi: 10.1038/ng.3774.

36. Qu, Y., Li, T., Liu, Z., Li, D., and Tong, W. (2023). DICTrank: The largest reference list of 1318 human drugs ranked by risk of drug-induced cardiotoxicity using FDA labeling. Drug Discovery Today 28, 103770. URL: https://doi.org/10.1016/j.drudis.2023.103770. doi: 10.1016/j.drudis.2023.103770.

37. Zhang, Z., Huang, S., Jackson, A., Jones, E.A., Banka, S., Yang, C., Zhao, S., Lv, K., Peng, S., Lin, Z., and Wang, H. (2026). Biallelic GLTP mutations cause nonsyndromic epidermal differentiation disorder via disrupted epidermal glucosylceramide transport. The Journal of clinical investigation 136, e198835. URL: https://doi.org/10.1172/jci198835. doi: 10.1172/jci198835.

38. Backman, A.P.E., Halin, J., Nurmi, H., Möuts, A., Kjellberg, M.A., and Mattjus, P. (2018). Glucosylceramide acyl chain length is sensed by the glycolipid transfer protein. PloS one 13, e0209230. URL: https://doi.org/10.1371/journal.pone.0209230. doi: 10.1371/journal.pone .0209230.

39. Jiang, Y.J., Uchida, Y., Lu, B., Kim, P., Mao, C., Akiyama, M., Elias, P.M., Holleran, W.M., Grunfeld, C., and Feingold, K.R. (2009). Ceramide stimulates ABCA12 expression via peroxisome proliferator-activated receptor *δ* in human keratinocytes. The Journal of biological chemistry 284, 18942–18952. URL: https://doi.org/10.1074/jbc.m109.006973. doi: 10.1074/jbc.m109.006973.

40. Akiyama, M. (2014). The roles of ABCA12 in epidermal lipid barrier formation and keratinocyte differentiation. Biochimica et biophysica acta 1841, 435–440. URL: https://doi.org/10.1016/j.bbalip.2013.08.009. doi: 10.1016/j.bbalip.2013.08.009.

41. Nawaz, S., Tariq, M., Ahmad, I., Malik, N.A., Baig, S.M., Dahl, N., and Klar, J. (2012). Non-bullous congenital ichthyosiform erythroderma associated with homozygosity for a novel missense mutation in an atp binding domain of ABCA12. European journal of dermatology : EJD 22, 178–181. URL: https://doi.org/10.1684/ejd.2011.1638. doi: 10.1684/ejd.2011.1638.

42. Bastian, F.B., Cammarata, A.B., Carsanaro, S., Detering, H., Huang, W.T., Joye, S., Niknejad, A., Nyamari, M., Farias, T.M.d., Moretti, S., Tzivanopoulou, M., Wollbrett, J., and Robinson-Rechavi, M. (2024). Bgee in 2024: focus on curated single-cell RNA-seq datasets, and query tools. Nucleic Acids Research 53, D878–D885. URL: https://doi.org/10.1093/nar/gkae1118. doi: 10.1093/nar/gkae1118.

43. Liu, T., Hwang, L., Burley, S.K., Nitsche, C.I., Southan, C., Walters, W.P., and Gilson, M.K. (2024). BindingDB in 2024: a FAIR knowledgebase of protein-small molecule binding data. Nucleic Acids Research 53, D1633–D1644. URL: https://doi.org/10.1093/nar/gkae1075. doi: 10.1093/nar/gkae1075.

44. Oughtred, R., Rust, J., Chang, C., Breitkreutz, B., Stark, C., Willems, A., Boucher, L., Leung, G., Kolas, N., Zhang, F., Dolma, S., Coulombe-Huntington, J., Chatr-aryamontri, A., Dolinski, K., and Tyers, M. (2020). The BioGRID database: A comprehensive biomedical resource of curated protein, genetic, and chemical interactions. Protein Science 30, 187–200. URL: https://doi.org/10.1002/pro.3978. doi: 10.1002/pro.3978.

45. Degtyarenko, K., Matos, P.d., Ennis, M., Hastings, J., Zbinden, M., McNaught, A., Alcantara, R., Darsow, M., Guedj, M., and Ashburner, M. (2007). ChEBI: a database and ontology for chemical entities of biological interest. Nucleic Acids Research 36, D344–D350. URL: https://doi.org/10.1093/nar/gkm791. doi: 10.1093/nar/gkm791.

46. Zdrazil, B., Felix, E., Hunter, F., Manners, E.J., Blackshaw, J., Corbett, S., Veij, M.d., Ioannidis, H., Lopez, D.M., Mosquera, J.F., Magarinos, M.P., Bosc, N., Arcila, R., Kizilören, T., Gaulton, A., Bento, A.P., Adasme, M.F., Monecke, P., Landrum, G.A., and Leach, A.R. (2023). The ChEMBL Database in 2023: a drug discovery platform spanning multiple bioactivity data types and time periods. Nucleic Acids Research 52, D1180–D1192. URL: https://doi.org/10.1093/nar/gkad1004. doi: 10.1093/nar/gkad1004.

47. Diehl, A.D., Meehan, T.F., Bradford, Y.M., Brush, M.H., Dahdul, W.M., Dougall, D.S., He, Y., Osumi-Sutherland, D., Ruttenberg, A., Sarntivijai, S., Slyke, C.E.V., Vasilevsky, N.A., Haendel, M.A., Blake, J.A., and Mungall, C.J. (2016). The Cell Ontology 2016: enhanced content, modularization, and ontology interoperability. Journal of Biomedical Semantics 7, 44. URL: https://doi.org/10.1186/s13326-016-0088-7. doi: 10.1186/s13326-016-0088-7.

48. Davis, A.P., Wiegers, T.C., Johnson, R.J., Sciaky, D., Wiegers, J., and Mattingly, C.J. (2022). Comparative Toxicogenomics Database (CTD): update 2023. Nucleic Acids Research 51, D1257–D1262. URL: https://doi.org/10.1093/nar/gkac833. doi: 10.1093/nar/gkac833.

49. Basu, A., Bodycombe, N.E., Cheah, J.H., Price, E.V., Liu, K., Schaefer, G.I., Ebright, R.Y., Stewart, M.L., Ito, D., Wang, S., Bracha, A.L., Liefeld, T., Wawer, M., Gilbert, J.C., Wilson, A.J., Stransky, N., Kryukov, G.V., Dancik, V., Barretina, J., Garraway, L.A., Hon, C.S.Y., Munoz, B., Bittker, J.A., Stockwell, B.R., Khabele, D., Stern, A.M., Clemons, P.A., Shamji, A.F., and Schreiber, S.L. (2013). An Interactive Resource to Identify Cancer Genetic and Lineage Dependencies Targeted by Small Molecules. Cell 154, 1151–1161. URL: https://doi.org/10.1016/j.cell.2013.08.003. doi: 10.1016/j.cell.2013.08.003.

50. Cannon, M., Stevenson, J., Stahl, K., Basu, R., Coffman, A., Kiwala, S., McMichael, J.F., Kuzma, K., Morrissey, D., Cotto, K., Mardis, E.R., Griffith, O.L., Griffith, M., and Wagner, A.H. (2024). DGIdb 5.0: rebuilding the drug–gene interaction database for precision medicine and drug discovery platforms. Nucleic Acids Research 52, D1227–D1235. doi: 10.1093/nar/gk ad1040.

51. Baron, J.A., Johnson, C.M.S.B., Schor, M.A., Olley, D., Nickel, L., Felix, V., Bello, S.M., Greene, C., Lichenstein, R., Bisordi, K., Koka, R., Bearer, C., Macatangay, R., Ada, N., Ballenger, K., Bliss, E., Colliver, L., Dobbins, G., Heitzig, H., Dixon, S., Semesky, P., Garth, J., Fairchild, M., Gaskin, P., Zahid, S., Castillo, R., Edwards, S., Widjaja, A., Usui, Y., Lynch, E., Clarkson, M., Detwiler, T., and Schriml, L.M. (2025). The DO-KB knowledgebase 2026 update: expanding programmatic and language access. Nucleic Acids Research 54, D1425–D1436. URL: https://doi.org/10.1093/nar/gkaf1213. doi: 10.1093/nar/gkaf1213.

52. Knox, C., Wilson, M., Klinger, C.M., Franklin, M., Oler, E., Wilson, A., Pon, A., Cox, J., Chin, N.E.L., Strawbridge, S.A., Garcia-Patino, M., Kruger, R., Sivakumaran, A., Sanford, S., Doshi, R., Khetarpal, N., Fatokun, O., Doucet, D., Zubkowski, A., Rayat, D.Y., Jackson, H., Harford, K., Anjum, A., Zakir, M., Wang, F., Tian, S., Lee, B., Liigand, J., Peters, H., Wang, R.Q.R., Nguyen, T., So, D., Sharp, M., Silva, R.d., Gabriel, C., Scantlebury, J., Jasinski, M., Ackerman, D., Jewison, T., Sajed, T., Gautam, V., and Wishart, D.S. (2023). DrugBank 6.0: the DrugBank Knowledgebase for 2024. Nucleic Acids Research 52, D1265–D1275. URL: https://doi.org/10.1093/nar/gkad976. doi: 10.1093/nar/gkad976.

53. Avram, S., Wilson, T.B., Curpan, R., Halip, L., Borota, A., Bora, A., Bologa, C.G., Holmes, J., Knockel, J., Yang, J.J., and Oprea, T.I. (2022). DrugCentral 2023 extends human clinical data and integrates veterinary drugs. Nucleic Acids Research 51, D1276–D1287. URL: https://doi.org/10.1093/nar/gkac1085. doi: 10.1093/nar/gkac1085.

54. Gonzalez-Cavazos, A.C., Tanska, A., Mayers, M., Carvalho-Silva, D., Sridharan, B., Rewers, P.A., Sankarlal, U., Jagannathan, L., and Su, A.I. (2023). DrugMechDB: A Curated Database of Drug Mechanisms. Scientific Data 10, 632. URL: https://doi.org/10.1038/s41597-023-02534-z. doi: 10.1038/s41597-023-02534-z.

55. Malone, J., Holloway, E., Adamusiak, T., Kapushesky, M., Zheng, J., Kolesnikov, N., Zhukova, A., Brazma, A., and Parkinson, H. (2010). Modeling sample variables with an Experimental Factor Ontology. Bioinformatics 26, 1112–1118. URL: https://doi.org/10.1093/bioinformatics/btq099. doi: 10.1093/bioinformatics/btq099.

56. Aleksander, S.A., Balhoff, J.P., Carbon, S., Cherry, J.M., Ebert, D., Feuermann, M., Gaudet, P., Harris, N.L., Hill, D.P., Kalita, P., Lee, R., Mi, H., Moxon, S., Mungall, C.J., Muruganujan, A., Mushayahama, T., Sternberg, P.W., Thomas, P.D., Auken, K.V., Wong, E.D., Wood, V., Ramsey, J., Siegele, D.A., Chisholm, R.L., Dodson, R., Fey, P., Aspromonte, M.C., Nugnes, M.V., Naser, X.A.C., Tosatto, S.C.E., Giglio, M., Nadendla, S., Antonazzo, G., Attrill, H., Brown, N.H., Santos, G.d., Marygold, S., Röper, K., Strelets, V., Tabone, C.J., Thurmond, J., Zhou, P., Zaru, R., Lovering, R.C., Logie, C., Chen, D., Naba, A., Christie, K., Corbani, L., Ni, L., Sitnikov, D., Smith, C., Seager, J., Cooper, L., Elser, J., Jaiswal, P., Gupta, P., Naithani, S., Carme, P., Rutherford, K., Pons, J.L.D., Dwinell, M.R., Hayman, G.T., Kaldunski, M.L., Kwitek, A.E., Laulederkind, S.J.F., Tutaj, M.A., Vedi, M., Wang, S.J., D’Eustachio, P., Aimo, L., Axelsen, K., Bridge, A., Hyka-Nouspikel, N., Morgat, A., Goldbold, G., Engel, S.R., Miyasato, S.R., Nash, R.S., Sherlock, G., Weng, S., Bakker, E., Berardini, T.Z., Reiser, L., Auchincloss, A., Argoud-Puy, G., Blatter, M.C., Boutet, E., Breuza, L., Casals-Casas, C., Coudert, E., Estreicher, A., Famiglietti, M.L., Gos, A., Gruaz-Gumowski, N., Hulo, C., Jungo, F., Mercier, P.L., Lieberherr, D., et al. (2025). The Gene Ontology knowledgebase in 2026. Nucleic Acids Research 54, D1779–D1792. URL: https://doi.org/10.1093/nar/gkaf1292. doi: 10.1093/nar/gkaf1292.

57. Sollis, E., Mosaku, A., Abid, A., Buniello, A., Cerezo, M., Gil, L., Groza, T., Güneş, O., Hall, P., Hayhurst, J., Ibrahim, A., Ji, Y., John, S., Lewis, E., MacArthur, J.A.L., McMahon, A., Osumi-Sutherland, D., Panoutsopoulou, K., Pendlington, Z., Ramachandran, S., Stefancsik, R., Stewart, J., Whetzel, P., Wilson, R., Hindorff, L., Cunningham, F., Lambert, S.A., Inouye, M., Parkinson, H., and Harris, L.W. (2022). The NHGRI-EBI GWAS Catalog: knowledgebase and deposition resource. Nucleic Acids Research 51, D977–D985. URL: https://doi.org/10.1093/nar/gkac1010. doi: 10.1093/nar/gkac1010.

58. Seal, R.L., Gordon, S.M., Lush, M.J., Wright, M.W., and Bruford, E.A. (2010). genenames.org: the HGNC resources in 2011. Nucleic Acids Research 39, D514–D519. URL: https://doi.org/10.1093/nar/gkq892. doi: 10.1093/nar/gkq892.

59. Wishart, D.S., Guo, A., Oler, E., Wang, F., Anjum, A., Peters, H., Dizon, R., Sayeeda, Z., Tian, S., Lee, B.L., Berjanskii, M., Mah, R., Yamamoto, M., Jovel, J., Torres-Calzada, C., Hiebert-Giesbrecht, M., Lui, V.W., Varshavi, D., Varshavi, D., Allen, D., Arndt, D., Khetarpal, N., Sivakumaran, A., Harford, K., Sanford, S., Yee, K., Cao, X., Budinski, Z., Liigand, J., Zhang, L., Zheng, J., Mandal, R., Karu, N., Dambrova, M., Schiöth, H.B., Greiner, R., and Gautam, V. (2021). HMDB 5.0: the Human Metabolome Database for 2022. Nucleic Acids Research 50, D622–D631. URL: https://doi.org/10.1093/nar/gkab1062. doi: 10.1093/nar/gkab1062.

60. Thul, P.J., and Lindskog, C. (2017). The human protein atlas: A spatial map of the human proteome. Protein Science 27, 233–244. URL: https://doi.org/10.1002/pro.3307. doi: 10.1002/pro.3307.

61. Gargano, M.A., Matentzoglu, N., Coleman, B., Addo-Lartey, E.B., Anagnostopoulos, A.V., Anderton, J., Avillach, P., Bagley, A.M., Bakštein, E., Balhoff, J.P., Baynam, G., Bello, S.M., Berk, M., Bertram, H., Bishop, S., Blau, H., Bodenstein, D.F., Botas, P., Boztug, K., Čady, J., Callahan, T.J., Cameron, R., Carbon, S.J., Castellanos, F., Caufield, J.H., Chan, L.E., Chute, C.G., Cruz-Rojo, J., Dahan-Oliel, N., Davids, J.R., Dieuleveult, M.d., Souza, V.d., Vries, B.B.A.d., Vries, E.d., DePaulo, J.R., Derfalvi, B., Dhombres, F., Diaz-Byrd, C., Dingemans, A.J.M., Donadille, B., Duyzend, M., Elfeky, R., Essaid, S., Fabrizzi, C., Fico, G., Firth, H.V., Freudenberg-Hua, Y., Fullerton, J.M., Gabriel, D.L., Gilmour, K., Giordano, J., Goes, F.S., Moses, R.G., Green, I., Griese, M., Groza, T., Gu, W., Guthrie, J., Gyori, B., Hamosh, A., Hanauer, M., Hanušová, K., He, Y.O., Hegde, H., Helbig, I., Holasová, K., Hoyt, C.T., Huang, S., Hurwitz, E., Jacobsen, J.O.B., Jiang, X., Joseph, L., Keramatian, K., King, B., Knoflach, K., Koolen, D.A., Kraus, M.L., Kroll, C., Kusters, M., Ladewig, M.S., Lagorce, D., Lai, M.C., Lapunzina, P., Laraway, B., Lewis-Smith, D., Li, X., Lucano, C., Majd, M., Marazita, M.L., Martinez-Glez, V., McHenry, T.H., McInnis, M.G., McMurry, J.A., Mihulová, M., Millett, C.E., Mitchell, P.B., Moslerová, V., Narutomi, K., Nematollahi, S., et al. (2023). The Human Phenotype Ontology in 2024: phenotypes around the world. Nucleic Acids Research 52, D1333–D1346. URL: https://doi.org/10.1093/nar/gkad1005. doi: 10.1093/nar/gkad1005.

62. Zhang, J., Humphreys, I.R., Pei, J., Kim, J., Choi, C., Yuan, R., Durham, J., Liu, S., Choi, H.J., Baek, M., Baker, D., and Cong, Q. (2025). Predicting protein–protein interactions in the human proteome. Science 390, eadt1630. doi: 10.1126/science.adt1630.

63. Panneerselvam, K., Porras, P., del-Toro, N., Perfetto, L., Shrivastava, A., Ragueneau, E., Reyes, J.J.M., Orchard, S., and Hermjakob, H. (2024). IntAct Database for Accessing IMEx’s Contextual Metadata of Molecular Interactions. Current Protocols 4, e70018. URL: https://doi.org/10.1002/cpz1.70018. doi: 10.1002/cpz1.70018.

64. Kanehisa, M., Furumichi, M., Sato, Y., Kawashima, M., and Ishiguro-Watanabe, M. (2022). KEGG for taxonomy-based analysis of pathways and genomes. Nucleic Acids Research 51, D587–D592. URL: https://doi.org/10.1093/nar/gkac963. doi: 10.1093/nar/gkac963.

65. Wishart, D.S., Bartok, B., Oler, E., Liang, K.Y.H., Budinski, Z., Berjanskii, M., Guo, A., Cao, X., and Wilson, M. (2020). MarkerDB: an online database of molecular biomarkers. Nucleic Acids Research 49, D1259–D1267. URL: https://doi.org/10.1093/nar/gkaa1067. doi: 10.1093/nar/gkaa1067.

66. Mayo Foundation for Medical Education and Research (2026). Mayo Clinic Diseases and Conditions: official clinical resource. Mayo Clinic. . URL: https://www.mayoclinic.org/diseases-conditions.

67. Louden, D.N. (2020). MedGen: NCBI’s Portal to Information on Medical Conditions with a Genetic Component. Medical Reference Services Quarterly 39, 183–191. URL: https://doi.org/10.1080/02763869.2020.1726152. doi: 10.1080/02763869.2020.1726152.

68. U.S. National Library of Medicine (2026). Medical Subject Headings (MeSH): official documentation. National Library of Medicine. . URL: https://www.nlm.nih.gov/mesh/meshhome.html.

69. Vasilevsky, N.A., Toro, S., Matentzoglu, N., Flack, J.E., Mullen, K.R., Hegde, H., Gehrke, S., Whetzel, P.L., Shwetar, Y., Harris, N.L., Ngu, M.S., Alyea, G.L., Kane, M.S., Roncaglia, P., Sid, E., Thaxton, C.L., Wood, V., Abraham, R.S., Achatz, M.I., Ajuyah, P., Amberger, J.S., Babb, L., Baker, J., Balhoff, J.P., Berg, J.S., Bhalla, A., Ros, X.B.D., Braun, I.R., Broeren, E.C., Byer, B.K., Byrne, A.B., Callahan, T.J., Carmody, L.C., Chan, L.E., Clause, A.R., Cohen, J.S., DeLuca, M., Deuitch, N.T., Flowers, M., Fraser, J., Fujiwara, T., Gitau, V., Goldstein, J.L., Gration, D., Groza, T., Gyori, B.M., Hankey, W., Hilton, J.A., Himmelstein, D.S., Hong, S.S., Hoyt, C.T., Huether, R., Hurwitz, E., Jacobsen, J.O.B., Kikuchi, A., Köhler, S., Korn, D.R., Lagorce, D., Laraway, B.J., Li, J.Y., Malheiro, A.J., McLaughlin, J., Meldal, B.H.M., Mohan, S., Moxon, S.A.T., Munoz-Torres, M.C., Nelson, T.H., Nicholas, F.W., Ochoa, D., Olson, D., Oprea, T.I., Oskotsky, T.T., Osumi-Sutherland, D., Paris, K., Parkinson, H.E., Pendlington, Z.M., Peng, X.P., Pizzino, A., Plon, S.E., Powell, B.C., Ratliff, J.C., Rehm, H.L., Remennik, L., Riggs, E.R., Roberts, S., Robinson, P.N., Ross, J.E., Schaper, K., Schilder, B.M., Schmidt, J.L., Sharp, E.W., Similuk, M.N., Smedley, D., Sneddon, T.P., Sparks, R., Stefancsik, R., Stupp, G.S., Sundar, S., Takatsuki, T. et al. (2025). Mondo: integrating disease terminology across communities. GENETICS 232, iyaf215. URL: https://doi.org/10.1093/genetics/iyaf215. doi: 10.1093/genetics/iyaf215.

70. Maglott, D. (2004). Entrez Gene: gene-centered information at NCBI. Nucleic Acids Research 33, D54–D58. URL: https://doi.org/10.1093/nar/gki031. doi: 10.1093/nar/gki031.

71. Schoch, C.L., Ciufo, S., Domrachev, M., Hotton, C.L., Kannan, S., Khovanskaya, R., Leipe, D., Mcveigh, R., O’Neill, K., Robbertse, B., Sharma, S., Soussov, V., Sullivan, J.P., Sun, L., Turner, S., and Karsch-Mizrachi, I. (2020). NCBI Taxonomy: a comprehensive update on curation, resources and tools. Database 2020, baaa062. URL: https://doi.org/10.1093/database/baaa062. doi: 10.1093/database/baaa062.

72. Hamosh, A. (2004). Online Mendelian Inheritance in Man (OMIM), a knowledgebase of human genes and genetic disorders. Nucleic Acids Research 33, D514–D517. URL: https://doi.org/10.1093/nar/gki033. doi: 10.1093/nar/gki033.

73. Buniello, A., Suveges, D., Cruz-Castillo, C., Llinares, M.B., Cornu, H., Lopez, I., Tsukanov, K., Roldán-Romero, J.M., Mehta, C., Fumis, L., McNeill, G., Hayhurst, J.D., Osorio, R.E.M., Barkhordari, E., Ferrer, J., Carmona, M., Uniyal, P., Falaguera, M.J., Rusina, P., Smit, I., Schwartzentruber, J., Alegbe, T., Ho, V.W., Considine, D., Ge, X., Szyszkowski, S., Tsepilov, Y., Ghoussaini, M., Dunham, I., Hulcoop, D.G., McDonagh, E.M., and Ochoa, D. (2024). Open Targets Platform: facilitating therapeutic hypotheses building in drug discovery. Nucleic Acids Research 53, D1467–D1475. URL: https://doi.org/10.1093/nar/gkae1128. doi: 10.1093/nar/gkae1128.

74. Rath, A., Olry, A., Dhombres, F., Brandt, M.M., Urbero, B., and Ayme, S. (2012). Representation of rare diseases in health information systems: The orphanet approach to serve a wide range of end users. Human Mutation 33, 803–808. URL: https://doi.org/10.1002/humu.22078. doi: 10.1002/humu.22078

75. Rodchenkov, I., Babur, O., Luna, A., Aksoy, B.A., Wong, J.V., Fong, D., Franz, M., Siper, M.C., Cheung, M., Wrana, M., Mistry, H., Mosier, L., Dlin, J., Wen, Q., O’Callaghan, C., Li, W., Elder, G., Smith, P.T., Dallago, C., Cerami, E., Gross, B., Dogrusoz, U., Demir, E., Bader, G.D., and Sander, C. (2019). Pathway Commons 2019 Update: integration, analysis and exploration of pathway data. Nucleic Acids Research pp. 48(D1), D489–D497. URL: https://doi.org/10.1093/nar/gkz946. doi: 10.1093/nar/gkz946.

76. Barbarino, J.M., Whirl-Carrillo, M., Altman, R.B., and Klein, T.E. (2018). PharmGKB: A worldwide resource for pharmacogenomic information. WIREs Systems Biology and Medicine 10, e1417. URL: https://doi.org/10.1002/wsbm.1417. doi: 10.1002/wsbm.1417.

77. PhenomeBrowser Consortium (2021). PhenomeBrowser: official data-download resource. PhenomeBrowser. . URL: https://phenomebrowser.net/downloads/.

78. Natale, D.A., Arighi, C.N., Blake, J.A., Bona, J., Chen, C., Chen, S.C., Christie, K.R., Cowart, J., D’Eustachio, P., Diehl, A.D., Drabkin, H.J., Duncan, W.D., Huang, H., Ren, J., Ross, K., Ruttenberg, A., Shamovsky, V., Smith, B., Wang, Q., Zhang, J., El-Sayed, A., and Wu, C.H. (2016). Protein Ontology (PRO): enhancing and scaling up the representation of protein entities. Nucleic Acids Research 45, D339–D346. URL: https://doi.org/10.1093/nar/gkw1075. doi: 10.1093/nar/gkw1075.

79. Kim, S., Chen, J., Cheng, T., Gindulyte, A., He, J., He, S., Li, Q., Shoemaker, B.A., Thiessen, P.A., Yu, B., Zaslavsky, L., Zhang, J., and Bolton, E.E. (2022). PubChem 2023 update. Nucleic Acids Research 51, D1373–D1380. URL: https://doi.org/10.1093/nar/gkac956. doi: 10.1093/nar/gkac956.

80. Ragueneau, E., Gong, C., Sinquin, P., Sevilla, C., Beavers, D., Grentner, A., Griss, J., Hogue, G.F.J., Li, N.T., Matthews, L., May, B., Milacic, M., Mohammadi, H., Petryszak, R., Rothfels, K., Shamovsky, V., Stephan, R., Tiwari, K., Weiser, J., Wright, A., Gillespie, M., Wu, G., Stein, L., Hermjakob, H., and D’Eustachio, P. (2025). The Reactome Knowledgebase 2026. Nucleic Acids Research 54, D673–D681. URL: https://doi.org/10.1093/nar/gkaf1223. doi: 10.1093/nar/gkaf1223.

81. Kuhn, M., Letunic, I., Jensen, L.J., and Bork, P. (2015). The SIDER database of drugs and side effects. Nucleic Acids Research 44, D1075–D1079. URL: https://doi.org/10.1093/nar/gkv1075. doi: 10.1093/nar/gkv1075.

82. SNOMED International (2026). SNOMED CT documentation and release information. SNOMED International. . URL: https://www.snomed.org/value-of-snomed-ct.

83. Szklarczyk, D., Santos, A., Mering, C.v., Jensen, L.J., Bork, P., and Kuhn, M. (2015). STITCH 5: augmenting protein–chemical interaction networks with tissue and affinity data. Nucleic Acids Research 44, D380–D384. URL: https://doi.org/10.1093/nar/gkv1277. doi: 10.1093/nar/gkv1277.

84. Szklarczyk, D., Nastou, K., Koutrouli, M., Kirsch, R., Mehryary, F., Hachilif, R., Hu, D., Peluso, M.E., Huang, Q., Fang, T., Doncheva, N.T., Pyysalo, S., Bork, P., Jensen, L.J., and Mering, C.v. (2024). The STRING database in 2025: protein networks with directionality of regulation. Nucleic Acids Research 53, D730–D737. URL: https://doi.org/10.1093/nar/gkae1113. doi: 10.1093/nar/gkae1113.

85. Symptom Ontology contributors (2024). Symptom Ontology (SYMP): OBO Foundry resource. OBO Foundry. . URL: https://obofoundry.org/ontology/symp.html.

86. Zhou, Y., Zhang, Y., Zhao, D., Yu, X., Shen, X., Zhou, Y., Wang, S., Qiu, Y., Chen, Y., and Zhu, F. (2023). TTD: Therapeutic Target Database describing target druggability information. Nucleic Acids Research 52, D1465–D1477. URL: https://doi.org/10.1093/nar/gkad751. doi: 10.1093/nar/gkad751.

87. Mungall, C.J., Torniai, C., Gkoutos, G.V., Lewis, S.E., and Haendel, M.A. (2012). Uberon, an integrative multi-species anatomy ontology. Genome Biology 13, R5. URL: https://doi.org/10.1186/gb-2012-13-1-r5. doi: 10.1186/gb-2012-13-1-r5.

88. Bateman, A., Martin, M.J., Orchard, S., Magrane, M., Adesina, A., Ahmad, S., Bowler-Barnett, E.H., Bye-A-Jee, H., Carpentier, D., Denny, P., Fan, J., Garmiri, P., Gonzales, L.J.d.C., Hussein, A., Ignatchenko, A., Insana, G., Ishtiaq, R., Joshi, V., Jyothi, D., Kandasaamy, S., Lock, A., Luciani, A., Luo, J., Lussi, Y., Marin, J.S.M., Raposo, P., Rice, D.L., Santos, R., Speretta, E., Stephenson, J., Totoo, P., Tyagi, N., Urakova, N., Vasudev, P., Warner, K., Wijerathne, S., Yu, C.W.H., Zaru, R., Bridge, A.J., Aimo, L., Argoud-Puy, G., Auchincloss, A.H., Axelsen, K.B., Bansal, P., Baratin, D., Neto, T.M.B., Blatter, M.C., Bolleman, J.T., Boutet, E., Breuza, L., Gil, B.C., Casals-Casas, C., Echioukh, K.C., Coudert, E., Cuche, B., Castro, E.d., Estreicher, A., Famiglietti, M.L., Feuermann, M., Gasteiger, E., Gaudet, P., Gehant, S., Gerritsen, V., Gos, A., Gruaz, N., Hulo, C., Hyka-Nouspikel, N., Jungo, F., Kerhornou, A., Mercier, P.L., Lieberherr, D., Masson, P., Morgat, A., Paesano, S., Pedruzzi, I., Pilbout, S., Pourcel, L., Poux, S., Pozzato, M., Pruess, M., Redaschi, N., Rivoire, C., Sigrist, C.J.A., Sonesson, K., Sundaram, S., Sveshnikova, A., Wu, C.H., Arighi, C.N., Chen, C., Chen, Y., Huang, H., Laiho, K., Lehvaslaiho, M., McGarvey, P., Natale, D.A., Ross, K., Vinayaka, C.R., Wang, Y., and Zhang, J. (2024). UniProt: the Universal Protein Knowledgebase in 2025. Nucleic Acids Research 53, D609–D617. URL: https://doi.org/10.1093/nar/gkae1010. doi: 10.1093/nar/gkae1010.

89. Agrawal, A., Balcı, H., Hanspers, K., Coort, S.L., Martens, M., Slenter, D.N., Ehrhart, F., Digles, D., Waagmeester, A., Wassink, I., Abbassi-Daloii, T., Lopes, E.N., Iyer, A., Acosta, J.M., Willighagen, L.G., Nishida, K., Riutta, A., Basaric, H., Evelo, C.T., Willighagen, E.L., Kutmon, M., and Pico, A.R. (2023). WikiPathways 2024: next generation pathway database. Nucleic Acids Research 52, D679–D689. URL: https://doi.org/10.1093/nar/gkad960. doi: 10.1093/nar/gkad960.

90. Remy, F., Demuynck, K., and Demeester, T. (2024). Biolord-2023: semantic textual representations fusing large language models and clinical knowledge graph insights. Journal of the American Medical Informatics Association 31, 1844–1855.

91. Liu, A., Mei, A., Lin, B., Xue, B., Wang, B., Xu, B., Wu, B., Zhang, B., Lin, C., Dong, C., et al. (2025). Deepseek-v3.2: Pushing the frontier of open large language models. arXiv preprintarXiv:2512.02556.

92. Google DeepMind (2026). Gemini 3.1 Flash-Lite—Model Card. https://deepmind.google/models/model-cards/gemini-3-1-flash-lite/. .

93. Anthropic (2026). Claude Sonnet 4.6 System Card. https://www-cdn.anthropic.com/78073f739564e986ff3e28522761a7a0b4484f84.pdf. .

94. OpenAI (2026). GPT-5.5 System Card. https://deploymentsafety.openai.com/gpt-5-5/gpt-5-5.pdf. .

95. xAI (2025). Grok 4.1 Fast and Agent Tools API. . URL: https://x.ai/news/grok-4-1-fast.

96. MiniMax (2026). The MiniMax-M2 Series: Mini Activations Unleashing Max Real-World Intelligence. arXiv preprint arXiv:2605.26494. doi: 10.48550/arXiv.2605.26494.1

