## supplementary files for "SPIDER: Contextualizing Biomedical Knowledge for Evidence-Grounded Scientific Reasoning"

### S1 SUPPLEMENTARY CASE STUDY

**Illustrative examples of relation-level conflict classification.** Each entity pair spans two evidence rows. Single arrows give the subject-to-object orientation of the non-null relation; ↔ denotes an undirected association or interaction, not reciprocal causation. Null rows test the same relationship.

**Table S1.** Selected true-conflict examples. ADPR, ADP-ribose; 2-APB, 2-aminoethoxydiphenyl borate; CRSwNP, chronic rhinosinusitis with nasal polyps; hCG, human chorionic gonadotropin; SARA, SMAD anchor for receptor activation.

| Entity pair | Relation | PMID | Condition |
|---|---|---|---|
| TRPM2 ← 2-aminoethoxy-diphenyl borate (2-APB) | INTERFERES_WITH | 18204483 | *Model*: HEK293 cells expressing TRPM2<br>*ADPR*: 500 $\mu$M<br>*Pipette*: ATP 5 mM; free $Ca^{2+}$ 200 nM<br>*2-APB*: 30 $\mu$M |
| | NOT_ASSOCIATED_WITH | 15806115 | *Model*: HEK293 cells expressing TRPM2<br>*ADPR*: 500 $\mu$M<br>*Pipette*: ATP 5 mM; free $Ca^{2+}$ 200 nM<br>*2-APB*: 75/150 $\mu$M |
| Chlorhexidine gluconate → gingivitis | ACTIVE_AGAINST | 23433693 | *Formulation*: 0.12% chlorhexidine rinse<br>*Co-intervention*: home oral care |
| | NOT_ASSOCIATED_WITH | 1524688 | *Formulation*: Peridex rinse<br>*Co-intervention*: Crest/Rembrandt dentifrice<br>*Duration*: 8 weeks |
| Cadmium → L-ascorbic acid | NEGATIVELY_REGULATES | 32741747 | *Species*: *Pontederia cordata*<br>*Tissue*: leaves<br>$Cd^{2+}$: 0.44 mM |
| | NOT_ASSOCIATED_WITH | 32691317 | *Species*: *Pontederia cordata*<br>*Tissue*: leaves<br>$Cd^{2+}$: 50/75 mg/L |
| Valproate → decreased circulating carnitine concentration | CAUSES | 18555168 | *Population*: children<br>*Treatment*: valproate<br>*Analyte*: serum free carnitine |
| | NOT_ASSOCIATED_WITH | 21196091 | *Population*: children, 1–15 years<br>*Treatment*: valproate monotherapy<br>*Duration*: $\geq$6 months<br>*Analyte*: serum free carnitine |
| Parkinson disease ↔ VPS13C | ASSOCIATED_WITH | 29622492 | *Variant*: rs2414739<br>*Population*: Han Chinese |
| | NOT_ASSOCIATED_WITH | 26804608 | *Variant*: rs2414739<br>*Population*: Taiwanese |
| HLA-B27 antigen ← reactive arthritis | DEPENDS_ON | 2532780 | *HIV status*: positive<br>*Disease*: reactive arthritis |
| | NOT_ASSOCIATED_WITH | 8832993 | *HIV status*: positive<br>*Disease*: reactive arthritis |
| Allergic rhinitis → nasal polyps | CONTRIBUTES_TO | 10476318 | *Rhinitis control*: poorly controlled<br>*Outcome*: nasal-polyp recurrence |
| | NOT_ASSOCIATED_WITH | 33737534 | *Disease*: CRSwNP<br>*Follow-up*: 1 year after sinus surgery<br>*Outcome*: nasal-polyp recurrence |
| Preeclampsia ↔ chorionic gonadotropin (hCG) | ASSOCIATED_WITH | 14526312 | *Population*: pregnant women<br>*Analyte*: maternal serum hCG<br>*hCG level*: unexplained elevation |
| | NOT_ASSOCIATED_WITH | 27648965 | *Population*: unselected pregnant women<br>*Analyte*: maternal hCG |
| ZFYVE9 (SARA) ↔ TGF-$\beta$ receptor type I | INTERACTS_WITH | 23222715 | *Pathway*: TGF-$\beta$/SMAD2/3 signaling<br>*Receptor state*: activated |

Continued on next page

Table S1 (continued)

| Entity pair | Relation | PMID | Condition |
|---|---|---|---|
| | NOT_ASSOCIATED_WITH | 22819827 | *Pathway*: TGF-$\beta$/SMAD signaling<br>*Assay*: co-immunoprecipitation |

**Table S2.** Selected context-dependent differences. LPS, lipopolysaccharide; OGD, oxygen–glucose deprivation; $LD_{50}$, median lethal dose.

| Entity pair | Relation | PMID | Condition |
|---|---|---|---|
| AZD3965 $\rightarrow$ glycolysis | NEGATIVELY_REGULATES | 33762304 | *Model*: MYCN-amplified neuroblastoma<br>*Treatment*: AZD3965 + phenformin |
| | POSITIVELY_REGULATES | 25281618 | *Model*: human tumor cells/xenografts<br>*Treatment*: AZD3965 alone |
| MIR155 (miR-155) $\rightarrow$ tumor necrosis factor-alpha | NEGATIVELY_REGULATES | 26265120 | *Model*: macrophages<br>*Stimulus*: LPS |
| | POSITIVELY_REGULATES | 25811992 | *Model*: mouse brain/BV2 microglia<br>*Stimulus*: ischemia/OGD |
| Abiraterone $\rightarrow$ dihydrotestosterone (DHT) | NEGATIVELY_REGULATES | 27561382 | *Cell line*: LNCaP<br>*Substrate*: 3$\beta$-diol<br>*Medium*: 10% charcoal/dextran-stripped serum |
| | POSITIVELY_REGULATES | 28890368 | *Model*: androgen-responsive prostate cancer cells<br>*Substrate*: pregnenolone<br>*Culture condition*: serum starvation |
| Erythema marginatum $\rightarrow$ hereditary angioedema | NOT_ASSOCIATED_WITH | 17976427 | *C1 inhibitor*: normal |
| | PRECEDES | 32078942 | *C1 inhibitor*: deficient<br>*Disease event*: edema attacks |
| Infantile hypertrophic pyloric stenosis $\leftrightarrow$ erythromycins | ASSOCIATED_WITH | 30567916 | *Exposure window*: first 2 weeks after birth |
| | NOT_ASSOCIATED_WITH | 12381214 | *Exposure window*: prenatal |
| Doxorubicin $\rightarrow$ NANOG | NEGATIVELY_REGULATES | 24944491 | *Cell lines*: MHCC97-L/HCCLM3<br>*Dose*: $LD_{50}$<br>*Exposure duration*: $\leq$4 h |
| | POSITIVELY_REGULATES | 28030837 | *Cell line*: HCT116<br>*Exposure*: repeated cycles<br>*Cell state*: therapy-induced senescence |
| Isoprenaline $\rightarrow$ calcium-transporting ATPases | NEGATIVELY_REGULATES | 12776352 | *Species*: rat<br>*Compartment*: myocardial mitochondria<br>*Context*: isoproterenol-induced injury |
| | POSITIVELY_REGULATES | 2953502 | *Species*: rat<br>*Compartment*: sarcoplasmic reticulum<br>*Context*: $\beta$-adrenergic stimulation |
| Saxagliptin $\rightarrow$ cytochrome c | NEGATIVELY_REGULATES | 25446674 | *Model*: rotenone-treated rat<br>*Tissue*: striatum<br>*Treatment*: saxagliptin |
| | POSITIVELY_REGULATES | 25264749 | *Model*: diabetic Goto-Kakizaki rat<br>*Tissue*: aorta<br>*Treatment*: saxagliptin + exercise |
| Arachidonic acid $\rightarrow$ superoxide | CAUSES | 16902399 | *Model*: human HLE-B3 lens epithelial cells<br>*Culture condition*: serum starvation<br>*Arachidonic acid*: 30–150 $\mu$M |
| | NOT_ASSOCIATED_WITH | 9023282 | *Model*: rat heart slices/perfused hearts |

**Table S3. Newly reported gene–disease relationships and selected evidence-linked KG paths for 22 cases.** Each row represents one case; multiple selected paths for the same entity pair are shown on separate lines within the row. Arrows show KG relation direction; one selected relation type and its source identifier appear above and below each edge, respectively. Paths provide context for mechanistic hypotheses. GlcCer, glucosylceramide. Cadherins denotes a family node; cited evidence concerns E-cadherin. HP:0004389 is an HPO association without a PMID.

| New study (PMID) | KG path: relation above; source PMID below |
|---|---|
| 42147793 | APOA4 —NEGATIVELY_REGULATES (33276196)→ Interleukin-6 —ASSOCIATED_WITH (27789465)— osteoarthritis |
| 42403019 | ATP5ME —ASSOCIATED_WITH (26344197)— ATP5F1D —ASSOCIATED_WITH (32877691)— TMEM70 —CAUSES (28173120)→ mitochondrial disease |
| 42114970 | CAMSAP1 —BINDS_TO (24706919)→ Microtubules —ASSOCIATED_WITH (23589492)— HOOK1 —ASSOCIATED_WITH (12075009)— male infertility |
| 41981912 | CHCHD4 —ASSOCIATED_WITH (29568061)— AIFM1 —ASSOCIATED_WITH (29605508)— mitochondrial disease |
| 41796654 | CNN2 —AFFECTS (23576568)→ Actin Cytoskeleton ←MODIFIES (11706047)— FLNA —ASSOCIATED_WITH (HP:0004389)— intestinal pseudo-obstruction |
| 42027873 | CNN3 —POSITIVELY_REGULATES (31798778)→ Extracellular Signal-Regulated MAP Kinases —OBSERVED_IN (28871467)→ vascular associated smooth muscle cell —PARTICIPATES_IN (24814649)→ atherosclerosis |
| 42589629 | COPS7B —ASSOCIATED_WITH (27833851)— CUL3 —OBSERVED_IN (23661552)→ glioblastoma<br>COPS7B —ASSOCIATED_WITH (27833851)— CUL3 —INTERACTS_WITH (31652307)→ SPHK1 —ASSOCIATED_WITH (19723667)— glioblastoma |
| 41838796 | CRLF1 protein —POSITIVELY_REGULATES (10966616)→ STAT3 Transcription Factor —OBSERVED_IN (25583328)→ Cardiomyocyte hypertrophy ←HAS_PHENOTYPE (33705529)— hypertrophic cardiomyopathy |
| 42139047 | DECR1 protein —ASSOCIATED_WITH (32427840)— Ferroptosis ←REGULATES (30921410)— Arachidonate 15-Lipoxygenase —OBSERVED_IN (8466102)→ asthma |
| 42560623 | DSC3 —POSITIVELY_REGULATES (30857973)→ Cadherins —ASSOCIATED_WITH (29237910)— endometrial carcinoma |
| 41933287 | DYRK2 ←NEGATIVELY_REGULATES (31515251)— Kruppel-Like Factor 4 —NEGATIVELY_REGULATES (25892014)→ renal fibrosis |
| 42415215 | FOXQ1 ←POSITIVELY_REGULATES (21285253)— TGF-beta 1 —CONTRIBUTES_TO (28760197)→ endometriosis<br>FOXQ1 —POSITIVELY_REGULATES (24005989)→ Chemokine CCL2 —ASSOCIATED_WITH (30453861)— endometriosis |
| 41642656 | GLTP —BINDS_TO (30550553)→ GlcCer ←DELIVERS (19429679)— ABCA12 —ASSOCIATED_WITH (22257947)— ichthyosis |

Continued on next page

Table S3 (continued)

| New study (PMID) | KG path: relation above; source PMID below |
|---|---|
| 41846428 | GPRC5B —REGULATES (31104767)→ matrix metalloproteinase-9 —ASSOCIATED_WITH (19020756)— osteosarcoma<br>GPRC5B —REGULATES (31104767)→ matrix metalloproteinase-9 ←POSITIVELY_REGULATES (33228783)— Chemokine CCL2 —ASSOCIATED_WITH (33228783)— osteosarcoma |
| 42203489 | HSPB7 —REGULATES (29078393)→ Actin Cytoskeleton —ASSOCIATED_WITH (28524745)— vascular associated smooth muscle cell —ASSOCIATED_WITH (26129868)— Marfan syndrome |
| 42327722 | MGST1 —BINDS_TO (33539732)→ ALOX5 —ASSOCIATED_WITH (30614796)— papillary thyroid carcinoma<br>MGST1 —NEGATIVELY_REGULATES (33539732)→ Ferroptosis ←NEGATIVELY_REGULATES (33017703)— NFE2L2 —OBSERVED_IN (32824922)→ papillary thyroid carcinoma |
| 41679664 | PDE7A —REGULATES (19379723)→ Tumor Necrosis Factor-alpha —CAUSES (15805183)→ pulmonary fibrosis |
| 41836349 | PIEZO1 —POSITIVELY_REGULATES (30562052)→ nitric oxide —AFFECTS (8788938)→ Myocytes, Smooth Muscle ←AFFECTS (33032339)— adenomyosis |
| 41881217 | PLOD2 —ASSOCIATED_WITH (12881513)— Collagen ←PRODUCES (7330584)— Fibroblasts —ASSOCIATED_WITH (6871095)— psoriasis |
| 42165997 | TMED2 —POSITIVELY_REGULATES (29212217)→ IGF1R —ASSOCIATED_WITH (27729281)— insulin-like growth factor 1 —ASSOCIATED_WITH (32977489)— osteosarcoma |
| 42400826 | XRN2 —NEGATIVELY_REGULATES (33626405)→ Cadherins —ASSOCIATED_WITH (17647062)— esophageal squamous cell carcinoma |
| 42206677 | ZNF469 —REGULATES (26560304)→ Fibrillar Collagens ←PRODUCES (26566727)— Myofibroblasts —ASSOCIATED_WITH (19877031)— pulmonary fibrosis |

### S2 SUPPLEMENTARY METHODS

#### S2.1 Relation Mapping

Relation labels from different knowledge graph sources were mapped using a two-stage procedure. First, rule-based matching was performed after lowercasing the labels, extracting text following colons, and replacing underscores and hyphens with spaces. Exact matches against canonical relation codes and predefined keywords were considered, together with simple morphological variants such as `-s` and `-ed`.

Relations that could not be matched by these rules were classified using an LLM. The model was provided with the initial 56 relation categories and their definitions. It could select an existing category or propose a new category when no existing category was suitable. After classification, two additional categories proposed by the model were incorporated into the schema, resulting in the final 58 relation types listed in Table S19. The resulting mapping was then applied to the corresponding triples.

**System prompt:** https://github.com/MAGIC-AI4Med/SPIDER/blob/d0fe117682bd7890a441d7f87c9bada41fcbe571/prompt/relation_mapping.system.txt

**User prompt:** https://github.com/MAGIC-AI4Med/SPIDER/blob/d0fe117682bd7890a441d7f87c9bada41fcbe571/prompt/relation_mapping.user.txt

#### S2.2 Entity consolidation and identifier assignment

**Source identifiers and candidate matching.** Entity consolidation used the identifiers and names retained in source records. Primary identifiers were the records' main IDs; alternative identifiers were additional IDs explicitly listed for the same entry. Cross-references linked records to entries in referenced databases or ontologies and did not necessarily indicate equivalence. Candidate pairs were generated when records contained the same normalized identifier, including its namespace (a shared identifier), or matching normalized names or synonyms.

Identifiers unsuitable for identity matching were excluded from merge support. These included publication identifiers, cross-references explicitly indicating related, broader or narrower concepts, and identifiers associated with multiple records within the same source and entity type. Full InChIKeys, which identify chemical structures, were exempt from the last exclusion.

**Matching criteria.** Candidate pairs required sufficient identity support and no conflicting attributes before they could proceed to clustering. Primary and alternative identifiers and explicitly equivalent cross-references generally provided strong support, whereas ordinary cross-references generally provided weak support. A pair qualified if it shared an identifier providing strong support in both records, had supporting type-specific attributes, or shared weak identifiers from at least two independent identifier families together with identical normalized source primary names. Here, identifier families were predefined groups of related identifier systems, and source primary names were the main names in the original records, not synonyms. Multiple identifiers within one family counted only once. Name agreement alone was insufficient.

Type-specific attributes could support or reject a match. For drug and metabolite records, chemical comparisons used full InChIKeys and standard InChI structure representations. Matching structures could provide support, whereas conflicting available structures prevented merging. Records were also kept separate if one described a polymer and the other a chemically discrete structure, or if both listed ChEBI identifiers but had none in common. For genes, matching primary names, chromosomal locations and locus types supported merging when attributes were available in both records; locus type was the source-provided classification of the gene locus. Conflicting recorded gene attributes or different recorded organisms for two pathways prevented merging.

**Cluster checks and output.** Eligible records were joined into entity clusters only after checks on all records in the proposed cluster. Each cluster represented one KG entity and had to remain within one permitted type group: disease–phenotype–symptom, drug–metabolite, anatomy–cell, biological process–pathway, or experimental factor–measurement. Other KG entity types were restricted to same-type clusters. Records marked current by their sources were kept separate from obsolete or withdrawn records. Chemical consistency was checked across all cluster members to prevent indirect merging of incompatible structures, and type combinations were checked again before output.

ENT identifiers provided a common reference for source records and relation endpoints associated with the same KG entity. Automatically formed clusters, including records not merged with others, received sequential identifiers consisting of `ENT:` followed by eight digits with leading zeros. These identifiers were unique within the build, did not replace source identifiers or encode biological meaning, and could change across rebuilds.

The final output contained 3,748,007 entities after consolidation and targeted manual corrections, including selected cluster merges, splits and removal of incorrect synonyms. Each entity retained a representative name, other unique names as synonyms, an available definition, compatible KG entity types and source annotations. A separate mapping table linked ENT identifiers to retained source identifiers by namespace. Available database and publication metadata were retained for downstream graph construction.

#### S2.3 Relation endpoint matching and aggregation

**Endpoint matching.** Endpoint matching assigned ENT identifiers to the head and tail entities in each source relation record. Lookup indexes linked retained source identifiers, names and synonyms to consolidated KG entities. Matching first used the complete normalized identifier, including its namespace, and tried the local identifier without the namespace only when no complete match was found. A unique identifier match was accepted directly. For multiple identifier matches, names were used to identify a unique entity matching both. If no identifier matched, the endpoint name was compared with indexed entity names and synonyms.

**Ambiguity and unmatched endpoints.** When identifier and name evidence did not uniquely identify an entity, the endpoint resolver assigned one candidate according to its selection rule. These assignments were included in aggregation, with remaining ambiguity recorded separately for head and tail endpoints. Endpoint types were not used as matching constraints.

Only records with both endpoints assigned ENT identifiers proceeded to aggregation. Records with an unmatched endpoint were stored separately, retaining their source information and any assigned endpoint. Ambiguous assignments were therefore distinguished from endpoints with no matching KG entity.

**Relation aggregation and retained information.** Relation aggregation combined records according to their resolved endpoints and standardized relation types. The relation types were obtained through the mapping described in Appendix S2.1. Records with the same head and tail ENT identifiers were grouped together; within each directed entity pair, each standardized relation type was retained once, while different types were retained together. Endpoint order was preserved at this step, reverse pairs remained separate, and reverse edges were not generated automatically. One aggregated record could therefore represent more than one standardized triple.

Each aggregated record retained source-file references and the number of contributing source records. Source-file references identified the input files contributing to that directed entity pair. Repeated occurrences of the same standardized relation type increased the source-record count without adding another relation type. Available source information was retained for downstream graph construction. Together with the consolidated entity data, these outputs provided the basis for the research-oriented subgraph described in Appendix S2.5.

#### S2.4 Post-extraction evidence prioritization

For each retained literature extraction record $i$, the evidence prioritization score was defined as

$$S_i = G_i \left(0.25C_i + 0.25A_i + 0.25T_i + 0.25J_i\right), \tag{5}$$

where $C_i$ denotes the extraction confidence, $A_i$ is a simplified publication-format adjustment, $T_i$ is the temporal relevance score, $J_i$ is the journal-quartile score, and $G_i$ is an integrity gate for retracted publications. All component scores were constrained to $[0, 1]$, and therefore $S_i \in [0, 1]$.

The `relevance` and `clarity` fields were excluded because they exhibited near-zero empirical variance in the extracted corpus and consequently provided little discriminatory information. The resulting score was used as an interpretable ranking measure rather than as a calibrated probability that an extracted relation was true.

**Extraction confidence.** The extraction component $C_i$ was read from the stored `confidence` field after score consolidation and local adjustment (Appendix S2.6.3). It received the same individual weight as each of the other three components. Bibliographic and publication-level information jointly contributed the remaining three quarters of the weighted sum. This design reduces reliance on the model-derived extraction confidence while retaining its ability to differentiate extraction results.

**Publication format.** Publication types were obtained from the PubMed metadata associated with each PMID. We did not construct a detailed evidence hierarchy among systematic reviews, clinical trials, observational studies, and general research articles, because the relative evidential value of these designs depends on the type of biomedical relation being evaluated. Instead, the publication component was used only to penalize publication formats that generally provide more limited primary evidence.

The simplified mapping is shown in Table S4.

**Table S4.** Simplified mapping from PubMed publication type to publication-format score.

| Publication type | $A_i$ |
|---|---|
| Recognized scholarly publication types not listed below (including journal articles, reviews, meta-analyses, clinical studies, and observational studies) | 1.00 |
| Case report | 0.80 |
| Editorial, letter, or comment | 0.60 |
| Missing or unrecognized publication type | 0.80 |

When multiple publication-type labels were assigned to the same PMID, the lowest applicable score was used. This ensures that a generic Journal Article label does not override a more specific weak format, such as Case Reports, Editorial, or Letter. Under this mapping, publication type can change the final triple score by at most $0.25 \times (1.00 - 0.60) = 0.10$.

**Temporal relevance.** Recent publications were assigned moderately higher temporal relevance, while older evidence retained a non-zero contribution. Let $y_i$ be the publication year, $Y = 2026$ the scoring reference year, and

$$\Delta y_i = \max(0, Y - y_i). \tag{6}$$

The temporal component was calculated using a ten-year half-life and a lower bound of $0.50$:

$$T_i = 0.50 + 0.50 \times 2^{-\Delta y_i/10}. \tag{7}$$

Accordingly, publications from the reference year, ten years earlier, and twenty years earlier received temporal scores of $1.00$, $0.75$, and $0.625$, respectively. Missing or invalid publication years were assigned $T_i = 0.50$.

**Journal quartile.** Journal information was linked primarily by normalized ISSN and secondarily by normalized journal title. We mapped SCImago 2025 subject-category quartiles to scores using the function $w$:

$$w(Q1) = 1.00, \qquad w(Q2) = 0.90, \qquad w(Q3) = 0.80, \qquad w(Q4) = 0.70. \tag{8}$$

Because a journal may be assigned to multiple subject categories, its journal component was calculated as the median of its category-specific quartile scores:

$$J_i = \mathrm{median}_{c \in \mathcal{C}_i}\, w(Q_{ic}), \tag{9}$$

where $\mathcal{C}_i$ denotes the set of SCImago subject categories assigned to the journal associated with record $i$, and $Q_{ic}$ denotes its quartile in category $c$. Journals that could not be matched to SCImago were assigned $J_i = 0.85$.

The journal component accounted for $25\%$ of the total score. Consequently, the maximum difference between Q1 and Q4 journals in the final triple score was

$$0.25 \times (1.00 - 0.70) = 0.075. \tag{10}$$

This limited contribution reflects that journal-level indicators are not direct measures of the validity of an individual article.

**Research-integrity gate.** To exclude invalidated evidence, the integrity gate was defined as

$$G_i = \begin{cases} 0, & \text{if the PMID was labeled Retracted Publication,} \\ 1, & \text{otherwise.} \end{cases} \tag{11}$$

Consequently, every triple extracted from a retracted publication received a score of zero irrespective of its other component scores.

#### S2.5 Construction of the research-oriented subgraph

We constructed a research-oriented subgraph from the integrated KG to define a set of biomedical entities and relations for literature-based provenance augmentation and subsequent evaluation. The integrated KG contained 3,748,007 entities and 12,752,997 KG triples, a substantial fraction of which represented concepts used primarily for taxonomic or hierarchical classification. These concepts are not typically reported as independent findings in the biomedical literature. We therefore applied a three-stage filtering procedure comprising semantic entity filtering, atomic relation normalization, and full-PubMed-supported entity filtering.

**Stage 1: semantic entity filtering.** The pipe-delimited `entity_type` field was split into atomic KG entity types. An entity was retained if at least one atomic type belonged to the predeclared set of KG entity types retained in the research-oriented subgraph

$$\mathcal{T}_{\text{research}} = \{\texttt{anatomy}, \texttt{cell}, \texttt{disease}, \texttt{drug}, \texttt{gene}, \\ \texttt{metabolite}, \texttt{pathway}, \texttt{phenotype}, \texttt{protein}, \texttt{symptom}\}. \tag{12}$$

This rule retains merged types such as `drug|metabolite` and `disease|phenotype`, because each contains at least one research-oriented component. Pure `taxon`, `biological_process`, `cellular_component`, `molecular_function`, `experimental_factor`, and `measurement` entities do not satisfy this rule and are excluded. A total of 990,394 entities passed Stage 1.

**Stage 2: atomic relation normalization.** The semicolon-delimited `map_relation` field was split into atomic relation labels. We removed the structural set

$$\mathcal{R}_{\text{structural}} = \{\texttt{IS\_A}, \texttt{HAS\_SUBTYPE}, \texttt{HAS\_PART}, \texttt{RESTRICTED\_TO}, \texttt{HAS\_ATTRIBUTE}, \texttt{LOCATED\_IN}\}. \tag{13}$$

For an aggregated relation record $b$, let $R(b) \subseteq \mathcal{R}$ denote its set of standardized relation labels. After excluding structural relations, the retained labels were

$$R_{\text{research}}(b) = R(b) \setminus \mathcal{R}_{\text{structural}}. \tag{14}$$

A record was retained if both endpoints passed Stage 1 and $R_{\text{research}}(b) \neq \varnothing$. Thus, a record containing `ASSOCIATED_WITH; IS_A` retained only `ASSOCIATED_WITH`, whereas a record containing only `IS_A; LOCATED_IN` was removed. This filtering retained 3,170,475 aggregated relation records.

**Stage 3: full-PubMed-supported entity filtering.** For every entity retained after Stage 1, we searched the titles and abstracts of 40,219,061 records in the locally downloaded full PubMed corpus using its preferred name and all available synonyms. This full-corpus screening step was used only to determine which entities were sufficiently represented in the literature; it was separate from the subsequent retrieval of the approximately 1.95 million articles processed by the provenance agent. Before matching, both entity labels and PubMed text were normalized using Unicode NFKC normalization and case folding. Non-alphanumeric characters were replaced with spaces, consecutive whitespace was collapsed, and matches were required to cover the complete normalized label with token boundaries. Therefore, differences in capitalization, punctuation, and spacing were tolerated, whereas partial-word and fuzzy matches were not accepted.

Before retrieval, entity names and synonyms were screened to reduce spurious PubMed matches. Normalized labels were excluded if they contained no letters, matched a predefined stopword, had fewer than three non-space characters (two for gene entities), or exceeded 120 characters including spaces or 12 whitespace-separated words. We also excluded labels beginning with “obsolete ” or “deprecated ” in their original forms (case-insensitive), and labels shared as names or synonyms by more than 20 distinct Stage 1 entities.

Entities were screened using predefined criteria based on name length, the number of Stage 1 entities sharing each normalized name, preferred-name status, gene-symbol capitalization, and the number of distinct matching PMIDs. These criteria allowed a single matching PubMed record for longer or less widely shared names meeting the specified conditions, while requiring at least two distinct PMIDs for other qualifying names. Matches were combined across each entity’s preferred name and synonyms, without counting the same PMID more than once. This strategy reduced false-positive retrievals while preserving entities supported by either highly specific terminology or reproducible literature evidence.

Among the 990,394 entities obtained after semantic-type filtering, 207,549 had at least one valid full-PubMed label match and 198,659 satisfied the high-confidence selection criterion. The remaining 791,735 entities did not provide sufficient literature evidence for the high-confidence subgraph and were excluded. A Stage 2 triple record was retained only when both endpoint entities belonged to the selected entity set. The resulting research-oriented subgraph contained 198,659 entities and 1,104,268 triple records, corresponding to 1,099,652 unique undirected entity pairs.

#### S2.6 Provenance agent system

The system comprised three specialized LLM agents for document-level entity identification and consolidation, evidence-grounded relation discovery and relation classification. It processed one PubMed abstract per run, with the PMID attached outside the model calls. All three agents used the shared configuration in Table S5; their roles and interfaces are summarized in Table S6.

**Table S5.** Shared model and inference configuration of the three provenance agents.

| Item | Setting |
|---|---|
| Model | `deepseek-v4-flash`, accessed through an OpenAI-compatible Chat Completions interface |
| Message format | One system message followed by one user message for every request |
| Decoding | Temperature $= 0.1$; `top_p`, frequency and presence penalties, and random seed were not set |
| Reasoning mode | DeepSeek thinking mode disabled |
| Token limits | No explicit output-token limit; the agent system received the complete original abstract |
| Sampling | One model and one sample per request |
| Output handling | JSON requested in the prompt, then extracted, parsed and validated locally; no API-level `response_format` or JSON-schema mode |
| Reliability | 300-s request timeout; at most three transport attempts per request, with 1-s and 2-s exponential-backoff delays before the second and third attempts |
| Batch execution | Up to 100 simultaneous LLM calls per process in the reported batch launcher; no enforced inter-request interval |

**Table S6.** Roles and interfaces of the three provenance agents.

| Agent | Information used | Decision and output | Empty-output handling |
|---|---|---|---|
| Entity Identification and Consolidation Agent | Complete abstract and the extraction entity-category schema | Identify biomedical mentions, assign one of the 9 extraction entity categories and consolidate abbreviations, synonyms and repeated surface forms into a document-level typed entity inventory | One additional round after 2 s, with a corrective instruction |
| Relation Discovery Agent | Complete abstract and document-level entity inventory | Identify potentially related entity pairs and their supporting passages, including evidence distributed across sentences; no canonical relation label is assigned | Up to two additional rounds, separated by 3 s |
| Relation Classification Agent | Candidate pairs and supporting passages; endpoint extraction entity categories; 58 canonical relation definitions; entity–relation compatibility constraints | Determine the canonical relation from evidence semantics, direction and polarity; reject unsupported or schema-incompatible candidates | Up to two additional rounds without a corrective instruction |

**Output checks and stopping rules.** Repeated assessment was triggered when an agent produced no usable output: no entity records could be constructed from the parsed response, no candidate pairs remained after relation-discovery checks, or no assigned relations remained after classification checks. The limits in Table S6 allowed at most two attempts for entity identification and three attempts each for relation discovery and classification, including the initial attempt. Assessment stopped once usable outputs were obtained or the corresponding limit was reached. If the limit was reached, the affected stage returned an empty result. Individual rejected candidates did not trigger further assessment when other valid outputs remained. These repetitions were confined to the same agent; rejected candidates were not returned to earlier agents.

##### S2.6.1 Entity Identification and Consolidation Agent

**System prompt:** https://github.com/MAGIC-AI4Med/SPIDER/blob/d0fe117682bd7890a441d7f87c9bada41fcbe571/prompt/entity_identification.system.txt

**User prompt:** https://github.com/MAGIC-AI4Med/SPIDER/blob/d0fe117682bd7890a441d7f87c9bada41fcbe571/prompt/entity_identification.user.txt

**Retry instruction:** https://github.com/MAGIC-AI4Med/SPIDER/blob/d0fe117682bd7890a441d7f87c9bada41fcbe571/prompt/entity_identification.retry.txt

###### S2.6.2 Relation Discovery Agent

**System prompt:** https://github.com/MAGIC-AI4Med/SPIDER/blob/d0fe117682bd7890a441d7f87c9bada41fcbe571/prompt/relation_discovery.system.txt

**User prompt:** https://github.com/MAGIC-AI4Med/SPIDER/blob/d0fe117682bd7890a441d7f87c9bada41fcbe571/prompt/relation_discovery.user.txt

###### S2.6.3 Relation Classification Agent

**System prompt:** https://github.com/MAGIC-AI4Med/SPIDER/blob/d0fe117682bd7890a441d7f87c9bada41fcbe571/prompt/relation_classification.system.txt

**User prompt:** https://github.com/MAGIC-AI4Med/SPIDER/blob/d0fe117682bd7890a441d7f87c9bada41fcbe571/prompt/relation_classification.user.txt

**Output validation and extraction acceptance.** Extraction acceptance combined textual support, structural validity and predefined scoring rules. The Relation Classification Agent assessed textual support. Local checks required candidate identifiers from relation discovery, non-empty and distinct endpoints matched to the entity inventory after case and whitespace normalization, and allowed relation labels. Both endpoints required valid extraction categories, with matching categories for `IS_A`, `HAS_SUBTYPE`, `SIMILAR_TO` and `DIFFERENT_FROM`. Directional category blacklists constrained `TREATS`, `PREVENTS`, `INDICATED_FOR`, `IMPROVES`, `DIAGNOSES`, `DETECTS`, `CONFIRMS`, `MEASURES` and `PERFORMED_VIA`.

Discovery- and classification-stage confidence inputs were validated before score construction. Supplied values had to be numerical, finite and within $[0, 1]$; invalid inputs caused prediction rejection. Only absent or explicitly null values received the predefined fallback of 0.5; valid zeros were retained during missing-value handling. The base confidence $C_i^{\mathrm{base}}$ was the larger of the discovery- and classification-stage scores, with discovery confidence averaged across contributing predictions. Evidence attribution and uncertainty entered the confidence score as

$$C_i = \mathrm{clip}_{[0,1]}\left(C_i^{\mathrm{base}} + 0.05\, a_i - 0.10\, b_i - \min\left(0.15,\ 0.05\, h_i\right)\right), \tag{15}$$

where $a_i$ and $b_i$ indicate any predefined author-conclusion and citation cues, respectively, and $h_i$ counts distinct predefined hedging entries. Cues were identified by case-insensitive substring matching in the evidence text. Conclusion and citation contributions were each counted once. Clipping applied to the resulting score; out-of-range raw inputs were rejected.

For structurally valid candidates, the endpoint-name-length score $L_i$ was 0.90 when both endpoint strings exceeded two characters after trimming leading and trailing whitespace, and 0.80 otherwise. The schema-validity score $R_i$ was fixed at 0.90.

Candidate consolidation used case- and whitespace-normalized keys. Duplicates were grouped by $(\mathrm{head}, \mathrm{relation}, \mathrm{tail})$, retaining the highest base confidence within each classification output and the highest $C_i$ across extraction outputs. Competing predictions were grouped by directed $(\mathrm{head}, \mathrm{tail})$ pairs within each abstract, without grouping by evidence span or experimental condition. Competing relation-label combinations were those defined in Methods "Identification of potentially incompatible relations". Selection used $C_i$ and predefined label-polarity rules. For comparisons involving `NOT_ASSOCIATED_WITH`, higher $C_i$ was preferred, with ties favoring `NOT_ASSOCIATED_WITH`. This consolidation was intended to reduce spurious competing predictions caused by misinterpreting background statements or experimental controls as study findings. It did not establish scientific contradiction and could also remove valid condition-specific relations reported within the same abstract.

Remaining candidates were retained only if they passed the validity checks and the extraction acceptance threshold:

$$F_i = 0.5C_i + 0.25L_i + 0.25R_i, \qquad F_i \geq 0.6, \tag{16}$$

equivalent to $C_i \geq 0.30$ for $L_i = 0.90$ and $C_i \geq 0.35$ for $L_i = 0.80$, given $R_i = 0.90$. The fallback allowed missing confidence inputs to proceed to scoring rather than imposing automatic rejection. These scores were heuristic quantities rather than calibrated probabilities.

Publication metadata contributed to the subsequent evidence-prioritization score $S_i$ (Appendix S2.4), which used the same $C_i$, rather than $F_i$, as its extraction component.

**Accepted extraction output.** Accepted outputs comprised extracted triples and their supporting literature evidence, recorded using the fields in Table S7 before alignment to KG entities. Supporting evidence records were retained separately, allowing multiple records to be associated with the same standardized triple after graph integration.

**Table S7.** Structured output for an accepted extracted triple and its supporting literature evidence before KG entity alignment.

| `Field` | Type | Meaning |
|---|---|---|
| `pmid` | string | Source PMID attached outside the model calls |
| `head` | string | Accepted head entity name |
| `head_type` | enum | Extraction entity category of the head |
| `relation` | enum | Accepted canonical relation label |
| `tail` | string | Accepted tail entity name |
| `tail_type` | enum | Extraction entity category of the tail |
| `evidence` | string | Supporting text span for the accepted triple in the source record |
| `confidence` | number | Extraction confidence after score consolidation and local adjustment |

#### S2.7 Condition-extraction module

The condition-extraction module extracted the experimental or contextual conditions associated with each literature evidence record and its extracted triple. It received only the mapped entity pair, the upstream-assigned relation and its definition, the original triple, the selected condition template, and the paper title and abstract. It did not reassess the relation label, entity mapping, or relation direction.

**Table S8.** Main configuration of the condition-extraction module.

| Item | Setting |
|---|---|
| Model | `deepseek-v4-flash`, accessed through an OpenAI-compatible Chat Completions interface |
| Input | Entity pair; assigned relation and definition; original extracted triple and its literature evidence record; condition template; paper title and abstract |
| Inference | Temperature $= 0.1$; `top_p`, frequency and presence penalties, and random seed were not set |
| Reliability | 120-s timeout per attempt; one initial request plus three retries |
| Output handling | JSON requested in the prompt, then parsed and validated locally; no API-level structured-output mode |

**Output fields.** In the runtime schema, `evidence_unit` identifies the evidence-linked triple being contextualized, and each returned `condition_set` corresponds to a condition instance. Linking the extracted condition instances to the corresponding triple and literature evidence record yields the condition-enriched evidence-linked triple $q_{i,j}^{\mathrm{cond}}$ defined in Equation 4.

**Condition templates.** The 18 condition templates, their selection criteria and complete condition-field definitions are provided in `condition_templates.json` in the SPIDER code repository. Templates are selected using the incompatible relation combination and the canonical undirected pair of endpoint extraction categories. For comparisons involving `NOT_ASSOCIATED_WITH`, selection instead uses the semantic family of the opposing relation.

**System prompt:** https://github.com/MAGIC-AI4Med/SPIDER/blob/d0fe117682bd7890a441d7f87c9bada41fcbe571/prompt/condition_extraction.system.txt
**User prompt:** https://github.com/MAGIC-AI4Med/SPIDER/blob/d0fe117682bd7890a441d7f87c9bada41fcbe571/prompt/condition_extraction.user.txt

#### S2.8 Condition-aware classification module

The condition-aware classification module classified each evidence-eligible comparison unit after all condition extractions had been assembled. It received both relation definitions, all condition-enriched evidence-linked triples and their condition-extraction statuses, the extracted condition instances, and all cross-relation condition-instance pairs. Raw titles and abstracts were not included at this stage. The pairs described information availability only; the classification module determined scientific equivalence and the importance of observed differences.

**Table S9.** Main configuration of the condition-aware classification module.

| Item | Setting |
| --- | --- |
| Model | `deepseek-v4-flash` in the full workflow |
| Input | Entity pair; two relation labels and definitions; required condition fields; all condition-enriched evidence-linked triples and condition instances for both relations; all cross-relation condition-instance pairs |
| Inference | Temperature $= 0.1$; `top_p`, frequency and presence penalties, and random seed were not set |
| Reliability | 120-s timeout per attempt; one initial request plus three retries |
| Output handling | JSON requested in the prompt, then parsed, normalized, and validated locally; no API-level structured-output mode |

**System prompt:** https://github.com/MAGIC-AI4Med/SPIDER/blob/d0fe117682bd7890a441d7f87c9bada41fcbe571/prompt/conflict_classification.system.txt

**User prompt:** https://github.com/MAGIC-AI4Med/SPIDER/blob/d0fe117682bd7890a441d7f87c9bada41fcbe571/prompt/conflict_classification.user.txt

#### S2.9 Entity Alignment and Relation Matching for Benchmark Evaluation

##### S2.9.1 Entity-Type Harmonization

To enable a consistent comparison across baseline models, all model-specific entity types were deterministically mapped to the following 9 benchmark extraction entity categories: ANATOMY,

CHEMICAL, GENE, ORGANISM, DISEASE, PROCESS, PROCEDURE, MEASUREMENT, DEVICE.

The complete model-specific mappings are provided in Tables S10–S12. Multiple original entity types mapped to the same benchmark extraction entity category are grouped in a single row. The mappings were applied directly from the original model types or extraction queries to the benchmark schema; no intermediate entity-type mapping was used.

Predicted entities that could not be assigned to any of the 9 benchmark extraction entity categories were excluded from the category-harmonized evaluation. A predicted triple was also excluded if either of its endpoints could not be mapped to a benchmark extraction entity category.

This many-to-one harmonization enables comparison among models with different native entity schemas and levels of granularity. As a consequence, some fine-grained distinctions in the original model outputs were not retained. For example, genes, proteins, DNA, RNA, mutations, genetic variants, chromosomes, and genomic regions were uniformly represented as `GENE` in the benchmark schema.

**Table S10.** Mapping of BERN2 entity types to the benchmark extraction entity-category schema.

| **BERN2 original entity type(s)** | **Benchmark extraction entity category** |
|---|---|
| `CELL_TYPE` | `ANATOMY` |
| `DRUG` | `CHEMICAL` |
| `DNA`, `GENE`, `RNA` | `GENE` |
| `SPECIES` | `ORGANISM` |
| `DISEASE` | `DISEASE` |

*Note.* BERN2 did not provide entity predictions corresponding to `PROCESS`, `PROCEDURE`, `MEASUREMENT`, or `DEVICE`. These categories were nevertheless retained in the common benchmark extraction entity-category schema, and BERN2 therefore produced no true-positive predictions for them.

**Table S11.** Mapping of GLiNER-Relex entity queries to the benchmark extraction entity-category schema.

| **GLiNER-Relex original entity query or queries** | **Benchmark extraction entity category** |
|---|---|
| `tissue`, `organ`, `cell`, `anatomical structure` | `ANATOMY` |
| `drug`, `chemical`, `compound`, `medication` | `CHEMICAL` |
| `gene`, `protein`, `enzyme` | `GENE` |
| `organism`, `species` | `ORGANISM` |
| `disease`, `disorder`, `symptom`, `medical condition` | `DISEASE` |
| `biological process`, `pathway` | `PROCESS` |
| `treatment`, `therapy`, `medical procedure` | `PROCEDURE` |
| `measurement`, `concentration`, `biomarker`, `clinical variable` | `MEASUREMENT` |
| `medical device`, `diagnostic instrument` | `DEVICE` |

*Note.* For GLiNER-Relex, the mapping was specified as part of the extraction configuration. Predictions outside the benchmark extraction entity-category schema were removed. Duplicate entity predictions within the same abstract were collapsed using the lower-cased entity surface form together with the mapped benchmark extraction entity category.

**Table S12.** Mapping of PubTator entity types to the benchmark extraction entity-category schema.

| PubTator original entity or semantic type(s) | Benchmark extraction entity category |
|---|---|
| Chemical, Drug | CHEMICAL |
| Gene, Protein, SNP, Genetic variant, Chromosome, Chromosome locus, Genomic region | GENE |
| Species | ORGANISM |
| Disease | DISEASE |

*Note.* Mutation-, variant-, chromosome-, and genomic-region-related entities were mapped directly to GENE according to their original entity or semantic types, consistent with the benchmark schema. No intermediate mapping or corrective override was applied. PubTator did not provide entity predictions corresponding to PROCEDURE, MEASUREMENT, DEVICE, ANATOMY or PROCESS; it therefore produced no true-positive predictions for these categories.

**Treatment of Unmapped and Duplicate Entities.** Entity predictions that did not correspond to any of the 9 benchmark extraction entity categories were removed before evaluation. This filtering was applied after model inference and before entity- and triple-level matching.

If a predicted triple contained an unmapped entity as either its subject or object, the entire triple prediction was removed. This ensured that all evaluated triple endpoints belonged to the common benchmark extraction entity-category schema.

For models capable of producing overlapping or duplicate entity predictions, duplicate entities within the same abstract were identified using the lower-cased entity surface form and mapped benchmark extraction entity category. Predictions with the same normalized surface form and benchmark extraction entity category were treated as the same entity for downstream evaluation.

**Implications of the Harmonization Procedure.** The harmonized schema was designed to support direct evaluation across models with heterogeneous native ontologies rather than to preserve every distinction available in each original model. The mapping therefore intentionally reduced the granularity of some entity types.

In particular:

- CELL_LINE and CELL_TYPE were evaluated as ANATOMY;
- genes, proteins, enzymes, DNA, and RNA were evaluated as GENE;
- mutations, SNPs, genetic variants, chromosomes, chromosome loci, and genomic regions were also evaluated as GENE;
- drugs, chemicals, compounds, and medications were evaluated as CHEMICAL;
- diseases, disorders, symptoms, and medical conditions were evaluated as DISEASE; and
- treatments, therapies, and medical procedures were evaluated as PROCEDURE.

Accordingly, the evaluation measured whether a model identified an entity under the corresponding benchmark-level category. It did not assess whether the model preserved distinctions among original subtypes that were merged into the same benchmark category.

Because the baseline models differed in their native entity-type coverage, not every model could generate predictions for all 9 benchmark extraction entity categories. Categories unsupported by a model were not removed from the benchmark. Instead, they were retained in the common evaluation schema, and the model produced no true-positive predictions for those categories. This procedure avoided adapting the benchmark schema separately to the coverage of each baseline model.

###### S2.9.2 Relation-label normalization

Relations were evaluated using the predefined benchmark vocabulary in Table S19. GLiNER-Relex predictions already used this vocabulary. PubTator predictions were mapped to benchmark labels according to Table S13. Relation labels were compared after case and whitespace normalization. BERN2 was excluded from relation evaluation because the baseline pipeline produced entity predictions only.

PubTator's `Positive_Correlation` and `Negative_Correlation` labels were both mapped to `ASSOCIATED_WITH`. This deliberately coarse mapping retained the presence of an association but did not evaluate its polarity. Neither label was automatically interpreted as regulation, and negative correlation was not interpreted as absence of association. Original relation labels were retained for auditing. Normalization preserved endpoint order and prediction multiplicity; no triples were removed or merged.

**Table S13.** PubTator relation-label mapping for compatible evaluation.

| PubTator label | Benchmark label |
|---|---|
| `Comparison` | `COMPARED_WITH` |
| `Association` | `ASSOCIATED_WITH` |
| `Cotreatment` | `COMBINED_WITH` |
| `Bind` | `BINDS_TO` |
| `Positive_Correlation` | `ASSOCIATED_WITH` |
| `Negative_Correlation` | `ASSOCIATED_WITH` |

*Note.* Mapping positive and negative correlation to association is a lossy vocabulary adaptation, not a claim of semantic equivalence. Correlation polarity is retained in the original-label audit field but is not used in compatible matching.

###### S2.9.3 Compatible triple matching

After relation-label normalization, candidate triples from the same abstract connecting the same aligned entity pair were evaluated using Table S14. Identical labels required consistent endpoint order for directed relations, whereas undirected relations allowed either order. The directionality of each benchmark label is specified in Table S19.

Different labels were accepted in either endpoint order unless they formed an applicable predefined incompatibility pattern. No additional synonymy, specificity, or inverse-relation requirements were imposed. Compatibility therefore indicates acceptance under the specified contradiction constraints rather than semantic equivalence or independent verification of textual support.

A predicted triple was counted as matched if at least one reference candidate was compatible, and a reference triple was counted as recovered if at least one predicted candidate was compatible. An incompatible candidate did not invalidate a match to another compatible candidate. No global one-to-one matching constraint was imposed.

**Table S14.** Rules for compatible triple matching after entity alignment. Relation labels refer to the normalized benchmark vocabulary.

**A. General matching rules**

| Relation labels | Relation directionality | Endpoint order | Compatible |
|---|---|---|---|
| Same | Undirected | Same or reversed | Yes |
| Same | Directed | Same | Yes |
| Same | Directed | Reversed | No |
| Different; no applicable conflict | Both undirected | Same or reversed | Yes |
| Different; no applicable conflict | One directed, one undirected | Same or reversed | Yes |
| Different; no applicable conflict | Both directed | Same | Yes |
| Different; no applicable conflict | Both directed | Reversed | Yes |
| Different; applicable conflict | Any | As specified in Panel B | No |

**B. Predefined incompatibility patterns**

| Relation-label pair | Same order | Reversed order |
|---|---|---|
| `NOT_ASSOCIATED_WITH` versus any other label | Incompatible | Incompatible |
| `SIMILAR_TO` versus `DIFFERENT_FROM` | Incompatible | Incompatible |
| `POSITIVELY_REGULATES` versus `NEGATIVELY_REGULATES` | Incompatible | Compatible |
| `TREATS` versus `CONTRAINDICATED_FOR` | Incompatible | Compatible |

*Notes.* Same order means that the predicted head and tail correspond to the reference head and tail, respectively; reversed order means that they correspond to the reference tail and head. If both orders are satisfied, as for self-loops, same-order incompatibility rules still apply. The label pairs in Panel B are interchangeable between prediction and reference. Acceptance in reversed order for the last two pairs means only that their same-order incompatibility condition is not satisfied.

#### S2.10 SPIDER-Bench: conflict-aware question-answering benchmark construction

**Evidence-to-question construction.** The 200 questions were derived from 118 conflict groups. For each group, we assembled both relation sides, their PMID-linked titles and abstracts, supporting passages and structured condition instances (Appendix S2.7). Each source-specific question was linked to a representative evidence record and a particular condition instance. For context-dependent differences, questions retained the conditions needed to distinguish the applicable finding. For true conflicts, questions used shared or scientifically comparable conditions and the reference conclusion “cannot be determined uniquely”, supported by opposing evidence. Missing conditions were not treated as evidence of equivalence. Questions specified the entities, measured outcome and necessary context without revealing the answer or relying on unshown source text. Four fixed conclusion-based options were matched to the question semantics; reference answers followed the source findings rather than the ordering of relation labels.

**Model-assisted generation.** LLM assisted question drafting; programmatic generation used `deepseek-v4-flash` through a Chat Completions interface, with temperature 0.1 and a maximum output of 4,000 tokens. Inputs comprised the entity and relation pair, both sides’ literature evidence, condition instances and fixed answer options. Outputs were JSON records containing conditions, answer modes, proposed reference answers and source identifiers.

**Validation, review and reproducibility.** Automated checks assessed source references, condition-instance indices, output completeness and compatibility between question wording and answer options. Biomedical expert review assessed evidential support, condition sufficiency or comparability, answer consistency and clarity using both sides' evidence. Unsupported conclusions, incorrect entity or relation interpretations, and unresolved ambiguity were not accepted. Final records retained conflict-group identifiers, source and condition links, reference answers and review information.

##### S2.10.1 Example questions

Table S15 presents two questions from the same nifedipine–apoptosis conflict group in the final benchmark. Their different study contexts identify which finding applies to the queried entity pair.

**Table S15.** Example condition-resolution questions from one conflict group.

| Item | Question | Reference answer | Source PMID |
|---|---|---|---|
| Q1 | How does nifedipine affect doxorubicin-induced apoptosis in cardiomyocytes? | **A.** Decreases or inhibits | 31285514 |
| Q2 | In spontaneously hypertensive rats, how does nifedipine affect apoptosis of carotid neointimal smooth-muscle cells? | **B.** Increases or activates | 11561148 |

**Shared answer options:** A, Decreases or inhibits; B, Increases or activates; C, No significant effect; D, Cannot be determined uniquely.

##### S2.10.2 Generation prompts

**Shared system prompt:** https://github.com/MAGIC-AI4Med/SPIDER/blob/d0fe117682bd7890a441d7f87c9bada41fcbe571/prompt/qa.system.txt

**Condition-resolution user prompt:** https://github.com/MAGIC-AI4Med/SPIDER/blob/d0fe117682bd7890a441d7f87c9bada41fcbe571/prompt/qa_condition_generation.user.txt

**True-conflict user template:** https://github.com/MAGIC-AI4Med/SPIDER/blob/d0fe117682bd7890a441d7f87c9bada41fcbe571/prompt/qa_true_conflict_generation.user.txt

#### S2.11 Question-answering metrics and statistical procedures

**Evaluation metrics.** Accuracy was calculated as the proportion of questions answered correctly. Macro-$F_1$ was the unweighted mean of the class-specific $F_1$ scores for the four displayed answer labels (A–D), rather than for the two question categories.

**Confidence intervals and paired comparisons.** The 118 conflict groups were used as the resampling units to preserve dependence between questions derived from the same relation comparison. For each of 10,000 bootstrap replicates, conflict groups were sampled with replacement, retaining all questions from each sampled group, and each metric was recalculated. The same sampled groups were used across settings. Pointwise 95% confidence intervals were obtained from the 2.5th and 97.5th percentiles of the bootstrap distributions; intervals for paired differences were calculated from the corresponding within-replicate metric differences.

Paired permutation tests used 10,000 permutations. Within each permutation, the outputs of the two compared settings were swapped or retained at the conflict-group level, with the same assignment applied to all questions in a group. The metric difference was recalculated after each permutation. Two-sided $P$ values were calculated as $(b+1)/(10{,}000+1)$, where $b$ was the number of permutations for which the absolute metric difference was at least as large as the observed absolute difference.

Holm correction was applied jointly across accuracy and macro-$F_1$ within each of two separate comparison families, rather than separately for each metric. The information-ablation family comprised eight tests comparing each of four configurations (triples alone, triples with literature evidence, triples with structured study conditions, and triples with both conditions and literature evidence) with LLM only on both metrics. The cross-KG family comprised ten tests comparing each of five baselines (CROssBARv2-KG, MedKGent-KG, RTX-KG2, BIOS v3, and PrimeKG) with SPIDER-KG on both metrics. Statistical significance was assessed at a Holm-adjusted $P < 0.05$. Bootstrap confidence intervals remained pointwise and were not adjusted for multiple comparisons.

#### S2.12 Downstream biomedical task evaluation

The four tasks assessed complementary biomedical applications: statement verification (BioMixQA), candidate causal-gene ranking at GWAS loci (Open Targets), publication-specific support for clinical interpretations (CIViC), and drug cardiotoxicity concern classification (DICTrank).

GWAS records were grouped by GRCh38 locus and trait, retaining all reference genes without adding those outside the candidate pool; a gene qualified if any GENCODE basic-transcript start site met the window criterion in Methods. CIViC retained accepted, unflagged predictive evidence at levels A–C, grouped by publication, clinical context and significance; identical inputs with conflicting or unknown support directions were excluded. DICTrank retained separate product records sharing an ingredient. Reference labels and private explanations were excluded from generation.

SPIDER-KG used identifier matching and BioLORD-based name retrieval, whereas external KGs used native identifiers, aliases and lexical retrieval. Relations, attributes and stored KG-associated literature passages were filtered for relevance and supplied with source identifiers, without live literature search. Final evidence was limited to 16,000 characters, or $\min(48{,}000, \max(16{,}000, 1{,}200m))$ for $m$ GWAS candidates. All final-answer requests used `deepseek-v4-flash` with temperature $= 0$ and `max_tokens` set to 1,500 for BioMixQA and CIViC and 2,048 for GWAS and DICTrank, identically across configurations within each task. Evidence-available subsets comprised questions whose final SPIDER-KG-supported prompt contained at least one selected relation, non-identity attribute or literature passage. For BioMixQA questions explicitly querying disease ontology identifiers, selected DOID mappings also counted as evidence. Names, aliases and identifiers used only for entity matching, citation identifiers and publication titles alone did not qualify. Selection did not use reference labels or answer correctness. This yielded 307 BioMixQA, 137 GWAS, 1,144 CIViC and 1,192/1,296 three-/four-class DICTrank questions, on which direct LLM and SPIDER-KG-supported answers were compared.

Full cohorts and evidence subsets used the metrics specified in Methods. For CIViC and BioMixQA accuracy, invalid responses and abstentions counted as incorrect. For DICTrank macro-F1, they counted as false negatives for the reference class, without introducing an additional class. GWAS rankings required every candidate identifier exactly once in a strict order. MRR averaged the reciprocal rank of the highest-ranked reference gene across tasks, assigning zero to abstentions, invalid rankings (including incomplete or tied rankings), and rankings with no reference-gene match.

For both full cohorts and evidence subsets, confidence intervals followed the cluster-bootstrap procedure in Appendix S2.11. For evidence-subset comparisons, Holm step-down correction was applied jointly to the five primary-metric comparisons: BioMixQA accuracy, GWAS MRR, CIViC accuracy, and DICTrank three- and four-class macro-F1, controlling the family-wise error rate at $\alpha = 0.05$. Resampling units were individual questions for BioMixQA, connected components of overlapping windows on the same chromosome for GWAS, shared-PMID components for CIViC, and ingredient/source-drug groups for DICTrank; subsets retained these assignments. Reported intervals were pointwise 95% confidence intervals for metric estimates, not paired differences, and were not adjusted for multiplicity.

#### S2.13 Calibration of similarity thresholds

##### S2.13.1 Predicted–reference entity matching

We randomly sampled 100 PubMed abstracts outside the 1,000-abstract extraction benchmark and divided them into 50 development abstracts and 50 validation abstracts. All candidates from an abstract remained in the same set. Biomedical annotators annotated reference entities in each abstract. We collected predictions from all evaluated systems and harmonized their entity categories (Appendix S2.9.1). Within each abstract and category, we paired each predicted entity with every reference entity. Annotators labeled whether each pair referred to the same entity. We computed BioLORD-2023 cosine similarity for each pair.

We tested thresholds from 0.50 to 1.00 in increments of 0.05. We classified a pair as a match when its similarity reached the threshold. We calculated matching precision, recall and F1 against the manual labels, pooling candidate-pair counts across systems and abstracts. We selected 0.70 by maximizing development-set F1 and fixed this threshold for validation and benchmark evaluation. With this threshold, the development-set precision, recall and F1 were 94.17%, 92.38% and 93.27%. The corresponding validation-set values were 93.64%, 91.76% and 92.69%.

##### S2.13.2 Entity-to-KG linking

For extracted entities without an exact match to normalized KG names or aliases, we retrieved the node with the highest BioLORD-2023 cosine similarity from the fixed research-oriented subgraph.

We selected the similarity threshold using a development set of 100 entity–candidate pairs sampled before threshold filtering and manually labeled as matches or non-matches by annotators blinded to similarity scores. Thresholds from 0.50 to 1.00 in increments of 0.05 were evaluated using balanced accuracy, defined as the mean of sensitivity and specificity. The selected threshold of 0.85 achieved a development-set balanced accuracy of 92.9% and precision of 94.9%.

After fixing the threshold, we evaluated the linking procedure on an independent random sample of 300 entity–candidate pairs drawn before threshold filtering. Test annotators were blinded to similarity scores and acceptance decisions. Evaluation yielded 167 true positives, 8 false positives, 13 false negatives and 112 true negatives. Precision was 95.43% (95% Wilson confidence interval, 91.24–97.67%), balanced accuracy was 93.06%, and the acceptance rate was 58.33% (175/300). Sensitivity for identifying correct candidate matches was 92.78%, specificity was 93.33%, and F1 was 94.08%.

### S3 SUPPLEMENTARY TABLES

**Table S16.** Primary data sources integrated into the biomedical knowledge graph. Download URLs identify source access points, and *Access* describes the retrieval method.

| Database | Version | Download URL | Access | Formats |
|---|---|---|---|---|
| Bgee | 15.2 | https://www.bgee.org/ftp/current/bgee_v15_2 | HTTPS (wget -r) | TSV, RDF/TTL, JSON-LD, SQL dump |
| BindingDB | 2025-10-01 | https://www.bindingdb.org/rwd/bind/downloads/ | HTTPS (wget) | TSV, SDF, MySQL/Oracle dump |
| BioGRID | 5.0.252 | https://downloads.thebiogrid.org/BioGRID/Latest-Release/ | HTTPS (wget) | TAB3, MITAB, PSI-MI XML |
| ChEBI | rel246 | https://ftp.ebi.ac.uk/pub/databases/chebi/archive/rel246/ | FTP / HTTPS | OBO, OWL, SDF, TSV, SQL |
| ChEMBL | 36 | https://ftp.ebi.ac.uk/pub/databases/chembl/ChEMBLdb/releases/chembl_36/ | HTTPS (wget -r) | SQLite / MySQL / PostgreSQL dumps, SDF, FASTA, H5, TXT |
| CL (Cell Ontology) | v2025-12-17 | http://purl.obolibrary.org/obo/cl.owl | HTTPS | OWL, OBO, JSON, TSV, SQLite |
| CTD | retrieved 2025-12-30 | https://ctdbase.org/reports/ | HTTPS (wget -r) | CSV, TSV, XML |
| CTRP | CTRPv2.0 (v20) | https://ctd2-data.nci.nih.gov/Public/Broad/CTRPv2.0_2015_ctd2_ExpandedDataset/ | HTTPS | TXT, XLSX, RDS |
| DGIdb | 2024-12 | https://www.dgidb.org/downloads | HTTPS | TSV |
| DOID | v2025-12-23 | http://purl.obolibrary.org/obo/doid.json | HTTPS | JSON, OBO, OWL |
| DrugBank | 5.1.13 | https://go.drugbank.com/releases/latest | Registered download | XML |
| DrugCentral | 2023-11-01 | https://drugcentral.org/download | HTTPS | SQL dump, TSV, SDF, SMILES |
| DrugMechDB | 2.0.1 | https://github.com/SuLab/DrugMechDB/releases | git / HTTPS | YAML, JSON |
| EFO | v3.85.0 | http://www.ebi.ac.uk/efo/efo.obo | HTTPS | OBO |
| GO | 2025-10-10 release | https://current.geneontology.org/ | HTTPS | OBO, OWL, JSON, GAF, GPAD, GPI |
| GWAS Catalog | 2025-12-23 | https://ftp.ebi.ac.uk/pub/databases/gwas/releases/latest/ | FTP | TSV, ZIP |
| HGNC | 2025-12-05 | https://www.genenames.org/download/ | HTTPS | TSV |
| HMDB | 5.0 | https://hmdb.ca/downloads | HTTPS | XML |
| HPA | v25.0 | https://www.proteinatlas.org/about/download | HTTPS | JSON, TSV |
| HPO | v2025-11-24 | http://purl.obolibrary.org/obo/hp.obo | HTTPS | OBO, OWL, JSON, HPOA, TSV |

Continued on next page

Table S16 – (continued)

| Database | Version | Download URL | Access | Formats |
|---|---|---|---|---|
| humanPPI | 2025-09-16 | https://conglab.swmed.edu/humanPPI/downloads/ | HTTPS | TSV / TAR.GZ |
| IntAct | 2025-08-08 | https://ftp.ebi.ac.uk/pub/databases/intact/current/ | FTP | PSI-MI TAB / XML |
| KEGG | 116.0 | https://www.kegg.jp/kegg/rest/ | KEGG REST API | TXT |
| MarkerDB | 2.0 | https://markerdb.ca/downloads | HTTPS | TSV |
| Mayo Clinic | Browser, Historical release unverified | https://www.mayoclinic.org/diseases-conditions | HTTPS (custom scraper) | JSON, CSV |
| MedGen | Rolling updates; historical release unverified | https://ftp.ncbi.nlm.nih.gov/pub/medgen/ | FTP | RRF, TSV |
| MeSH | 2025 | https://www.nlm.nih.gov/databases/download/mesh.html | HTTPS | XML |
| Mondo | v2025-12-02 | http://purl.obolibrary.org/obo/mondo.owl | HTTPS | OBO, OWL, JSON |
| NCBI Gene | Rolling updates; historical release unverified | https://ftp.ncbi.nlm.nih.gov/gene/DATA/ | FTP | TSV (gzipped) |
| NCBITaxon | v2025-12-03 | http://purl.obolibrary.org/obo/ncbitaxon.obo | HTTPS | OBO |
| OMIM | Rolling updates; historical release unverified | https://omim.org/downloads/ | Registered API | JSON, TSV |
| Open Targets | 25.12 | https://ftp.ebi.ac.uk/pub/databases/opentargets/platform/ | FTP / rsync | Parquet, JSON |
| Orphanet | 2025-12 | https://www.orphadata.com/ | HTTPS (custom scraper) | TSV, HTML, XML |
| Pathway Commons | v14 | https://www.pathwaycommons.org/archives/PC2/ | HTTPS | BioPAX, SIF, GMT |
| PharmGKB | Rolling updates; historical release unverified | https://api.clinpgx.org/v1/download/file/data/ | HTTPS | TSV, JSON, ZIP |
| Phenome Browser | Historical release unverified | https://phenomebrowser.net/downloads/ | HTTPS | TAR.GZ |
| PRO | release 72.0 | https://proconsortium.org/download/ | HTTPS | OBO, TXT |
| PubChem | Periodic exports; historical release unverified | https://ftp.ncbi.nlm.nih.gov/pubchem/ | FTP | SDF, CSV, XML, ASN.1, TTL, JSON |
| Reactome | v95 | https://reactome.org/download-data | HTTPS | BioPAX, SBGN, SBML, TSV |
| SIDER | 4.1 | http://sideeffects.embl.de/download/ | HTTPS | TSV |
| SNOMED CT | International 20251201 | https://mlds.ihtsdotools.org/ | Registered download | RF2 (TSV snapshot) |
| STITCH | v5.0 | http://stitch.embl.de/cgi/download.pl | HTTPS | TSV (gz) |
| STRING | v12.0 | https://stringdb-downloads.org/download/ | HTTPS | TSV (gz), FASTA |
| SYMP | v2024-02-09 | http://purl.obolibrary.org/obo/symp.owl | HTTPS | OWL, TAR.GZ |
| TTD | 10.1.01 | https://idrblab.net/ttd/full-data-download | HTTPS | TSV, XLSX, SDF |
| UBERON | v2025-12-04 | http://purl.obolibrary.org/obo/uberon.owl | HTTPS | OBO, OWL, JSON |

Continued on next page

Table S16 – (continued)

| Database | Version | Download URL | Access | Formats |
|---|---|---|---|---|
| UniProtKB | 2025_04 (Swiss-Prot + TrEMBL) | https://ftp.uniprot.org/pub/databases/uniprot/current_release/ | FTP | XML, DAT, FASTA |
| WikiPathways | 20251210 | https://data.wikipathways.org/ | HTTPS | GPML, GMT, SVG, RDF |

**Table S17.** KG entity types, within-source entity counts, and data features by source database. Counts refer to entity objects after deduplication and normalization within each source, rather than distinct entities after cross-source KG integration.

| KG entity type | Source database | Within-source entity count | Data feature |
|---|---|---|---|
| drug | DrugBank | 17,458 | DrugBank ID, name, SMILES, MOA, function |
| | ChEMBL | 16,675 (4,019 with MOA) | ChEMBL ID, name, SMILES, mechanism of action |
| | PubChem | 179,555 | CID, name, SMILES, synonyms |
| | ChEBI | 117,124 | ChEBI ID, name, synonyms, definition, status |
| | MeSH | 10,371 | MeSH ID, name, synonyms, definition |
| | EFO | 54 | EFO ID, name, EXACT synonyms, type-compatible exact cross-references, definition |
| gene | NCBI Gene | 86,599 | Gene ID, symbol, description |
| | HGNC (gene) | 44,921 (+5,286 withdrawn/obsolete) | HGNC ID, symbol, name, locus type, cross-references |
| | HGNC (gene group) | 2,035 (+972 external-resource entries) | Gene-group ID, name, external resource links |
| protein | PRO (Protein Ontology) | 255,372 (+5,570 obsolete) | PRO ID, protein name, synonyms, definition |
| disease | MONDO | 30,373 | MONDO ID, external IDs, name, synonyms, definition, category, clinical features, genetics, treatments, prognosis |
| | Disease Ontology (DO) | 11,997 | DOID, alt_id, external_id, name, synonyms, definition, is_a |
| | Orphanet | 9,787 | ORPHA code, label, disorder type, synonyms, external references |
| | HPO | 12,762 | MONDO ID, database ID, disease name, associated phenotypes |
| | PharmGKB | 1,272 (mapping only) | MONDO ID, name, PharmGKB ID |
| | Mayo Clinic | 553 (ID-matched) | MONDO ID, name, clinical features, diagnostic methods, treatments |
| | Open Targets | 12,628 | Normalized ID, name, description, synonyms |
| | OMIM | 4,298 | Disease, name, description, clinical features, pathogenesis, inheritance, PMIDs |
| | MeSH | 5,131 | MeSH ID, name, synonyms, definition |
| | MarkerDB | 566 | Disease ID, name |

Continued on next page

Table S17 – (continued)

| KG entity type | Source database | Within-source entity count | Data feature |
|---|---|---|---|
| | EFO | 3,779 | EFO ID, name, EXACT synonyms, type-compatible exact cross-references, definition, ontology relations |
| phenotype | HPO | 19,762 | HP ID, name, definition, synonyms, alt IDs, is_a |
| | Open Targets | 2,454 | Normalized ID, name, description, synonyms, parents |
| | EFO | 246 | EFO ID, name, EXACT synonyms, type-compatible exact cross-references, definition, ontology relations |
| anatomy | UBERON | 14,677 | UBERON ID, name, alt_id, synonym, definition |
| | MeSH | 1,776 | MeSH ID, name, synonyms, definition |
| | EFO | 148 | EFO ID, name, EXACT synonyms, type-compatible exact cross-references, definition, ontology relations |
| taxon | NCBI Taxonomy Database | 2,693,994 (including 44 ontology auxiliary records) | Taxon or ontology auxiliary ID, alt_id, name, synonym, rank |
| | MeSH | 3,937 | MeSH ID, name, synonyms, definition |
| | EFO | 450 | EFO ID, name, EXACT synonyms, type-compatible exact cross-references, definition, ontology relations |
| cell | CL (Cell Ontology) | 3,238 | CL ID, name, synonyms, definition |
| | EFO | 1,580 (including 1 obsolete) | EFO ID, name, EXACT synonyms, type-compatible exact cross-references, definition, lifecycle status |
| metabolite | HMDB | 217,960 | HMDB ID, name, synonyms, SMILES, InChI, formula, CAS, status |
| symptom | Symptom Ontology | 894 | SYMP ID, name, synonym, definition |
| pathway | WikiPathways | 850 | Pathway ID, name, pathway IRI, organism, revision, source RDF file |
| | Reactome | 2,934 (including 64 edge-derived supplements) | Reactome ID, name, pathway IRI, organism, available pathway descriptions |
| | HMDB | 49,604 | SMPDB ID, name, pathway IRI, organism; extracted from metabolite–pathway edges |
| | KEGG | 489 | KEGG pathway ID, name, pathway IRI, organism; extracted from disease–pathway edges |
| biological process | Gene Ontology | 30,817 (including 5,664 obsolete) | GO ID, name, synonyms, definition, namespace, lifecycle status |
| | MeSH | 1,625 | MeSH ID, name, synonyms, definition |
| | EFO | 183 | EFO ID, name, EXACT synonyms, type-compatible exact cross-references, definition, ontology relations |
| cellular component | Gene Ontology | 4,574 (including 516 obsolete) | GO ID, name, synonyms, definition, namespace, lifecycle status |

Continued on next page

Table S17 – (continued)

| KG entity type | Source database | Within-source entity count | Data feature |
|---|---|---|---|
| | EFO | 2 | EFO ID, name, EXACT synonyms, type-compatible exact cross-references, definition, ontology relations |
| molecular function | Gene Ontology | 12,805 (including 2,662 obsolete) | GO ID, name, synonyms, definition, namespace, lifecycle status |
| experimental factor | EFO | 1,642 | EFO ID, name, EXACT synonyms, type-compatible exact cross-references, definition, ontology relations |
| measurement | EFO | 7,584 | EFO ID, name, EXACT synonyms, type-compatible exact cross-references, definition, ontology relations |

**Table S18.** Within-source object counts by endpoint group and source database. Counts refer to objects after deduplication and normalization within each source, rather than deduplicated KG triples after cross-source integration. For HumanPPI, mapping 17,849 protein-pair predictions to Ensembl genes produced 167,136 records.

| Endpoint group | Source database | Within-source object count |
|---|---|---|
| Gene-gene | BioGRID | 1,063,441 |
| | STRING | 1,404,260 |
| | IntAct | 10,621 |
| | humanPPI | 17,849 |
| | Human Protein Atlas (HPA) | 23,594 |
| | HGNC (gene-group–gene-group) | 1,259 |
| Gene-protein | UniProt | 21,802 |
| Gene–cellular component | Gene Ontology | 62,056 |
| Gene–biological process | Gene Ontology | 64,708 |
| Gene–molecular function | Gene Ontology | 50,500 |
| Biological process–biological process | Gene Ontology | 55,991 |
| | MeSH | 2,024 |
| Molecular function–molecular function | Gene Ontology | 12,905 |
| Cellular component–cellular component | Gene Ontology | 6,653 |
| Biological process–molecular function | Gene Ontology | 1,319 |
| Biological process–cellular component | Gene Ontology | 142 |
| Cellular component–molecular function | Gene Ontology | 43 |
| Gene-anatomy | Bgee | 5,742,650 |
| Gene-cell | Bgee | 242,886 |

Continued on next page

Table S18 – (continued)

| Endpoint group | Source database | Within-source object count |
|---|---|---|
| Drug-gene | BindingDB | 279,857 |
| | TTD | 32,919 |
| | DrugCentral | 28,664 |
| | DGIdb (PubChem-only IDs) | 22,657 |
| | DGIdb (other IDs) | 61,094 |
| | BioGRID | 10,707 |
| | Open Targets (MOA) | 11,634 |
| | Open Targets (known-drug × disease) | 16,160 |
| | Open Targets (known-drug × HPO) | 5,267 |
| | PharmGKB/ClinPGx | 9,975 |
| | STITCH (filtered) | 152,551 |
| Drug-drug | DrugBank | 1,728,548 |
| | DrugCentral | 7,621 |
| | ChEBI | 275,634 |
| | MeSH | 14,346 |
| Disease-gene | Open Targets | 214,417 |
| | Open Targets (known-drug) | 34,931 |
| | HPO | 16,966 |
| | Orphanet | 7,936 |
| | PharmGKB/ClinPGx | 7,718 |
| | MONDO | 6,109 |
| | OMIM | 3,496 |
| Disease-drug | DrugCentral | 42,348 |
| | Open Targets (traits/EFO) | 29,828 |
| | CTD | 28,427 |
| | Open Targets (known-drug) | 24,581 |
| | Open Targets | 10,929 |
| Drug-*(mechanistic pathway) | DrugMechDB | 32,641 |
| Disease-disease | MeSH | 8,084 |
| | MONDO | 40,642 |
| | Orphanet | 19,296 |
| | Disease Ontology (DO) | 16,888 |
| | Open Targets | 14,778 |
| Disease-phenotype | HPO | 255,709 |
| | Open Targets | 162,346 |
| | Orphanet | 115,627 |
| | OMIM | 89,522 |
| | PhenomeBrowser | 5,314 |
| | MONDO | 1,114 |
| Disease-pathway | CTD (Reactome ID) | 414,261 |
| Disease-anatomy | MONDO | 1,013 |
| Disease-taxon | MONDO | 2,224 |

Continued on next page

Table S18 – (continued)

| Endpoint group | Source database | Within-source object count |
|---|---|---|
| Gene-phenotype | PhenomeBrowser (DeepPheno-derived) | 1,486,298 |
| | PhenomeBrowser (text-mined) | 403,236 |
| | HPO | 366,172 |
| | Open Targets (known-drug) | 6,320 |
| Drug-phenotype | PhenomeBrowser | 175,713 |
| | Open Targets (known-drug) | 3,770 |
| Phenotype-phenotype | Open Targets | 2,966 |
| Metabolite-phenotype | PhenomeBrowser | 27,850 |
| Pathogen-phenotype | PhenomeBrowser | 32,421 |
| Anatomy-anatomy | UBERON | 36,877 |
| | MeSH | 2,277 |
| Anatomy-taxon | UBERON | 2,675 |
| Taxon-taxon | NCBI Taxonomy | 2,693,992 |
| | MeSH | 4,268 |
| Cell-* | CL (Cell Ontology) | 13,805 |
| Metabolite-protein | HMDB | 863,759 |
| Metabolite-pathway | HMDB | 815,803 |
| Metabolite-anatomy | HMDB | 67,548 |
| Metabolite-disease | HMDB | 27,670 |
| Symptom-symptom | Symptom Ontology | 895 |
| Protein-* edge | PRO (Protein Ontology) | 1,042,855 |
| EFO typed entity–typed entity | EFO | 28,932 |

**Table S19.** Predefined relation types, directionality, and definitions used for relation normalization. Direction-invariant relations do not distinguish between the two endpoint orders, whereas direction-sensitive relations distinguish the roles of the head and tail entities.

| Directionality | Relation type | Definition |
|---|---|---|
| **Direction-invariant** *Undirected endpoints* | `ASSOCIATED_WITH` | explicit association, link or correlation when no more specific label applies. |
| | `NOT_ASSOCIATED_WITH` | an explicitly reported absence of association/relationship or a null effect, lack of correlation or equivalent negative evidence; never use it merely because no positive relation is supported. |
| | `SIMILAR_TO` | explicit similarity. |
| | `DIFFERENT_FROM` | explicit difference. |
| | `INTERACTS_WITH` | directly interacts with. |
| | `BINDS_TO` | directly binds to. |
| | `COMBINED_WITH` | used in combination with another intervention or entity. |
| | `COMPARED_WITH` | explicitly compared with. |
| | `RELATIVE_TO` | spatially relative to. |

Continued on next page

Table S19 – (continued)

| Directionality | Relation type | Definition |
|---|---|---|
| **Direction-sensitive** *Directed endpoints* | IS_A | is a type or class of. |
| | HAS_SUBTYPE | has a subtype. |
| | HAS_PART | has a part or component. |
| | CONTAINS | contains or includes. |
| | HAS_ATTRIBUTE | has a characteristic, property, classification label or evidence tag. |
| | MAPS_TO | maps or corresponds to. |
| | ENCAPSULATES | encapsulates or encloses. |
| | LOCATED_IN | located or found in. |
| | RESTRICTED_TO | restricted to a tissue, cell type, region or context. |
| | DERIVED_FROM | is derived from or originates from. |
| | OBSERVED_IN | observed in a tissue, organism, sample, lead, condition or context. |
| | OCCURS_IN | occurs in. |
| | EVALUATED_IN | explicitly evaluated or assessed in or by a setting, population, specimen or modality when MEASURES, DIAGNOSES, DETECTS, or OBSERVED_IN does not fit better. |
| | SPREADS_TO | spreads, invades or metastasizes to. |
| | PARTICIPATES_IN | participates in or is involved in a process. |
| | PERFORMED_VIA | a procedure or action is performed using a method or modality. |
| | UNDERGOES | undergoes a process or transformation. |
| | TRANSFORMS_TO | transforms or converts into. |
| | PRECEDES | temporally or procedurally precedes. |
| | DEPENDS_ON | requires or depends on. |
| | CAUSES | directly causes or results in. |
| | POSITIVELY_REGULATES | activates, induces, promotes or increases. |
| | NEGATIVELY_REGULATES | inhibits, suppresses or decreases. |
| | REGULATES | generic regulation when polarity is unclear. |
| | AFFECTS | exerts a general effect or influence when polarity, causation or treatment is not specified. |
| | CONTRIBUTES_TO | contributes to or helps drive. |
| | MODIFIES | functionally or chemically modifies. |
| | INTERFERES_WITH | interferes with. |
| | SUSTAINS | maintains a disease or pathological state. |
| | TREATS | used to treat or manage a disease or condition. |
| | CONTRAINDICATED_FOR | explicitly should not be used or is advised against for a disease, condition or patient population. |
| | INDICATED_FOR | indicated or used for a disease or condition when an actual treatment effect is not clearly expressed. |
| | PREVENTS | used to prevent a disease or outcome. |
| | IMPROVES | improves a condition, symptom, feature, measurement or outcome. |
| | PREDISPOSES_TO | increases susceptibility or risk. |
| | HAS_PHENOTYPE | has or is characterized by a phenotype or feature. |

Continued on next page

Table S19 – (continued)

| Directionality | Relation type | Definition |
|---|---|---|
| | `MANIFESTS_AS` | manifests or presents as. |
| | `MIMICS` | mimics another phenotype or condition. |
| | `MEASURES` | measures or quantifies a variable. |
| | `DIAGNOSES` | diagnoses a disease or condition. |
| | `DETECTS` | detects a target. |
| | `INDICATES` | indicates a condition or state. |
| | `CONFIRMS` | confirms a diagnosis or finding. |
| | `TARGETS` | directly targets. |
| | `ACTIVE_AGAINST` | is active against a target, pathogen or other entity. |
| | `PRODUCES` | generates or produces a product. |
| | `DELIVERS` | delivers a therapeutic or active entity. |
| | `USED_BY` | is used, utilized or employed by another entity. |
| | `SUBSTRATE_OF` | is a substrate of. |

**Table S20.** Overall entity- and triple-extraction performance on the literature extraction benchmark. Entity matching is constrained to the same benchmark extraction entity category and entity scores are micro-averaged. Triple scores use relation-label-compatible matching. Values are fractions; the best result in each column is bold. A dash indicates that the method does not perform triple extraction. All configurations in the two SPIDER groups used the same agent workflow. The primary configuration used DeepSeek-V4-Flash and was adopted for subsequent experiments; the single-pass LLM baseline used the same backbone without the staged agent workflow.

| | | Entity extraction | | | Triple extraction | | |
|---|---|---|---|---|---|---|---|
| **Method class** | **Method/model** | **Precision** | **Recall** | **F1** | **Precision** | **Recall** | **F1** |
| Conventional baseline | BERN2 | 0.477 | 0.322 | 0.385 | – | – | – |
| | GLiNER-Relex | 0.777 | 0.449 | 0.569 | 0.636 | 0.007 | 0.014 |
| | PubTator | 0.725 | 0.401 | 0.517 | 0.154 | 0.027 | 0.047 |
| SPIDER: alternative backbone | Claude Sonnet 4.6 | 0.923 | 0.702 | 0.797 | 0.703 | 0.503 | 0.586 |
| | DeepSeek V3.2 | 0.863 | 0.809 | 0.835 | 0.647 | 0.540 | 0.589 |
| | Gemini 3.1 Flash-Lite | 0.915 | 0.734 | 0.815 | 0.675 | 0.519 | 0.587 |
| | GPT-5.5 | 0.814 | 0.815 | 0.815 | 0.473 | 0.545 | 0.507 |
| | Grok 4.1 Fast | 0.903 | 0.709 | 0.794 | 0.634 | 0.500 | 0.559 |
| | MiniMax M2.5 | 0.844 | 0.623 | 0.717 | 0.566 | 0.355 | 0.436 |
| Single-pass LLM baseline | DeepSeek-V4-Flash | 0.785 | 0.737 | 0.761 | 0.527 | 0.421 | 0.468 |
| SPIDER: primary configuration | SPIDER (DeepSeek-V4-Flash) | **0.975** | **0.885** | **0.928** | **0.821** | **0.767** | **0.793** |

**Table S21.** Entity-extraction performance stratified by benchmark extraction entity category. Each cell reports precision/recall/F1; the highest F1 per category is bold. Dashes denote an unsupported extraction entity category.

**(a) Biomedical extraction entity categories**

| Method class | Method/model | Anatomy | Chemical | Gene | Organism | Disease |
|---|---|---|---|---|---|---|
| Conventional baseline | BERN2 | 0.248/0.098/0.140 | 0.553/0.500/0.526 | 0.363/0.576/0.445 | 0.087/0.376/0.142 | 0.605/0.639/0.621 |
| | GLiNER-Relex | 0.868/0.458/0.599 | 0.907/0.504/0.648 | 0.873/0.236/0.372 | 0.448/0.661/0.534 | 0.820/0.769/0.794 |
| | PubTator | –/–/– | 0.846/0.669/0.748 | 0.768/0.540/0.634 | 0.158/0.599/0.250 | 0.798/0.838/0.818 |
| SPIDER: alternative backbone | Claude Sonnet 4.6 | 0.892/0.751/0.816 | 0.959/0.835/0.892 | 0.928/0.823/0.872 | 0.911/0.761/0.830 | 0.941/0.893/0.917 |
| | DeepSeek V3.2 | 0.883/0.778/0.827 | 0.959/0.860/0.907 | 0.946/0.800/0.867 | 0.902/0.841/0.870 | 0.938/0.887/0.912 |
| | Gemini 3.1 Flash-Lite | 0.914/0.717/0.804 | 0.952/0.862/0.904 | 0.861/0.862/0.862 | 0.891/0.792/0.838 | 0.961/0.833/0.892 |
| | GPT-5.5 | 0.837/0.778/0.806 | 0.927/0.878/0.901 | 0.905/0.849/0.876 | 0.847/0.838/0.843 | 0.899/0.896/0.898 |
| | Grok 4.1 Fast | 0.848/0.772/0.808 | 0.963/0.832/0.893 | 0.933/0.828/0.877 | 0.914/0.783/0.843 | 0.968/0.800/0.876 |
| | MiniMax M2.5 | 0.864/0.652/0.743 | 0.935/0.698/0.799 | 0.948/0.625/0.753 | 0.792/0.679/0.731 | 0.953/0.657/0.778 |
| Single-pass LLM baseline | DeepSeek-V4-Flash | 0.857/0.683/0.760 | 0.883/0.841/0.861 | 0.928/0.593/0.723 | 0.492/0.841/0.621 | 0.955/0.811/0.877 |
| SPIDER: primary configuration | SPIDER (DeepSeek-V4-Flash) | **0.985/0.830/0.901** | **0.991/0.933/0.961** | **0.986/0.903/0.943** | **0.983/0.957/0.970** | **0.994/0.916/0.953** |

**(b) Clinical and experimental extraction entity categories**

| Method class | Method/model | Process | Procedure | Measurement | Device |
|---|---|---|---|---|---|
| Conventional baseline | BERN2 | –/–/– | –/–/– | –/–/– | –/–/– |
| | GLiNER-Relex | 0.530/0.293/0.377 | 0.595/0.182/0.278 | 0.535/0.104/0.174 | 0.556/0.119/0.196 |
| | PubTator | –/–/– | –/–/– | –/–/– | –/–/– |
| SPIDER: alternative backbone | Claude Sonnet 4.6 | 0.824/0.361/0.502 | 0.930/0.460/0.615 | 0.863/0.461/0.601 | 0.796/0.317/0.454 |
| | DeepSeek V3.2 | 0.765/0.543/0.635 | 0.747/0.794/0.770 | 0.743/0.752/0.748 | 0.678/0.643/0.660 |
| | Gemini 3.1 Flash-Lite | 0.773/0.323/0.456 | 0.900/0.676/0.772 | 0.849/0.578/0.688 | 0.732/0.675/0.702 |
| | GPT-5.5 | 0.596/0.681/0.636 | 0.761/0.756/0.758 | 0.654/0.720/0.685 | 0.742/0.556/0.635 |
| | Grok 4.1 Fast | 0.684/0.375/0.484 | 0.898/0.535/0.670 | 0.815/0.591/0.685 | 0.649/0.627/0.638 |
| | MiniMax M2.5 | 0.570/0.494/0.529 | 0.797/0.568/0.663 | 0.698/0.550/0.615 | 0.670/0.508/0.578 |
| Single-pass LLM baseline | DeepSeek-V4-Flash | 0.541/0.618/0.577 | 0.761/0.673/0.714 | 0.603/0.659/0.630 | 0.327/0.722/0.451 |
| SPIDER: primary configuration | SPIDER (DeepSeek-V4-Flash) | **0.956/0.791/0.866** | **0.959/0.895/0.926** | **0.934/0.824/0.875** | **0.907/0.849/0.877** |

**Table S22.** PMID-level provenance coverage of the research-oriented subgraph before and after literature augmentation. Literature-extracted entities were linked to KG nodes by exact matching against normalized names and aliases; entities without an exact match were assigned to the single highest-scoring node in the subgraph by BioLORD-2023 cosine similarity, provided that the similarity was at least 0.85. $L_0$ denotes the research-oriented subgraph before literature augmentation. $L_0$ provenance denotes the number and percentage of native entities or undirected entity pairs linked to at least one PMID. $L_1$ denotes the literature-augmented research-oriented subgraph, containing the union of native and literature-derived data. $L_1$ provenance denotes the number and percentage linked to at least one PMID in the literature-augmented research-oriented subgraph. $\Delta$ is the absolute improvement in provenance coverage from $L_0$ to $L_1$, measured in percentage points (pp).

| Level | $L_0$ | $L_0$ provenance (%) | $L_1$ | $L_1$ provenance (%) | $\Delta$ (pp) |
|---|---|---|---|---|---|
| Entity | 198,659 | 28,944 (14.57%) | 198,659 | 129,118 (64.99%) | +50.43 |
| Entity pair (undirected) | 1,099,652 | 373,457 (33.96%) | 3,411,323 | 2,707,041 (79.35%) | +45.39 |

**Table S23.** Performance on SPIDER-Bench under different knowledge-input settings. KG-derived inputs were retrieved from SPIDER-KG. Study conditions denote structured conditions, and publication evidence denotes PMID-linked titles and abstracts. All settings were evaluated on the same 200 questions from 118 conflict groups. Macro-$F_1$ was calculated across answer labels A–D. Each metric cell reports the estimate followed by its pointwise percentile 95% confidence interval in brackets, calculated from 10,000 conflict-group bootstrap resamples. The intervals describe individual metric estimates, not differences between settings. Superscripts indicate two-sided paired conflict-group permutation tests against LLM only, using 10,000 permutations. Holm correction was applied jointly across accuracy and macro-$F_1$ for the four non-reference settings (eight tests in one family): $^{\mathrm{ns}}$, $P_{\mathrm{adj}} \geq 0.05$; $^{***}$, $P_{\mathrm{adj}} < 0.001$.

| Knowledge-input setting | Correct | Accuracy (95% CI) | Macro-F1 (95% CI) |
|---|---|---|---|
| LLM only | 101/200 | 0.505 [0.446, 0.564] | 0.503 [0.442, 0.560] |
| Triples | 107/200 | $0.535^{\mathrm{ns}}$ [0.474, 0.596] | $0.532^{\mathrm{ns}}$ [0.468, 0.589] |
| Triples + publication evidence | 156/200 | $0.780^{***}$ [0.721, 0.835] | $0.779^{***}$ [0.719, 0.831] |
| Triples + study conditions | 154/200 | $0.770^{***}$ [0.711, 0.825] | $0.770^{***}$ [0.710, 0.824] |
| Triples + study conditions + publication evidence | 157/200 | $0.785^{***}$ [0.727, 0.838] | $0.784^{***}$ [0.724, 0.835] |

**Table S24.** Performance on SPIDER-Bench across biomedical knowledge graphs. Each KG setting denotes the LLM augmented with the corresponding graph. SPIDER-KG used triples, structured study conditions and PMID-linked titles and abstracts. CROssBARv2-KG and MedKGent-KG used triples with provenance-linked publication titles and abstracts; RTX-KG2, BIOS v3 and PrimeKG used triples only. All settings were evaluated on the same 200 questions from 118 conflict groups. Macro-$F_1$ was calculated across answer labels A–D. Each metric cell reports the estimate followed by its pointwise percentile 95% confidence interval in brackets, calculated from 10,000 conflict-group bootstrap resamples. The intervals describe individual metric estimates, not differences between settings. Both metrics are higher-is-better. Superscripts indicate two-sided paired conflict-group permutation tests against SPIDER-KG, using 10,000 permutations. Holm correction was applied jointly across accuracy and macro-$F_1$ for the five external KGs (ten tests in one family, separate from the information-ablation family): $^{***}$, $P_{\mathrm{adj}} < 0.001$.

| Knowledge graph | Correct | Accuracy (95% CI) | Macro-F1 (95% CI) |
|---|---|---|---|
| SPIDER-KG | 157/200 | 0.785 [0.727, 0.838] | 0.784 [0.724, 0.835] |
| CROssBARv2-KG | 95/200 | $0.475^{***}$ [0.417, 0.533] | $0.472^{***}$ [0.412, 0.527] |
| MedKGent-KG | 85/200 | $0.425^{***}$ [0.362, 0.485] | $0.423^{***}$ [0.359, 0.480] |
| RTX-KG2 | 95/200 | $0.475^{***}$ [0.414, 0.537] | $0.473^{***}$ [0.410, 0.534] |
| BIOS v3 | 100/200 | $0.500^{***}$ [0.441, 0.558] | $0.497^{***}$ [0.434, 0.553] |
| PrimeKG | 100/200 | $0.500^{***}$ [0.439, 0.558] | $0.498^{***}$ [0.434, 0.555] |

**Table S25.** Full-cohort performance across four downstream biomedical datasets. Columns report accuracy (CIViC and BioMixQA), mean reciprocal rank (MRR; Open Targets GWAS), and macro-F1 (DICTrank three- and four-class classification); higher is better. Sample sizes are shown in the headers. Knowledge-graph names denote the LLM augmented with the corresponding graph. Cells give estimates and pointwise 95% percentile confidence intervals from 10,000 task-specific cluster-bootstrap resamples. Methods share the same full cohort within each task. Intervals describe individual estimates, not differences between methods. Dashes indicate methods not included in the CIViC comparison.

| **Method** | **CIViC** Accuracy $n = 1{,}616$ | **BioMixQA** Accuracy $n = 311$ | **Open Targets GWAS** MRR; $n = 138$ | **DICTrank** 3-class classification Macro-F1; $n = 1{,}211$ | **DICTrank** 4-class classification Macro-F1; $n = 1{,}318$ |
|---|---|---|---|---|---|
| LLM only | 0.543 [0.510, 0.575] | 0.875 [0.836, 0.910] | 0.820 [0.744, 0.887] | 0.466 [0.439, 0.493] | 0.348 [0.327, 0.368] |
| CROssBARv2-KG | 0.636 [0.605, 0.667] | 0.910 [0.878, 0.939] | 0.824 [0.750, 0.892] | 0.463 [0.435, 0.490] | 0.377 [0.349, 0.406] |
| MedKGent-KG | 0.602 [0.568, 0.636] | 0.894 [0.859, 0.926] | 0.867 [0.792, 0.930] | 0.470 [0.442, 0.497] | 0.349 [0.326, 0.372] |
| RTX-KG2 | — | 0.894 [0.859, 0.926] | 0.851 [0.774, 0.918] | 0.483 [0.455, 0.510] | 0.362 [0.338, 0.386] |
| BIOS v3 | — | 0.862 [0.820, 0.897] | 0.863 [0.796, 0.922] | 0.481 [0.453, 0.509] | 0.353 [0.330, 0.374] |
| PrimeKG | — | 0.894 [0.859, 0.926] | 0.840 [0.771, 0.903] | 0.468 [0.440, 0.495] | 0.349 [0.325, 0.374] |
| SPIDER-KG | 0.684 [0.652, 0.718] | 0.955 [0.929, 0.977] | 0.863 [0.793, 0.922] | 0.494 [0.466, 0.522] | 0.372 [0.349, 0.396] |

**Table S26.** Primary-metric performance on SPIDER-KG evidence-available subsets, using the same metrics as Table S25 (higher is better). Subsets contain questions for which valid, selected external evidence reached the SPIDER-KG final answerer, irrespective of answer correctness. Selected DOID mappings count as evidence for BioMixQA questions explicitly querying disease identifiers; identifiers used only for entity matching are excluded. Both methods use the same subset and the original full-cohort predictions; sample sizes are shown in the headers. Cells report estimates and pointwise 95% percentile confidence intervals from 10,000 task-specific cluster-bootstrap resamples generated independently of those for Table S25. Holm-adjusted $P$ values compare SPIDER-KG with LLM only using 10,000 two-sided paired cluster permutations. The five primary evaluations form one multiplicity family; secondary metrics are excluded. Superscripts: $^{***}$, $P_{\mathrm{adj}} < 0.001$; $^{**}$, $0.001 \leq P_{\mathrm{adj}} < 0.01$; $^{*}$, $0.01 \leq P_{\mathrm{adj}} < 0.05$; $^{\mathrm{ns}}$, $P_{\mathrm{adj}} \geq 0.05$.

| **Method** | **CIViC** Accuracy $n = 1{,}144$ | **BioMixQA** Accuracy $n = 307$ | **Open Targets GWAS** MRR; $n = 137$ | **DICTrank** 3-class classification Macro-F1; $n = 1{,}192$ | **DICTrank** 4-class classification Macro-F1; $n = 1{,}296$ |
|---|---|---|---|---|---|
| LLM only | 0.558 [0.517, 0.597] | 0.879 [0.840, 0.915] | 0.825 [0.752, 0.894] | 0.464 [0.436, 0.491] | 0.346 [0.326, 0.367] |
| SPIDER-KG | $0.758^{***}$ [0.717, 0.799] | $0.961^{***}$ [0.938, 0.980] | $0.869^{\mathrm{ns}}$ [0.801, 0.929] | $0.493^{\mathrm{ns}}$ [0.465, 0.522] | $0.371^{\mathrm{ns}}$ [0.348, 0.395] |
| Holm-adjusted $P$ value | 0.0005 | 0.0005 | 0.1263 | 0.1263 | 0.1263 |
